# Performance and Robustness of Parameter Estimation from Phylogenetic Trees Using Neural Networks

**DOI:** 10.1101/2024.08.02.606350

**Authors:** Tianjian Qin, Koen J. van Benthem, Luis Valente, Rampal S. Etienne

**Affiliations:** Groningen Institute for Evolutionary Life Sciences, University of Groningen, Nijenborgh 7, Groningen, 9747 AG, The Netherlands; Naturalis Biodiversity Center, Darwinweg 2, Leiden, 2333 CR, The Netherlands

**Keywords:** graph neural network, recurrent neural network, machine learning, regression

## Abstract

Species diversification is characterized by speciation and extinction, the rates of which can, under some assumptions, be estimated from time-calibrated phylogenies. However, maximum likelihood estimation methods (MLE) for inferring rates are limited to simpler models and can show bias, particularly in small phylogenies. Likelihood-free methods to estimate parameters of diversification models using deep learning have started to emerge, but how robust neural network methods are at handling the intricate nature of phylogenetic data remains an open question. Here we present a new ensemble neural network approach to estimate diversification parameters from phylogenetic trees that leverages different classes of neural networks (dense neural network, graph neural network, and long short-term memory recurrent network) and simultaneously learns from graph representations of phylogenies, their branching times and their summary statistics. Our best-performing ensemble neural network (which corrects graph neural network result using a recurrent neural network) can compute estimates faster than MLE and is less affected by tree size. Our analysis suggests that the primary limitation to accurate parameter estimation is the amount of information contained within a phylogeny, as indicated by its size and the strength of effects shaping it. In cases where MLE is unavailable, our neural network method provides a promising alternative for estimating phylogenetic tree parameters. If there are detectable phylogenetic signals present, our approach delivers results that are comparable to MLE but without inherent biases.

## Introduction

Identifying the underlying mechanisms shaping biodiversity is an important goal in the fields of evolutionary biology and ecology. Species diversification processes can often be characterized by speciation and extinction rates, which can be estimated from time-calibrated phylogenies (Nee et al., 1997) as long as the assumed model structure of diversification resembles the true underlying data generation process (Louca and Pennell, 2021). Time-calibrated phylogenies contain branching times and topological relationships between species and offer a complementary source of information to the often incomplete fossil record (Kidwell and Flessa, 1996). The increasing availability of reconstructed phylogenies has empowered many studies seeking explanations for the underlying diversity patterns using modelling approaches (Etienne et al., 2016; Morlon, 2014; Wagner, 2000). One type of models – birth-death models – are often used to estimate speciation, extinction and diversification rates from reconstructed phylogenetic trees (Hey, 1992; Nee, 2001; Nee et al., 1994, 1997).

Likelihood-based approaches, such as maximum likelihood estimation (MLE) and Bayesian inference, can be used to infer not only speciation and extinction rates, but also possibly existing evolutionary and ecological signals, such as diversity-dependence or trait-dependence of rates from branching times and other information sources (Alexander et al., 2016; Etienne et al., 2014; Foote, 1997; Valente et al., 2015). However, MLE approaches are only mathematically tractable for simple diversification models (Janzen et al., 2015; Lambert et al., 2023). In addition, MLE tends to be biased when estimating diversification parameters from phylogenetic trees. The degree of bias depends on the size of the phylogenetic tree (the number of tips): as tree size increases, the estimates become asymptotically more unbiased (Etienne et al., 2016). MLE may also perform worse on complex models with a high number of parameters (Ward, 2008).

An alternative to these likelihood-based approaches for parameter estimation is Approximate Bayesian Computation (ABC), which approximates the posterior distribution of parameters without requiring explicit calculation of a likelihood function. ABC is often seen as a good substitute to MLE when a likelihood function of a model is not available, as long as simulations of the model are fast and tractable (Beaumont, 2010; Beaumont et al., 2002; Janzen et al., 2015). However, studies using ABC for parameter estimation in phylogenetics remain scarce (Bokma, 2010; Kutsukake and Innan, 2013; Rabosky, 2009; Xie et al., 2023). This is partly due to the fact that is is often difficult to identify adequate summary statistics in ABC, which makes the application and development of this potentially powerful approach challenging.

A promising class of tools that may help overcome the limitations of likelihood-based methods and ABC are machine learning approaches, such as neural networks. Neural networks are comprised of layers of nodes, or “neurons”, which process input data and learn to recognize patterns between input and output data from training data (Charu C, 2018). Classic feed-forward neural networks have achieved good results in tasks such as image recognition and natural language processing (Zhu et al., 2018). Another class of neural networks, graph neural networks, are designed specifically for graph-structured data, such as social networks, molecular structures, and ecological interaction networks. They can capture the dependencies and relationships inherent in data types that can be naturally represented as graphs (Kipf and Welling, 2016) and have shown strong performance in various tasks involving graph representation learning (Li et al., 2020; Rampáek et al., 2022; Ying et al., 2018). Phylogenetic trees can also be viewed as graphs, suggesting that graph neural networks have potential applicability in phylogenetics. Recurrent neural networks, another type of neural network, are designed to handle sequential data, such as time series, by maintaining a memory of previous inputs (Sak et al., 2014; Salehinejad et al., 2017). Recurrent neural networks can process inputs of varying lengths and capture time-dependent features, making them particularly well-suited for tasks where the order of data points is crucial, such as learning parameters from branching times when viewed as time-series data.

Owing to the rapid development of both hardware capability and deep learning algorithms, applications of neural networks in phylogenetic analyses have started to emerge (e.g. Moi and Dessimoz (2022), Reiman et al. (2020), Voznica et al. (2022), Lambert et al. (2023), and Lajaaiti et al. (2023)). For instance, phylogenetic deep learning approaches have been shown to provide reliable estimates of parameters in epidemiological, birth-death, and trait-dependent speciation models Lajaaiti et al. 2023; Lambert et al. 2023; Voznica et al. 2022. Despite their potential, employing neural networks for estimating parameters based on the whole phylogenetic tree, especially those associated with diversification, poses significant challenges and requires further systematic research regarding their performance, accuracy and robustness. Specifically, feed-forward linear neural networks usually require a large amount of data to be able to generalize well on the patterns within the data (Zhu et al., 2018); producing graph representations for graph-level learning can be challenging given the need to aggregate information across diverse graph sizes and topologies (Ying et al., 2018); the capability of the recurrent neural networks to predict parameters from whole sequences is often challenging (Sak et al., 2014). Hence, how robust neural network methods are at handling the intricate nature of phylogenetic data remains an open question.

In this study, we explore the capabilities of neural networks in research on species diversification using phylogenies. We first develop various neural network architectures and protocols for transforming phylogenetic trees and branching times into formats compatible with popular neural network frameworks. We then investigate predictive performance on simulated data for ensemble learning strategies, which combine different neural network classes to maximize data utilization and enhance performance. We also assess the determinants of estimation accuracy and robustness for both neural network and MLE methods under various diversification scenarios. Finally, we implement our trained neural networks on empirical phylogenetic datasets and compare their estimations to those of MLE.

Our analyses encompass three different diversification scenarios for which likelihood-based inference approaches already exist: a constant-rate birth-death (BD) scenario, with constant speciation and extinction rates over time (Stadler, 2011); a diversity-dependent diversification (DDD) scenario, where the number of species in a clade negatively affects the speciation rate (Etienne et al., 2012); and a protracted birth-death (PBD) scenario, where speciation takes time and does not always proceed to completion (Rosindell et al., 2010). Applying our new methodology to phylogenetic trees simulated under a broad range of the parameter space, our findings indicate that neural network approaches are as effective, if not more so, than MLE in recovering parameters from phylogenetic data simulated under these stochastic processes. Trained neural networks can be conveniently applied to empirical trees for parameter estimation. To facilitate this, we present a new R package, “EvoNN,” capable of performing such analyses based on phylogenetic trees (empirical or simulated) supplied by the user (Qin, 2024).

## Materials and Methods

### Software Environment and Computational Budget

We used a hybrid programming environment with PyTorch 1.12.1 (Imambi et al., 2021), PyTorch Geometric 2.3.1 (Fey and Lenssen, 2019), Python 3.7.1 (Python, 2021), CUDA 12.2.2 (Luebke, 2008) and R 4.2.1 (R Core Team, 2013). The procedures of simulation, data transformation, and maximum likelihood estimation were handled through parallel CPU computations on the Hábrók high-performance computing cluster of the University of Groningen. The total computational budget for these processes was approximately 3000 hours (used CPU time). Our neural networks were trained, optimized and evaluated on the NVIDIA A100 and V100 tensor core GPUs of the Hábrók cluster. The estimated computational budget was 1500 hours (used GPU time, excluding CPU time for dataset loading and saving). We implemented a user-friendly tool to estimate parameters from phylogenetic trees using the neural network approach developed in this study in the new R package “evoNN”.

### Simulation Approaches

To train the neural networks, we simulated phylogenetic trees using different functions from different R packages. For each simulated dataset we kept trees with only extant lineages, mimicking reconstructed phylogenies. The settings for the parameters used to simulate the trees were selected to limit the maximum total number of nodes (including root, internal and tip nodes, here and after, we always refer to the total number of nodes) for the trees in each dataset. After simulation, we further filtered out all trees containing more than 3000 nodes to avoid the creation of excessively large matrices that could deplete the available memory space allocated to the GPUs during the GNN training process. Such trees are uncommon under the settings we used – typically fewer than 5 trees with more than 3000 nodes (~1500 tips) are present within each set of phylogenies we acquired from simulation. We also filtered out all trees containing less than 5 nodes (3 tips) to ensure successful data transformation and summary statistic computation. Small trees inherently carry limited informational content. The exclusion of these trees is unlikely to impact performance of the neural networks on the remaining trees (typically fewer than 100 trees with less than 5 nodes were present for each parameter setting).

To consider different diversification processes, we simulated 100,000 random birth-death trees (BD phylogenies), 100,000 diversity-dependent trees (DDD phylogenies) and 100,000 protracted birth-death trees (PBD phylogenies). The amount of simulated data is bounded by the resource and time limits of the computing cluster. All trees have an identical crown age of 10 time units (*t* = 10) to reduce the dimension of data complexity. This age was chosen arbitrarily, as we can always rescale the trees in time. For simulating BD trees, we used the “rlineage” function from R package “ape” (Paradis and Schliep, 2019) to generate complete trees and then pruned all the extinct lineages; for DDD trees, we used the “dd_sim” function from R package “DDD” (Etienne et al., 2012); for PBD trees we used the “pbd_sim” function from our R package “eveGNN” (a codebase of phylogeny simulation, data transformation, neural network training and MLE computation for our study), which is similar to the function with the same name in the original R package “PBD” (Etienne and Rosindell, 2012), but only outputs necessary data for our study.

In our simulation approach we randomly sampled the (log) parameters required for each scenario (BD, DDD and PBD) from uniform distributions. The upper bound for the extinction rates were proportionally dependent on the drawn speciation rate to avoid cases where extinction rates could be larger than speciation rates, because in such cases the whole tree likely goes extinct. Furthermore, to prevent a huge number of evolutionary events that would deplete available computational time and memory, we also imposed an overall cap of 1.5 on the extinction rates. See Table 1 for the detailed parameter distribution settings used in the simulations.

**Table 1.** Parameter settings for the simulated tree datasets. The type column specifies which function is used to generate the trees. The columns specify the crown age (age), the number of trees in the data set (*N*), the lower (*a*) and the upper (*b*) bounds of the parameters for the tree simulations, all the parameters being sampled from *U* (*a, b*), except for *λ*_1_ of the protracted birth-death scenario. *λ*_1_ is computed as *λ*_1_ = 10*^i^* where *i* is sampled from *U* (*−*3, 1). *U* denotes uniform distribution. Sub-table A shows the parameter distributions of the constant-rate birth-death model and the diversity-dependent-diversification model, *λ*: intrinsic speciation rate/birth rate; *µ*: intrinsic extinction rate/death rate; *K*: carrying capacity. Sub-table B shows the parameter distributions of the protracted birth-death model, *λ*_1_: speciation-initiation rate of good species; *λ*_2_: speciation-completion rate; *λ*_3_: speciation-initiation rate of incipient species; *µ*_1_: extinction rate of good species; *µ*_2_: extinction rate of incipient species. *^∗^*In diversity-dependent-diversification simulations, the maximum extinction rate is capped at 1.5 if 0.9*λ >* 1.5.

| Type | Age | N | $\lambda_0$ | | $\mu_0$ | | $K$ | |
| --- | --- | --- | --- | --- | --- | --- | --- | --- |
| | | | $a$ | $b$ | $a$ | $b$ | $a$ | $b$ |
| BD | 10 | 100,000 | 0.1 | 0.8 | 0.0 | $0.9\lambda_0$ | - | - |
| DDD | 10 | 100,000 | 0.1 | 4.0 | 0.0 | $0.9\lambda_0^*$ | 10 | 1000 |

**B:** Parameter settings for PBD trees
| Type | Age | N | $\lambda_1$ | | $\log_{10}(\lambda_2)$ | | $\lambda_3$ | | $\mu_1$ | | $\mu_2$ | |
| --- | --- | --- | --- | --- | --- | --- | --- | --- | --- | --- | --- | --- |
| | | | $a$ | $b$ | $a$ | $b$ | $a$ | $b$ | $a$ | $b$ | $a$ | $b$ |
| PBD | 10 | 100,000 | 0.1 | 1.0 | -3 | 1 | 0.1 | 1.0 | 0.0 | $0.8\lambda_1$ | 0.0 | $0.8\lambda_3$ |

### Data Preparation

We employed three different basic neural network architectures: a dense neural network (DNN), a graph neural network (GNN), and a long short-term memory (LSTM) recurrent network, as illustrated in Figure 1 (see Appendix C for a detailed description). Each of these architectures was refined through validation and required different input data. For the DNN, the input data consisted of a total of 54 summary statistics (Appendix N) for each simulated tree. In the GNN, the full phylogeny was interpreted as a graph and could in that form be used as input data (as illustrated in Figure 2). In the LSTM, we treated branching times of the phylogenies as sequential or time-series data (Sak et al., 2014). Given its recurrent architecture, LSTM is adept at sequence prediction tasks, making it particularly suitable for estimating tree parameters from entire sequences of branching times.

**Fig. 1.**
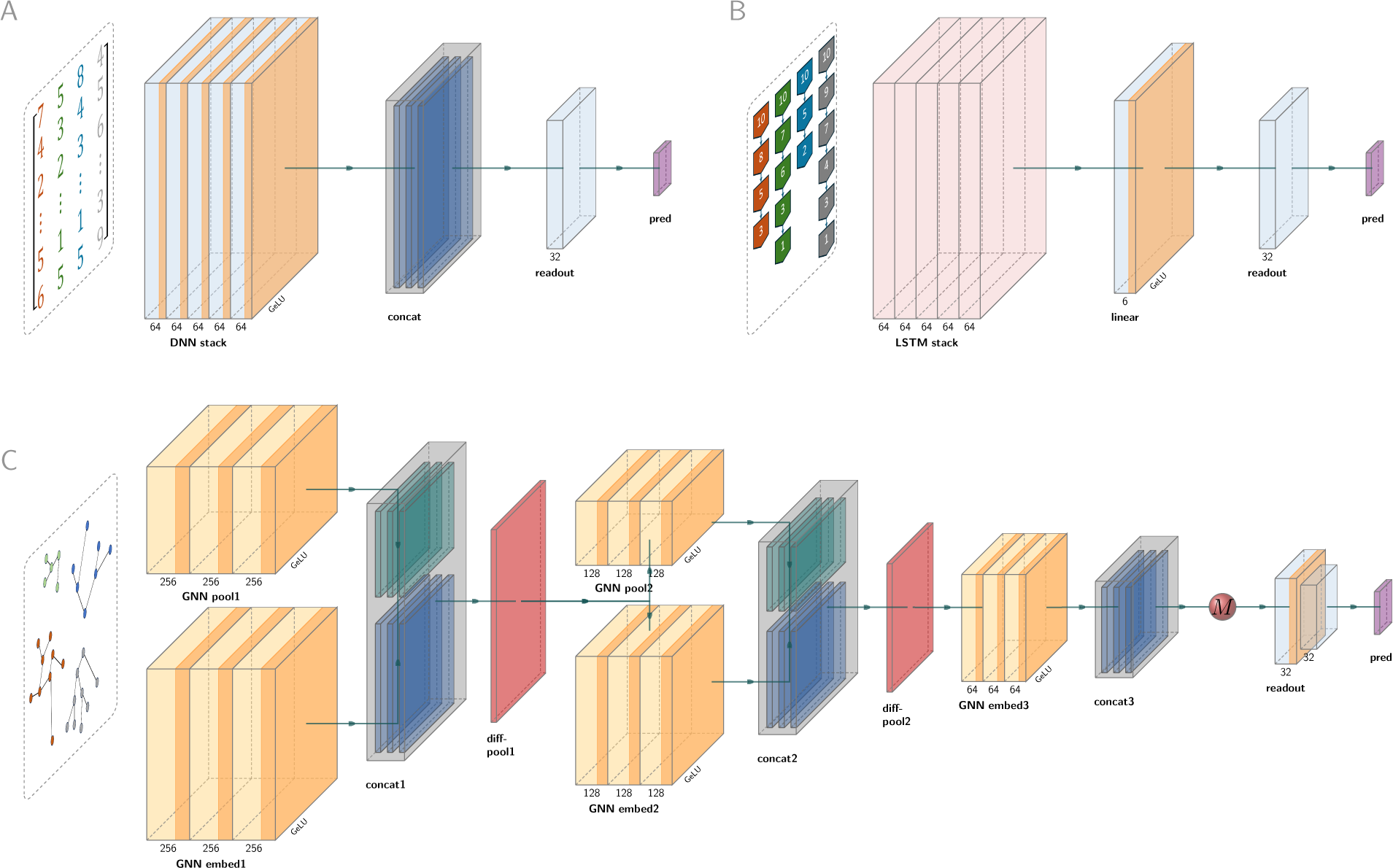
Illustration of the neural network architectures. From left to right, for each neural network, the inputs are filtered through the layers, and the network ultimately outputs the final predictions of the parameters through the readout layers. A: dense neural network (DNN), whose input data are summary statistics. The major component of the DNN is a stack comprising five linear layers (“DNN stack”), each followed by a Batch Normalization for 1D Inputs operator (BatchNorm1D, not shown in figure) and Gaussian Error Linear Units (GELU, the orange band within the boxes). Learned features from all the linear layers within the stacks are collected and concatenated (“concat”). A single linear readout layer (“readout”) outputs *n* predicted parameters (“pred”). B: long short-term memory recurrent neural network (LSTM), whose input data are the branching times. The major component of the LSTM is a stack of five LSTM recurrent neural network layers (“LSTM stack”). Learned features are processed by a linear layer accompanied by a GELU (“linear”), then passed to a single linear readout layer (“readout”) that outputs *n* predicted parameters (“pred”). C: graph neural network (GNN), whose input data is a graph representation of the phylogeny. GNN is assembled from five modules. Each module comprises the same number of GraphSAGE (sample-and-aggregate graph convolutional neural network) operators. Each operator is accompanied by a BatchNorm1d (not shown in the figure) operator and then a GELU activation function (illustrated by the orange bands within the yellow boxes). Learned features from all the GraphSAGE operators within a module are collected and concatenated. The differentiable pooling (DiffPool) technique is adopted to perform graph coarsening. In the first coarsening operation, the graph data inputs are passed to two GNN modules (“GNN pool1” and “GNN embed1”). The pooling group reduces the graph size, while the embedding group captures the node features. The filtered data from each GraphSAGE operator are concatenated (“concat1”) and then passed to a DiffPool layer (“diff-pool1”), which finalizes the first coarsening operation. The second coarsening operation is applied in the same way as the first (as represented by “GNN pool1”, “GNN embed2”, “concat2”), and the outputs from the second DiffPool layer (“diff-pool2”) are passed to the final (fifth) GNN module (“GNN embed3”). After the final GNN module, the outputs are concatenated (“concat3”) and transformed by a global mean pooling operation (red ball “M”) to create a final graph representation. This graph representation is passed to a readout layer group (“readout” as represented by light blue boxes) consisting of two linear layers to perform graph-level regression which ultimately outputs a vector of *n* predicted parameters (“pred” as represented by a purple box). Only the first linear layer is followed by GELU (see the orange band of the first linear layer). See Appendix C for the detailed description and technical details.

**Fig. 2.**
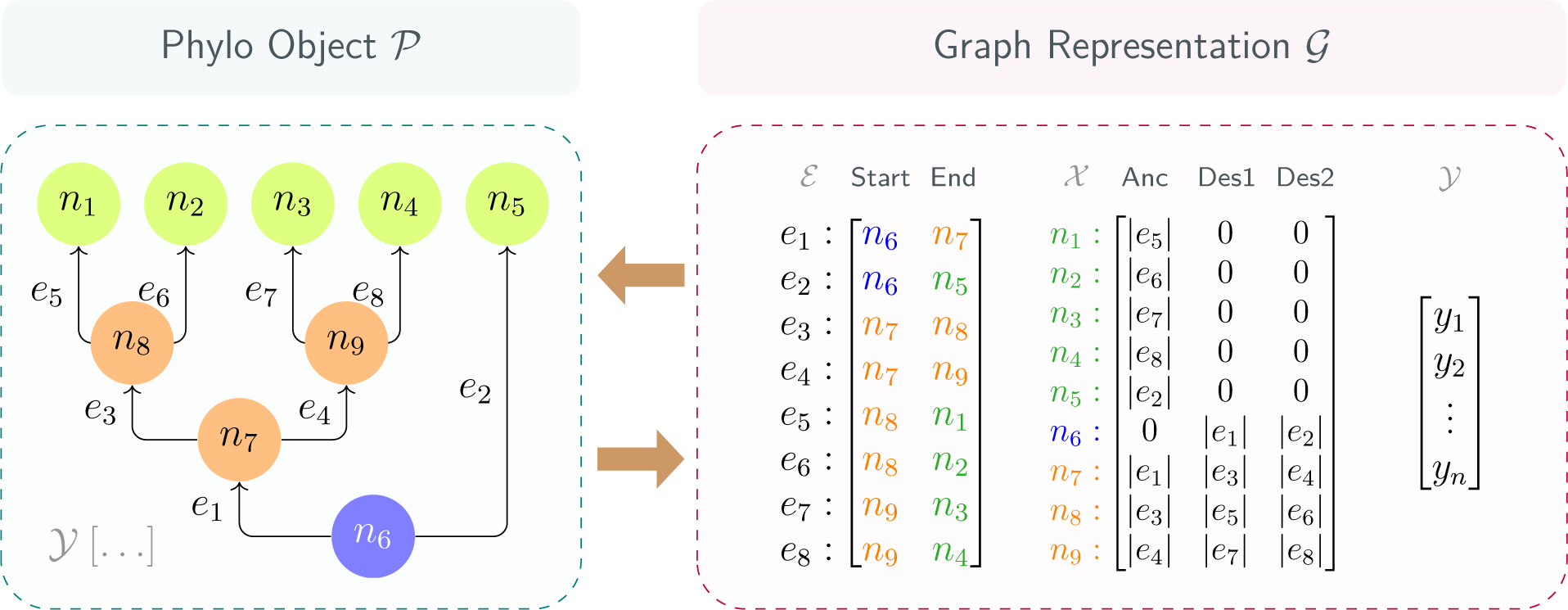
Illustration of data transformation between a “phylo” object and its graph representation. The left panel shows a visualization of a “phylo” object. The blue circle represents the root node, orange circles represent the internal nodes and green circles represent the tip nodes. Arrows represent directed edges between each pair of the nodes. The right panel shows the transformed graph data structure. The adjacency list is denoted as *E*. Each row of the adjacency list represents one edge, the first column represents the starting node and the second column represents the end node. Note that the adjacency list is transposed (in the example into *E* ^2^*^×^*^8^) after converting to a tensor. The node feature matrix is denoted as *X*. Each row of the node feature matrix represents the features contained in one node, the first column represents the distance from the node to its direct ancestor node, the second and the third columns represent the distances from the node to its two descendants. In the node feature matrix, the distances from a node to non-existing nodes (e.g. the tip nodes have no descendants, and the root node has no ancestor) are represented by zeros. The node and edge labels before the colons (including the colons) are placed here for visual assistance. After transformation, we use graph-level attributes *Y* to store the parameters used to generate the “phylo” object. The node labels are given by *n*_1_*, n*_2_*, n*_3_*, …, n*_9_, the edge labels are given by *e*_1_*, e*_2_*, e*_3_*, …, e*_8_, the edge lengths are given by *|e*_1_*|, |e*_2_*|, |e*_3_*|, …, |e*_8_|. The generating parameters are given by a vector [*y*_1_*, y*_2_*, …, y_n_*] where *n* is the number of parameters

Therefore, our data compound comprises three major components: the phylogenetic trees, their corresponding summary statistics, and their branching times, to maximize the use of available data. The functions needed for the data transformations are either available in PyTorch or implemented in our package eveGNN and described in more detail in Appendix A.

### Ensemble Learning Strategies

To leverage all available data and improve prediction accuracy, we combined GNN, DNN, and LSTM using bagging, stacking, and boosting, which are typical ensemble learning strategies (Graczyk et al., 2010). With bagging, we trained GNN, DNN and LSTM independently on the same dataset, translated their original outputs to parameter predictions (we will use “readout” hereafter to refer to this translation) and then aggregated the predictions. We used four aggregation methods: mean, median, max and min.

With stacking, we use GNN, DNN and LSTM in the same architecture but without their own readout layers. Instead, we combined the features learned from DNN, LSTM and GNN and fed them to a meta-learner comprising linear neural network layers that learns the best readout parameter predictions from these combined features. GNN, DNN, LSTM and the meta-learner were trained simultaneously.

With boosting, the neural networks were trained sequentially. Boosting strategies offered various pathways for enhancing model performance. We started with a GNN to make initial predictions and explored the effectiveness of both DNN and LSTM for correcting residuals, either individually or in sequence. We used “Boost SS” to refer to correcting GNN’s residuals by DNN (from summary statistics)“Boost BT” to refer to correcting GNN’s residuals by LSTM (from branching times); “Boost SS+BT” to refer to correcting GNN’s residuals by DNN and then correcting DNN’s residuals of residuals by LSTM; “Boost BT” to refer to correcting GNN’s residuals by LSTM (from branching times); “Boost BT+SS” to refer to correcting GNN’s residuals by LSTM and then correcting LSTM’s residuals of residuals by DNN.

See Figure 3 for a simplified illustration of the ensemble learning strategies.

**Fig. 3.**
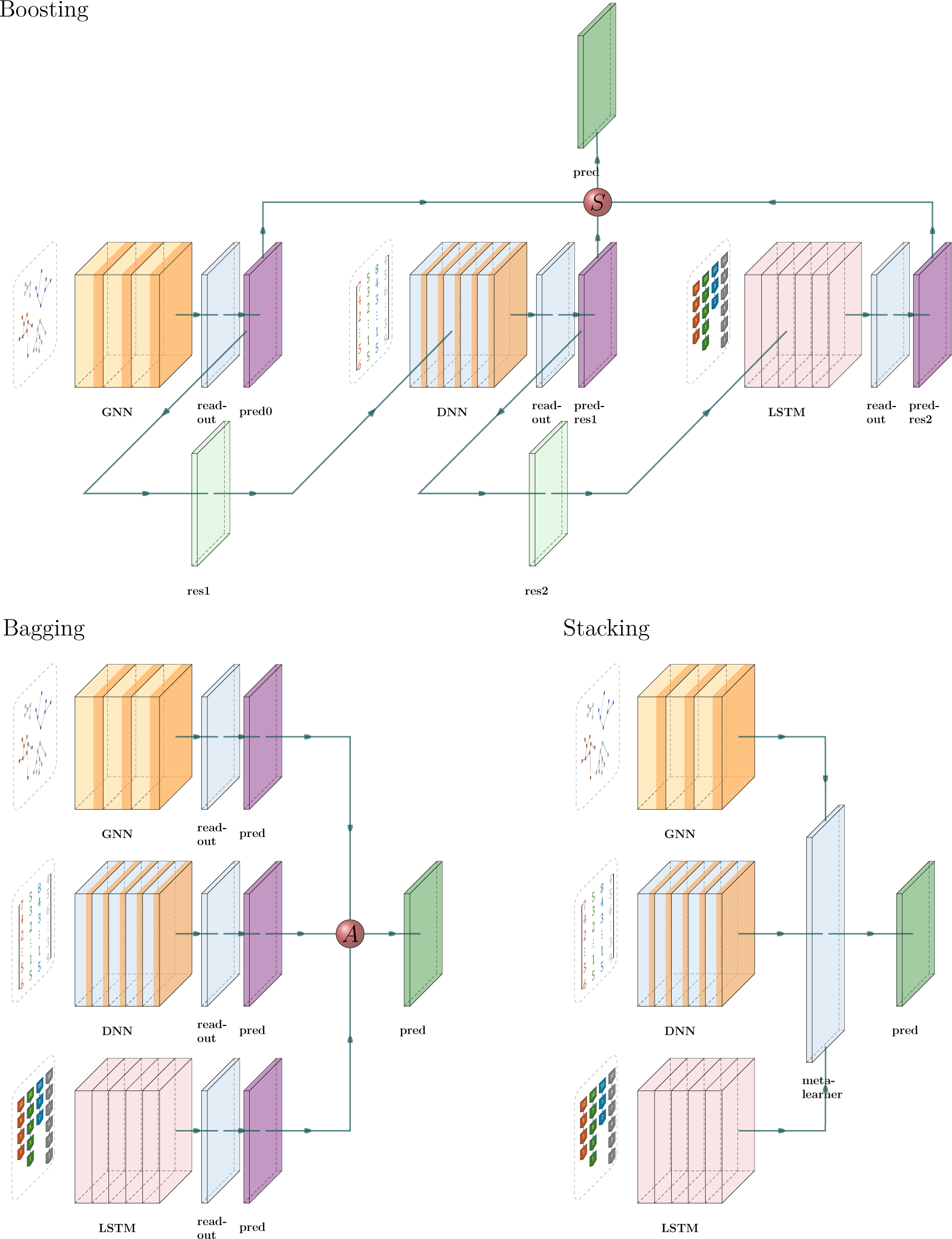
Illustration of the ensemble learning strategies to combine the graph neural network (GNN), the dense neural network (DNN) and the long short-term memory recurrent neural network (LSTM). The neural networks are largely simplified. With boosting, GNN, DNN and LSTM were trained sequentially to iteratively correct residuals. For example, DNN is trained to predict the residuals of GNN predictions. Subsequently, LSTM is trained to predict the residuals of the residuals after DNN corrected the GNN predictions. The final prediction comes from the initial prediction by GNN minus two learned residual terms by DNN and LSTM. With bagging, we trained GNN, DNN and LSTM independently, translated their original outputs to parameter predictions and then aggregated the predictions. With stacking, we trained GNN, DNN and LSTM simultaneously but without readout. We directly concatenated the outputs from GNN, DNN and LSTM and then used a meta-learner to make predictions from the outputs. With bagging, we trained GNN, DNN and LSTM independently (“GNN”, “DNN” and “LSTM” blocks of boxes), translated their outputs to parameter predictions through their own readout layers (three “readout” boxes next to the neural networks and three “pred” boxes next to the readout layers) and then aggregated the predictions (red ball “A”). With stacking, we trained GNN, DNN and LSTM simultaneously (“GNN”, “DNN” and “LSTM” blocks of boxes) but without their own readout layers. We combined the features from DNN, LSTM and GNN and fed to a meta-learner (“meta-learner”) comprising linear neural network layers to output parameter predictions. With boosting, there can be different pathways. In our illustration, GNN, DNN and LSTM were trained sequentially to iteratively correct residuals. First, the GNN is trained from the graphs to make the initial predictions (see “GNN”, “readout” and then “pred0”) and from predicted and ground truth values of the parameters we computed the residuals (“res1”)second, the DNN is trained to predict these residuals from the summary statistics (see “DNN”, “readout” and then “pred-res1”), learning to correct the GNN’s errorslastly, the LSTM is trained to predict the residuals of the residuals (see “LSTM”, “readout” and then “pred-res2”), which is the initial predictions minus the predicted residuals by the DNN, from branching times, to further improve the predictive accuracy. Finally, we subtracted the two residual terms from the initial predictions (red ball “S”) to make the corrected predictions. See Appendix D for a detailed explanation.

### Training Neural Networks

Prior to training, each dataset of 100,000 trees was randomly shuffled and subsequently divided into two segments. The first segment, consisting of 90% of the dataset, was allocated for training purposes, while the remaining 10% was used as the validation dataset for monitoring and fine-tuning the neural network performance. The training session is carried out by epochs, each consisting of three major steps: first, performing forward pass on the training dataset; second, assessing the prediction accuracy; and lastly, performing back-propagation (adjusting the weights of the neuron connections to improve the neural network performance). Back-propagation, requires quantifying the error between the neural networks predictions and the actual ground truth values. We quantified the ‘total loss’ as the sum of the residual error and other terms for facilitating neural network training. We represented total loss using a loss function which sums up all the loss terms (see Appendix B for more detail).

We use the AdamW (Adaptive Moment Estimation with decoupled weight decay) optimizer (Loshchilov and Hutter, 2017) to iteratively update the neural networks’ parameters to minimize the loss function. We used default AdamW argument settings.

During training, we adopted mini-batches of size 64 (data of 64 simulated trees per mini-batch) to reduce GPU memory usage. The total number of epochs was manually optimized to avoid underfitting and overfitting. This was done by comparing the loss metrics for the training dataset to those of the validation datasets at every epoch.

Overfitting is indicated by a training loss that continues to decrease while the validation loss starts to increase, whereas underfitting is suggested by both training and validation losses being high and decreasing at a similar rate. Analyzing these loss trends over time can help to optimize hyper-parameters (“settings” that might alter neural network behavior or impact performance).

Under the BD scenario, the neural networks were trained to predict two parameters: birth rate (*λ*) and death rate (*µ*). Under the DDD scenario, the neural networks were trained to predict three parameters: speciation rate (*λ*), extinction rate (*µ*) and carrying capacity (*K*). Under the PBD scenarios, the neural networks were trained to predict five parameters: speciation rate of the good species (*λ*_1_), speciation completion rate (*λ*_2_), speciation rate of the incipient species (*λ*_3_), extinction rate of the good species (*µ*_1_) and extinction rate of the incipient species (*µ*_2_).

### Baseline Benchmark

Maximum likelihood estimation (MLE) approaches have been developed for the BD, DDD and PBD scenarios (Etienne et al., 2012, 2014; Etienne and Rosindell, 2012). Per scenario, we simulated additional testing datasets each comprising 10,000 phylogenies using the same parameter spaces as the training datasets. For these testing datasets, we adopted the MLE approaches to estimate the parameters of each phylogeny from their branching times under different scenarios. For the BD trees, we estimated their birth and death rates. For the DDD trees, we estimated their speciation rate, extinction rate and carrying capacity. There is a limitation in the MLE approach for PBD trees because to allow for a computation of the likelihood, the speciation initiation rates of good species and incipient species need to be equal (Etienne et al., 2014). We therefore only estimated four initial parameters: speciation initiation rate (for both good and incipient species, assuming they are the same), speciation-completion rate, extinction rate of good species and extinction rate of incipient species, although in our simulation we have five independently sampled parameters. The MLE results are used as a baseline benchmark to evaluate the performance of the neural networks on tree parameter estimation.

We estimated two types of benchmarks using the MLE approaches: one with ground truth parameter values set as the starting point of the MLE searching process, the other with a starting point randomly sampled from the parameter space of the simulation. We consider the first type of benchmarks as a best-case MLE performance (as in real applications ground truth parameters are not known) and the other type as a typical-case MLE performance, which mimics the pragmatic approach if true parameter values are not known. Note that in practice it is possible to achieve better performance than the typical-case, e.g. by optimizing from several starting points.

We also explored the effectiveness of different optimization approaches for MLE on the DDD phylogenies. We used the “Simplex” (Lagarias et al., 1998) optimizer to compute the baseline benchmarks for all the analyses. See Appendix E for reasons and for a detailed comparison between the optimizers.

### Performance Analysis

From the same testing datasets used for baseline benchmark computation, we analysed the patterns of residuals (differences between ground truth and predicted values, which can be viewed as the goodness of fit) by visually examining their relation to true values and the total node counts of the phylogenetic trees, which include root, internal and tip nodes. Considering the complex nature of residual patterns, which may vary according to specific characteristics of the simulation processes (for instance, carrying capacity effects in DDD and protracted speciation in PBD), as well as the performance and robustness of the estimation methods, we calculated error metrics locally for three different phylogeny size ranges, as a global metric could be misleading.

The main case study we decided to focus on is based on simulated trees under the DDD scenario, because it involves more evolutionary mechanisms than the simple birth-death scenario while containing fewer parameters than the protracted birth-death scenario. This simplifies our analyses on the neural network performance while maintaining enough complexity to challenge the capability of our proposed methods. From this case study we identified and selected the most effective MLE optimization algorithm, neural network architecture and ensemble strategy, which we then applied to BD and PBD scenarios. We therefore only analysed the best-performing neural network methods against the typical and best MLE cases on BD and PBD. Additionally, for the PBD scenario, we computed a composite parameter called the mean duration of speciation from the speciation completion rate, the speciation rate of incipient species and the extinction rate of incipient species (Etienne et al., 2014), because MLE can arguably better estimate the mean duration of speciation than the original parameters.

### Robustness Analysis

We assessed the robustness of the estimation results of both neural networks and MLE by measuring the consistency with which these approaches produce similar estimates for phylogenies generated under identical parameter settings. In the previous simulations, each parameter combination was sampled and used only once, whereas in the robustness analysis we repeatedly use identical parameter settings to generate sets of phylogenies (bootstrapping) under the DDD scenario. Even when the same parameters are used, the resulting phylogenies can vary substantially in size, topology and structure due to stochasticity. Such an evaluation helps assess the neural networks’ ability to abstract the underlying parameter influences from the phylogenetic data, regardless of heterogeneity. For each parameter combination, 1000 trees were generated randomly. We used a total of 80 sets of parameter combinations, thus 80,000 phylogenies in total. Specifically, we used all combinations of speciation rates *λ* = 1.0, 1.5, 2.0, 2.5, 3.0, extinction rates *µ* = 0.2, 0.4, 0.6, 0.8 and carrying capacities *K* = 200, 400, 600, 800.

MLE is computationally more expensive than predicting from already trained neural networks, and computational time rapidly increases with the size of the phylogenies. We thus performed MLE on only 2000 simulated phylogenies. To ensure fair visual and numerical comparisons when plotting the results of these analyses, extreme MLE estimates were not shown in the figures (they exceeded the fixed range of the y-axis) and excluded from the computation of the mean absolute errors of the MLE estimates. Neural network results were randomly sub-sampled to match the MLE data count, maintaining equivalent visual density and facilitating a more accurate performance comparison between approaches. For the neural networks, the mean absolute errors were computed on the complete dataset without sub-sampling and exclusion. In the figure, on average (we simulated the testing datasets many times throughout the study), out of 2000 samples, 5-150 samples per MLE figure panel, and 1-3 per neural network figure panel fell beyond the axis range. For the most underperforming method (Boost BT+SS) 600-1500 samples fell beyond the axis range.

We did not analyze the robustness of BD and PBD scenarios, because BD is a special case of DDD (if we set carrying capacity to an infinite value) and PBD related parameters can hardly be estimated accurately using MLE methods (Etienne et al., 2014).

The complete code base for this study, including simulations, data processing, neural network training, evaluation, and both data analysis and visualization tools, is available in the GitHub repository eveGNN (Qin, 2023).

### Empirical Tree Estimation

We deployed pre-trained neural networks to estimate phylogenetic parameters from a dataset of 199 empirical phylogenetic trees curated by Condamine et al. (2019), with a tip count ranging from 20 to 1500. To align with the training conditions of our neural networks, which were trained on simulated phylogenies spanning exactly 10 time units (Myr), we rescaled the crown ages of all empirical trees to this duration. The parameter estimations we present are therefore rescaled. All the selected empirical trees are reconstructed phylogenies and fully bifurcated (each root or internal node has exactly two descendants). If an empirical tree fails an ultrametric (all tip-ends are aligned at the present) test due to branch length precision issue, we forced all its tips to end exactly at the present by extending the shorter tips to align with the longest one. See Appendix M for meta information of the empirical trees.

We used two distinct neural networks, each pre-trained on simulated trees from one of two evolutionary scenarios (BD or DDD) to estimate parameters from the empirical trees. For the BD scenario, we estimated the parameters *λ* (speciation rate) and *µ* (extinction rate); for the DDD scenario, we estimated *λ*, *µ*, and *K* (carrying capacity). We did not estimate parameters for the PBD scenarios because neither neural networks, nor MLE approaches could recover the parameters accurately from the simulated phylogenies. In addition to our neural network estimates, we used MLE methods for parameter estimation to provide a comparative assessment of the results. The MLE methods were set to use default starting points of likelihood optimization, as we do not know the true parameters of the empirical phylogenies.

We used the same bootstrapping method described before to quantify the uncertainties of both MLE and neural network estimates from empirical data. The process involves three main steps: first, estimating parameters from empirical phylogenies using MLE and pre-trained neural networks; second, simulating a set of phylogenies under a specified diversification scenario (such as BD, DDD, or PBD) using the MLE and neural network estimates; and third, re-estimating parameters from the simulated phylogenies using MLE and neural networks. The estimates derived from the bootstrapped phylogenies form a distribution.

We applied this uncertainty computation to a selected set of empirical phylogenies from the Condamine dataset (Condamine et al., 2019) under the DDD scenario (see Appendix F for details). The criteria for selection were phylogenies with more than 300 and less than 1000 nodes, and maximum likelihood estimates (MLE) of *K* (carrying capacity) being less than 1000. The distributions of MLE and neural network estimates from the bootstrapped phylogenies was compared to the original MLE and neural network estimates from empirical phylogenies. For each set of parameters estimated from empirical phylogenies, we bootstrapped 1000 simulated phylogenies.

Our R package “EvoNN” (Qin, 2024) provides functions to perform the uncertainty (bootstrap) analyses.

## Results

### Performance Analysis

We evaluated the performances of various neural networks, both individually and in combination through ensemble strategies, in predicting parameters from simulated DDD phylogenies. These predictions were benchmarked against best-case and typical-case MLE results using the Simplex optimizer.

Among all the methods, we consider boosting GNN with LSTM as the most robust method based on the goodness of fit (see Figure 4, Figure 5, Figure 6 and Figure 7), the mean absolute errors (see Appendix G, Figure 14) and robustness (see rows named Boost BT in Figure 8 and Figure 15). Both neural networks and MLE approaches generally struggle with small phylogenies (see Figure 4, Figure 5 and Figure 6 for larger errors represented by the yellow data points, see also Appendix G, Figure 14). Performance improves significantly on medium and large phylogenies for both neural network and MLE approaches.

**Fig. 4.**
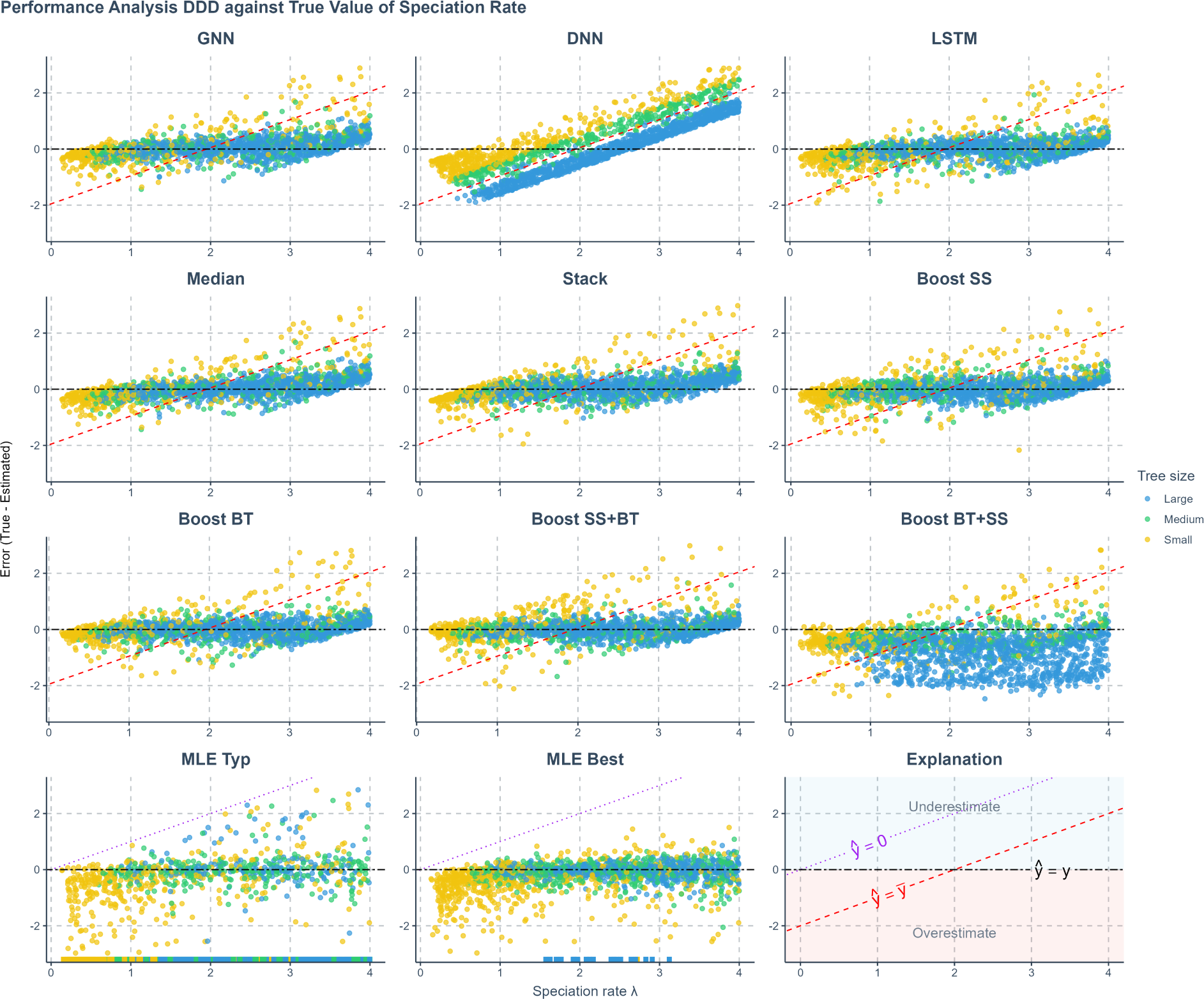
Prediction error of various methods applied to phylogenies simulated under a diversity-dependent diversification scenario, against true values of the speciation rate. The errors shown (y-axis) are the differences between the true parameters (x-axis) used to simulate the phylogenies and the values predicted or estimated by each method. Each panel represents an estimation method. Phylogenies are categorized based on their size: yellow for small phylogenies with fewer than 200 nodes (including root, internal, and tip nodes), green for medium-sized phylogenies with 200 to 500 nodes, and blue for large phylogenies with more than 500 nodes, refer to Appendix G, Figure 17 for how the tree sizes are distributed. GNN: Predictions obtained by the graph neural network using the phylogenies. DNN: Predictions by the dense neural network using summary statistics. LSTM: Predictions by the long short-term memory recurrent neural network using branching times. Median: Bagging strategy that takes the median value of the predictions from GNN, DNN, and LSTM. Stack: Stacking strategy that utilizes a meta-learner to integrate results from GNN, DNN, and LSTM. Boost SS: Boosting strategy that corrects GNN results using DNN. Boost BT: Boosting strategy that corrects GNN results using LSTM. Boost SS+BT: Sequential correction of GNN errors first using DNN, followed by LSTM. Boost BT+SS: Sequential correction of GNN errors first using LSTM, followed by DNN. MLE Typ: Maximum Likelihood Estimation results using a random starting point within the parameter space of the training dataset for each parameter’s optimization. MLE Best: MLE results using the true parameter values as the starting points for optimization. Red dashed lines in panels representing neural network results indicate the mid-points of the parameter spaces (*ŷ* = *ȳ* where *ŷ* denotes a estimated parameter and *ȳ* denotes the mid-point of the parameter space). Data points close to purple dotted lines (*ŷ* = 0) in MLE result panels indicate near-zero estimates. Black two-dash lines indicate accurate estimates (*ŷ* = *y* where *y* denotes the true parameter value). In the MLE result panels, small squares spreading along the x-axis signify optimization failures. Due to significantly lower accuracy, other aggregation methods from the bagging strategy are not displayed on the plot. *λ*: Speciation rate.

**Fig. 5.**
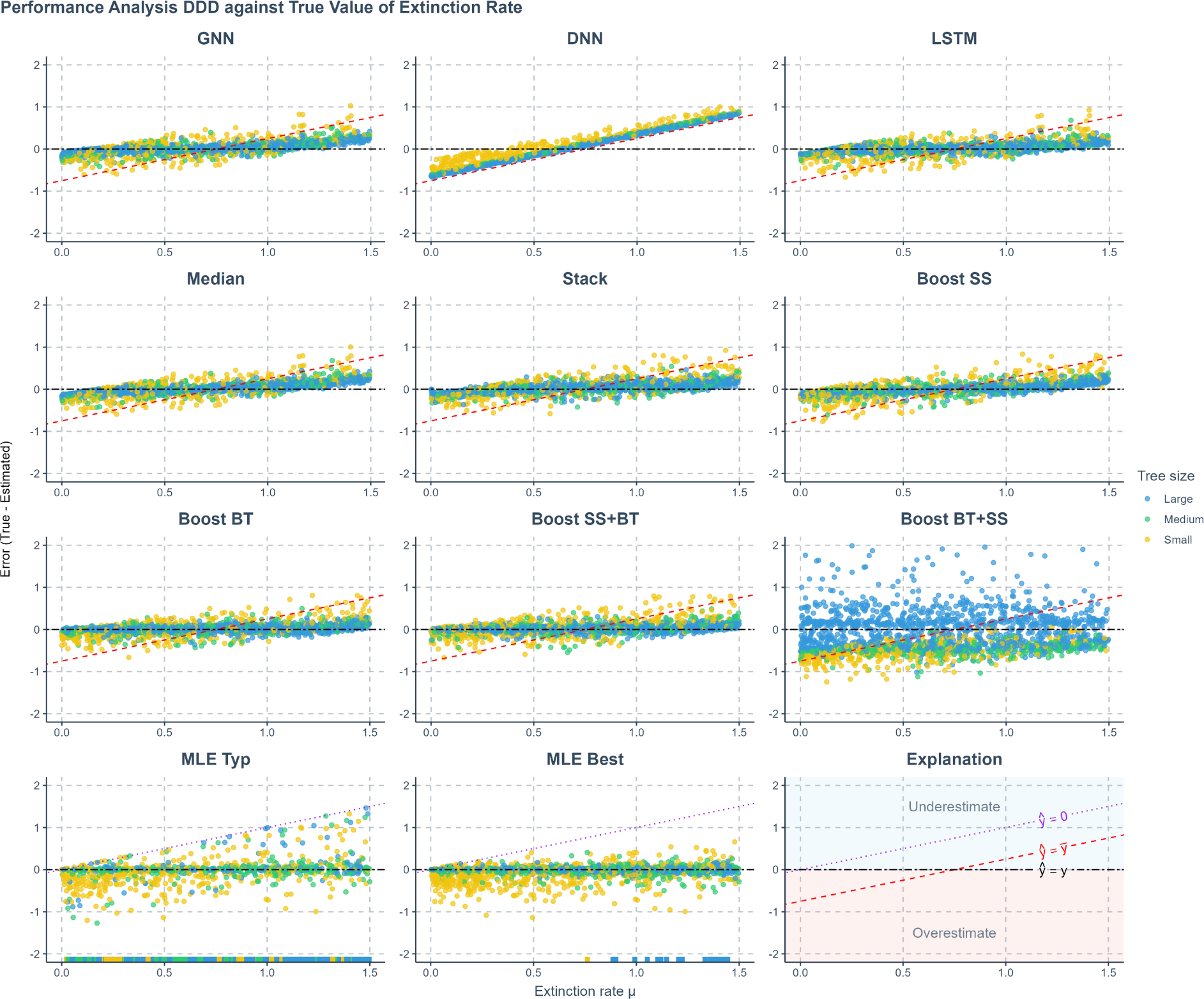
Prediction error of various methods applied to phylogenies simulated under a diversity-dependent diversification scenario, against true values of the extinction rate. The errors shown (y-axis) are the differences between the true parameters (x-axis) used to simulate the phylogenies and the values predicted or estimated by each method. Each panel represents an estimation method. Phylogenies are categorized based on their size: yellow for small phylogenies with fewer than 200 nodes (including root, internal, and tip nodes), green for medium-sized phylogenies with 200 to 500 nodes, and blue for large phylogenies with more than 500 nodes. GNN: Predictions obtained by the graph neural network using the phylogenies. DNN: Predictions by the dense neural network using summary statistics. LSTM: Predictions by the long short-term memory recurrent neural network using branching times. Median: Bagging strategy that takes the median value of the predictions from GNN, DNN, and LSTM. Stack: Stacking strategy that utilizes a meta-learner to integrate results from GNN, DNN, and LSTM. Boost SS: Boosting strategy that corrects GNN results using DNN. Boost BT: Boosting strategy that corrects GNN results using LSTM. Boost SS+BT: Sequential correction of GNN errors first using DNN, followed by LSTM. Boost BT+SS: Sequential correction of GNN errors first using LSTM, followed by DNN. MLE Typ: Maximum Likelihood Estimation results using a random starting point within the parameter space of the training dataset for each parameter’s optimization. MLE Best: MLE results using the true parameter values as the starting points for optimization. Red dashed lines in panels representing neural network results indicate the mid-points of the parameter spaces (*ŷ* = *ȳ* where *ŷ* denotes a estimated parameter and *ȳ* denotes the mid-point of the parameter space). Data points close to purple dotted lines (*ŷ* = 0) in MLE result panels indicate near-zero estimates. Black two-dash lines indicate accurate estimates (*ŷ* = *y* where *y* denotes the true parameter value). In the MLE result panels, small squares spreading along the x-axis signify optimization failures. Due to significantly lower accuracy, other aggregation methods from the bagging strategy are not displayed on the plot. *µ*: extinction rate.

**Fig. 6.**
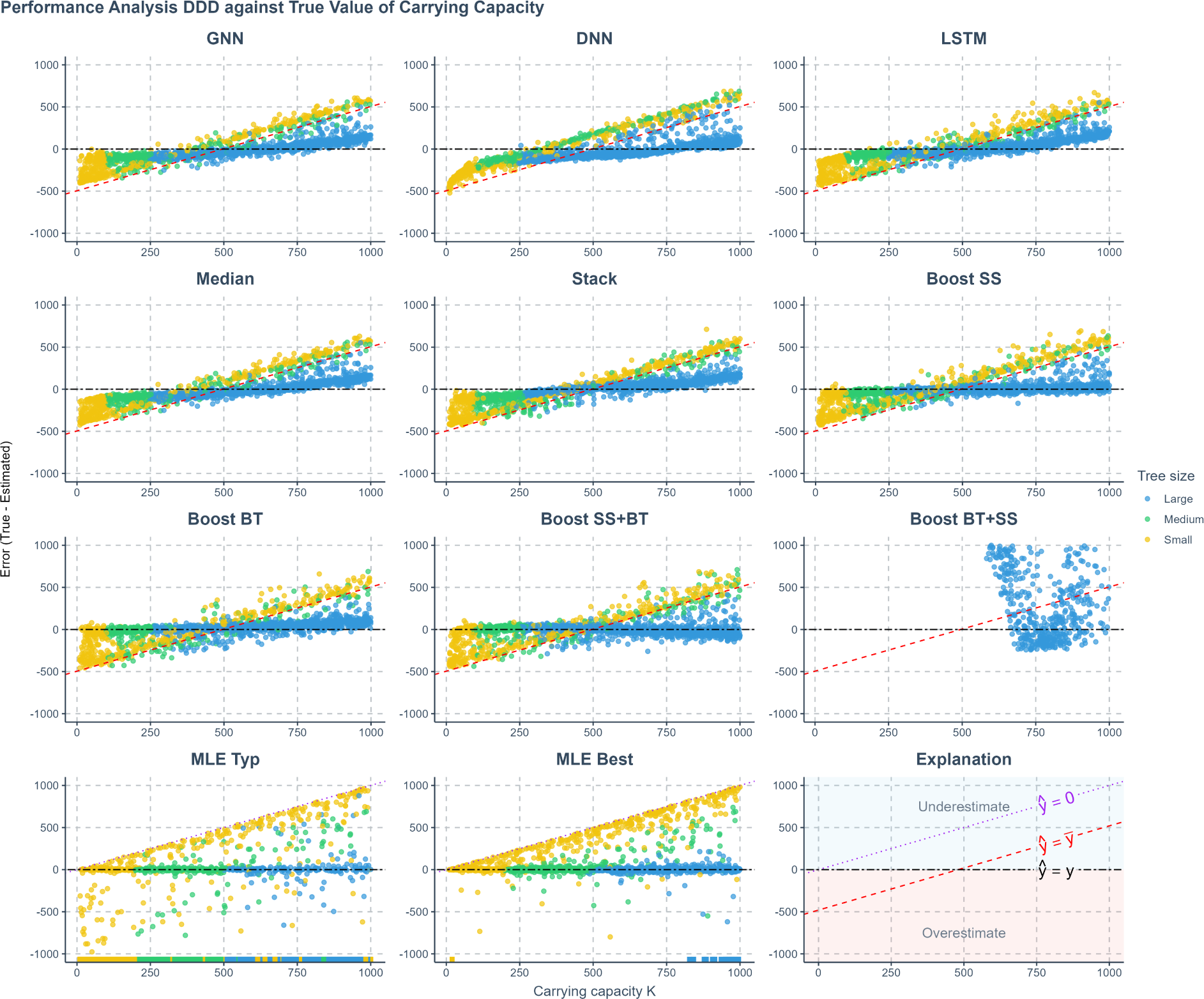
Prediction error of various methods applied to phylogenies simulated under a diversity-dependent diversification scenario, against true values of the carrying capacity. The errors shown (y-axis) are the differences between the true parameters (x-axis) used to simulate the phylogenies and the values predicted or estimated by each method. Each panel represents an estimation method. Phylogenies are categorized based on their size: yellow for small phylogenies with fewer than 200 nodes (including root, internal, and tip nodes), green for medium-sized phylogenies with 200 to 500 nodes, and blue for large phylogenies with more than 500 nodes. GNN: Predictions obtained by the graph neural network using the phylogenies. DNN: Predictions by the dense neural network using summary statistics. LSTM: Predictions by the long short-term memory recurrent neural network using branching times. Median: Bagging strategy that takes the median value of the predictions from GNN, DNN, and LSTM. Stack: Stacking strategy that utilizes a meta-learner to integrate results from GNN, DNN, and LSTM. Boost SS: Boosting strategy that corrects GNN results using DNN. Boost BT: Boosting strategy that corrects GNN results using LSTM. Boost SS+BT: Sequential correction of GNN errors first using DNN, followed by LSTM. Boost BT+SS: Sequential correction of GNN errors first using LSTM, followed by DNN. MLE Typ: Maximum Likelihood Estimation results using a random starting point within the parameter space of the training dataset for each parameter’s optimization. MLE Best: MLE results using the true parameter values as the starting points for optimization. Red dashed lines in panels representing neural network results indicate the mid-points of the parameter spaces (*ŷ* = *ȳ* where *ŷ* denotes a estimated parameter and *ȳ* denotes the mid-point of the parameter space). Data points close to purple dotted lines (*ŷ* = 0) in MLE result panels indicate near-zero estimates. Black two-dash lines indicate accurate estimates (*ŷ* = *y* where *y* denotes the true parameter value). In the MLE result panels, small squares spreading along the x-axis signify optimization failures. Due to significantly lower accuracy, other aggregation methods from the bagging strategy are not displayed on the plot. *K*: carrying capacity.

**Fig. 7.**
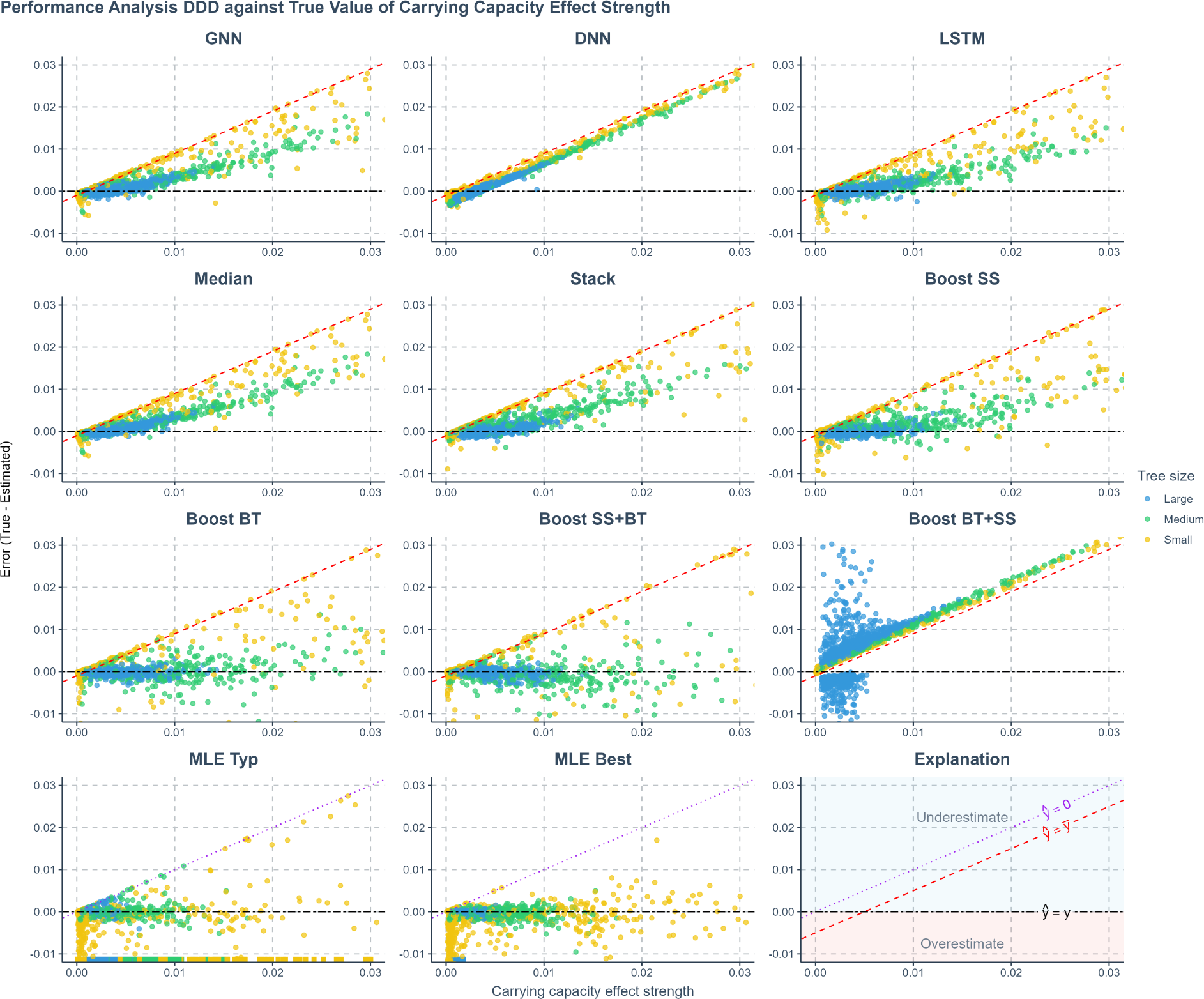
Prediction error of estimated carrying capacity effect computed from estimated values of speciation rate, extinction rate and carrying capacity using various methods applied to phylogenies simulated under a diversity-dependent diversification scenario, plotted against the true carrying capacity effect computed from true parameters. The errors shown (y-axis) are the differences between the values of true carrying capacity effect (x-axis) used to simulate the phylogenies and the values predicted or estimated by each method. Each panel represents an estimation method. Phylogenies are categorized based on their size: yellow for small phylogenies with fewer than 200 nodes (including root, internal, and tip nodes), green for medium-sized phylogenies with 200 to 500 nodes, and blue for large phylogenies with more than 500 nodes. GNN: Predictions obtained by the graph neural network using the phylogenies. DNN: Predictions by the dense neural network using summary statistics. LSTM: Predictions by the long short-term memory recurrent neural network using branching times. Stack: Stacking strategy that utilizes a meta-learner to integrate results from GNN, DNN, and LSTM. Boost SS: Boosting strategy that corrects GNN results using DNN. Boost BT: Boosting strategy that corrects GNN results using LSTM. Boost SS+BT: Sequential correction of GNN errors first using DNN, followed by LSTM. MLE Typ: Maximum Likelihood Estimation results using random starting points for parameter optimization. MLE Best: MLE results using the true parameter values as the starting points for optimization. Red dashed lines in panels representing neural network results indicate the mid-points of the parameter spaces (*ŷ* = *ȳ* where *ŷ* denotes an estimated parameter and *ȳ* denotes the mid-point of the parameter space). Data points close to purple dotted lines (*ŷ* = 0) in MLE result panels indicate near-zero estimates. Black two-dash lines indicate accurate estimates (*ŷ* = *y* where *y* denotes the true parameter value). In the MLE result panels, small squares spreading along the x-axis signify optimization failures. The title of the figure shows how the carrying capacity effect is computed, *λ*: Speciation rate. *µ*: extinction rate. *K*: carrying capacity.

**Fig. 8.**
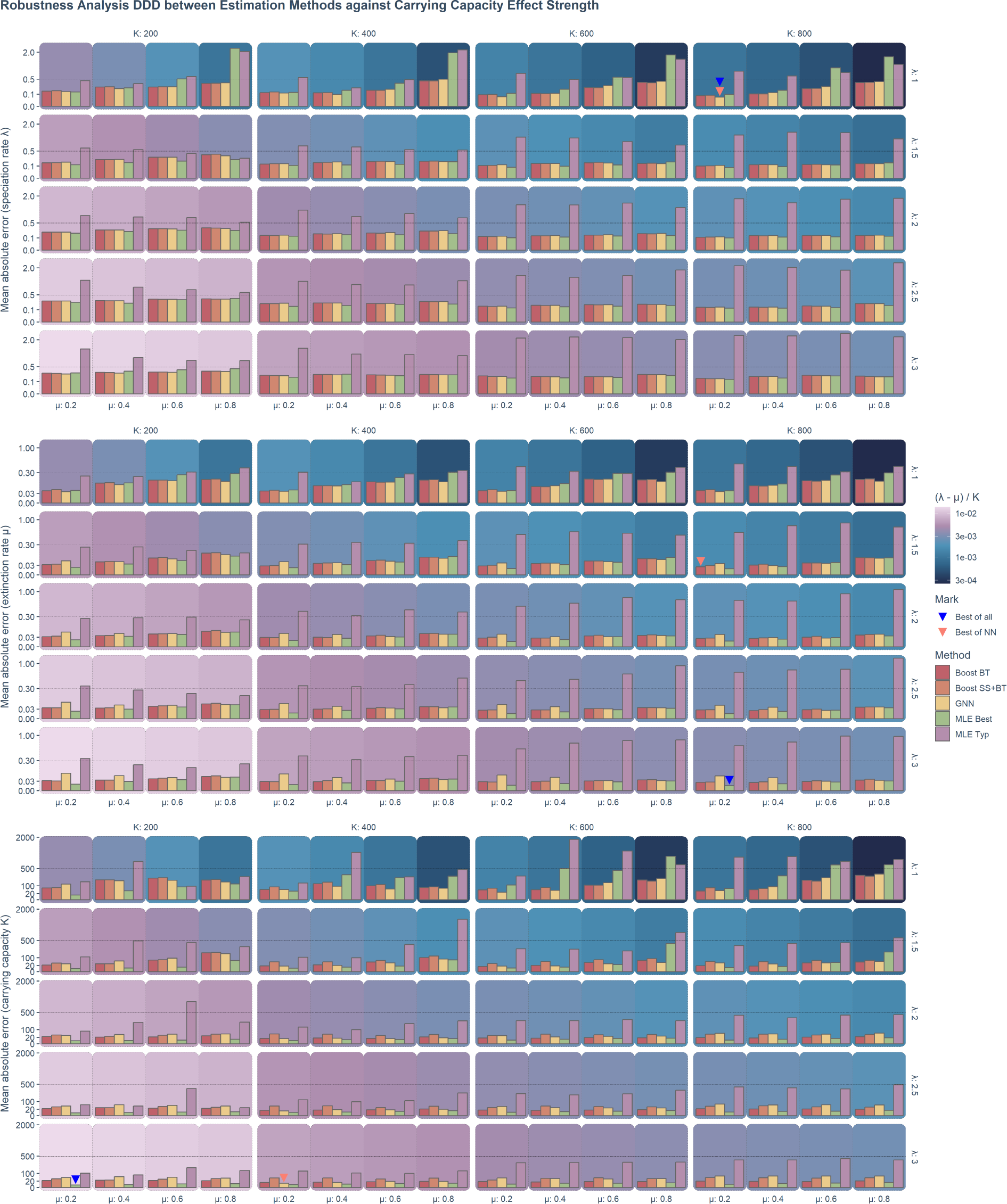
The robustness (mean absolute error) of neural network and maximum likelihood estimation was assessed for 80 sets of phylogenies, each containing 1000 trees randomly simulated under a diversity-dependent diversification scenario, employing identical parameter settings but varying in size, topology, and structure. The robustness results of the speciation rate, the extinction rate and the carrying capacity are shown from top to bottom. For each panel group associated to a parameter, each panel contains the robustness of different estimation methods (the MLE and neural networks) under a combination of parameters indicated by the facet strip labels. Each facet column represents the robustness under a specific carrying capacity (*K*) setting used in the simulation of the phylogenies. Each facet row represents a specific speciation rate (*λ*). Each group of the bars represents a specific extinction rate (*µ*) as shown by the x-axis. The background color of a panel represents the carrying capacity effect strength (calculated as (*λ − µ*)*/K* and visualized in “log10” scale), from bottom-left to top-right, the carrying capacity effect strength decreases. The color of a bar represents the associated estimation method. Boosting BT: Graph neural network with long short-term memory recurrent neural network correcting its residuals using branching times. Boosting SS + BT: Graph neural network with dense neural network and long short-term memory recurrent neural network correcting residuals sequentially using summary statistics and branching times. GNN: Graph neural network. MLE Best: Maximum likelihood estimation using true parameters as the starting points. MLE Typ: Maximum likelihood estimation using a random value as the starting point of optimization for each parameter. X-axis: Represents extinction rate (*µ*) settings. Y-axis: Represents the mean absolute error in a square-root transformed scale. Some bars are marked; for each parameter, the blue triangle represents the greatest possible robustness achieved among all the estimation methods, the red triangle represents the greatest possible robustness achieved among the neural network methods.

The MLE implementation sometimes fails to find an optimal solution. In our visualizations, failed MLE estimations are indicated by small squares spreading along the x-axis to avoid misinterpretation. MLE tended to give small or near-zero estimates, particularly on the extinction rate and the carrying capacity. This phenomenon is more prominent when starting optimization from a random point. For all figures showing the MLE error, the ideal situation is that all the data points lie near the horizontal black two-dash reference lines (at which the error is 0) and do not spread along or near the purple dotted reference lines (which suggests near-zero MLE estimations). See the last two panels of Figure 4, Figure 5 and Figure 6 for details.

Neural networks often return values closer to the parameter space’s mid-points (indicated by red dashed lines), a result of making “safer” predictions that minimize loss compared to random guesses. Consequently, neural networks usually overestimate at low true values and underestimate at high true values (see Figure 4, Figure 5 and Figure 6). These errors are mitigated or partially corrected when the neural networks are trained in tandem through boosting strategies, e.g. boosting GNN results with DNN or LSTM or both (see the panels of Boost SS, Boost BT and Boost SS+BT in Figure 4, Figure 5 and Figure 6). This happens particularly for large phylogenies (the blue data points in Appendix G, Figure 14) when the underlying true carrying capacity (*K*) is large, or for small phylogenies (the yellow data points) when the underlying true speciation rate (*λ*) is small.

However, boosting strategies can introduce their own challenges. When boosting GNN results first with LSTM and then with DNN, the DNN failed to identify a general pattern of errors from LSTM results. This led to overfitting on the training dataset at the second epoch of the training session (the total loss in the validation dataset started to increase and became much larger than the total loss in the training dataset), which, in turn, resulted in poor performance on the testing dataset (see the panels named Boost BT+SS in Figure 4, Figure 5, Figure 6 and Appendix G, Figure 14).

Upon further analysis of the residuals, we observed that inaccuracies in the predictions were largely influenced by the size of the phylogeny (Figure 4, Figure 5 and Figure 6). For neural network approaches, the prediction errors for speciation rate, extinction rate, and carrying capacity tended to increase as the size of the phylogeny decreased, especially in phylogenies with fewer than 200 nodes. Systematic error was also identified in the estimation of carrying capacity: neural networks generally overestimated this parameter in smaller phylogenies and underestimated it in larger ones. Boosting strategies were effective in mitigating or partially correcting systematic errors, and enhancing prediction accuracy, particularly for carrying capacity (see the rows of Boost SS, Boost BT and Boost SS+BT in Appendix G, Figure 14).

We calculated the strength of the carrying capacity effect using the formula 1*/K^t^* = (*λ − µ*)*/K*, where *λ* represents the true speciation rate, *µ* the true extinction rate, *K* the true carrying capacity, and *K^t^* the diversity at which speciation becomes zero for linear negative diversity-dependence (Etienne et al., 2012). A larger (*λ − µ*)*/K* value corresponds to smaller *K^t^* and therefore a stronger carrying capacity effect. Phylogenies exhibiting a stronger carrying capacity effect typically have more accurate estimates.

However, in the case of smaller phylogenies, neural networks tended to underestimate speciation and extinction rates while overestimating carrying capacity when the carrying capacity effect is weak, and the reverse is observed when the effect is strong. In contrast, MLE tends to overestimate speciation and extinction rates while underestimating carrying capacity under conditions of weak carrying capacity effect, with the reverse occurring under strong effects, except for the carrying capacity which is always underestimated (see Appendix G, Figure 14). Neural network methods tend to underestimate the carrying capacity effect. This phenomenon can be mitigated by the boosting strategies, especially the Boost BT method, which achieved similar performance to the best case MLE estimates (see Figure 7).

Unlike GNN and LSTM, DNN cannot by itself reliably recover speciation and extinction rates from the summary statistics of the phylogenies, with its predictions mostly clustering around the mid-points of the parameter space (around the red dashed lines in the DNN panels in Figure 4, Figure 5 and Figure 6). The overall accuracy of the carrying capacities recovered by the DNN is also inferior compared to the other approaches (see the row named DNN in Appendix G, Figure 14).

Among all ensemble learning strategies, boosting consistently outperformed both bagging and stacking in enhancing prediction accuracy compared to using neural networks independently, as can be seen, for instance, by the lower mean absolute prediction errors in Appendix G, Figure 14. Boosting strategies also exhibited better performance in recovering the true values of the carrying capacity effect (see Figure 7). The most effective neural network approaches overall matched or even surpassed the results of MLE while exhibiting no bias, even on smaller phylogenies. Overall, sequential boosting of GNN results first with DNN and then with LSTM (Boost SS+BT) led to best performance in terms of prediction accuracy, except for estimating carrying capacity on large phylogenies with around 2000 nodes (see the row named Boost SS+BT in Appendix G, Figure 14 this strategy led to overestimation on very large phylogenies). However, Boost SS+BT led to more overestimation on the true values of the carrying capacity effect, as compared to the strategy boosting the GNN results with only LSTM (Boost BT, see Figure 7).

### Robustness Analysis

As a proxy for robustness of each method, we used the mean absolute errors of the parameters estimated from sets of phylogenies simulated under identical true parameters. Our analysis indicates that the robustness of the methods against phylogenetic heterogeneity (e.g., phylogenies of very different sizes, topologies and other characteristics) depends on the values of the underlying true parameters. We observed that the strength of the carrying capacity effect critically influences robustness. Generally, a weaker carrying capacity effect (associated to a smaller value of (*λ − µ*)*/K*) tends to diminish the robustness of both MLE and neural network methods across all parameters: speciation rate, extinction rate, and carrying capacity (as can be seen in Figure 8 and Appendix G, Figure 16 and Figure 15, by observing the increase of error along with the darkening background colors from light pink to dark blue).

When the carrying capacity effect is weak, neural network methods typically exhibit greater robustness in estimating speciation and extinction rates compared to the best-case MLE results (see Figure 8 and Appendix G, Figure 16). When the carrying capacity effect is exceptionally strong, the best-case MLE results can outperform neural networks particularly when estimating carrying capacity. Typical-case MLE results consistently show less robustness compared to all neural network methods.

A higher extinction rate generally decreases the robustness of all methods in estimating any parameter. A higher speciation rate enhances the robustness of carrying capacity estimates across all methods, although its impact on the robustness of speciation and extinction rate estimates is not consistent. A higher carrying capacity generally decreases the robustness of all methods in estimating carrying capacity.

Note that MLE typical-case results often contained more extreme estimations than best-case results, consequently, the exclusion of extreme values could lead to a wrong impression in the figures that when the carrying capacity effect is weak, the typical-case MLE is more robust than the best-case MLE. This is particularly prominent for the speciation rate. The exclusion of these extreme values is crucial, however, as they are rare and their magnitude can obstruct meaningful interpretation and comparison.

We find that DNN alone (estimating parameters from summary statistics) shows the worst robustness among all the methods and LSTM alone (estimating parameters from branching times) shows the greatest robustness overall. Among all the estimation methods, the MLE best-case achieved the greatest possible robustness in estimating the extinction rate and the carrying capacity while GNN alone (estimating parameters from phylogenies) achieved the greatest possible robustness in estimating the speciation rate. Among the neural network methods, GNN alone achieved the greatest possible robustness in estimating the speciation rate and the carrying capacity while Boost BT (boosting GNN estimates with LSTM) achieved the the greatest possible robustness in estimating the extinction rate. See Figure 8 and Appendix G, Figure 15 and Figure 16 for details.

### Empirical Data

MLE estimates of carrying capacity are typically lower than those of the neural networks, especially in smaller phylogenies (Figure 9). However, as the size of the phylogenies increases, MLE estimates tend to converge towards those produced by neural networks. Similarly, MLE estimates of net diversification rate (computed as *λ − µ*) also align more closely with neural network estimates in larger phylogenies Figure 9.

**Fig. 9.**
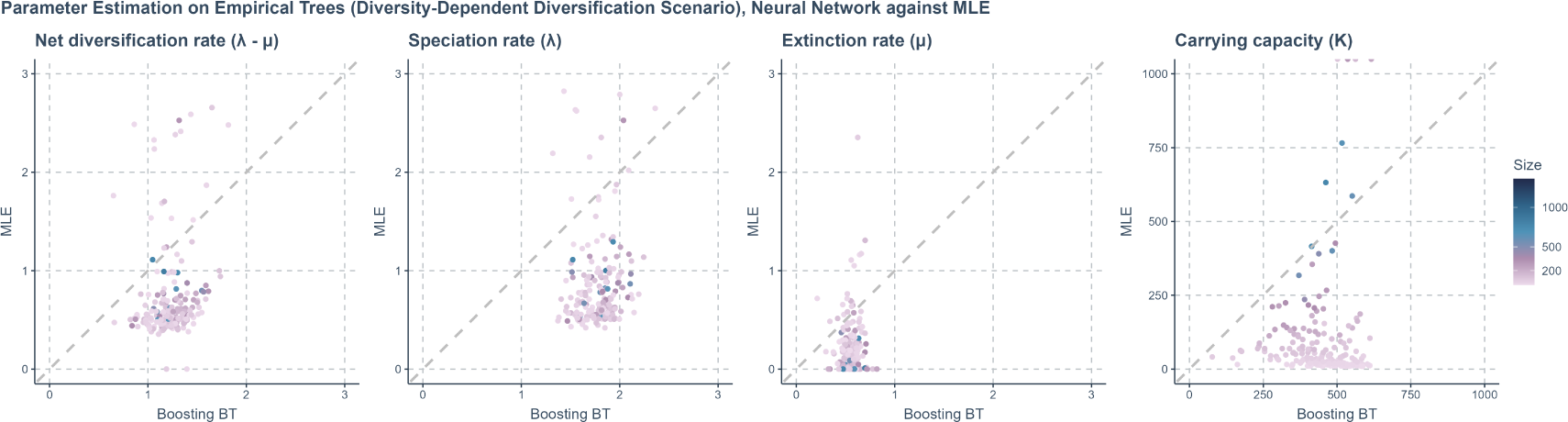
Comparison of the estimations of Maximum Likelihood Estimation (MLE) and neural network methods (specifically, Boosting BT, which refers to using graph neural network to make first predictions and then using a long short-term memory recurrent neural network to correct for residuals) on empirical trees under a diversity-dependent diversification scenario. Each panel, arranged from left to right, focuses on a specific parameter being estimated. X-axis: Represents the estimated values of the neural network. Y-axis: Represents estimated the values from MLE. A gray dashed line is included in each panel to indicate where the estimations from the neural network and MLE are exactly the same. The color of the points varies from purple to blue, with the gradient representing the size of the phylogenies measured by the total number of nodes (including root, internal, and tip nodes).

**Fig. 10.**
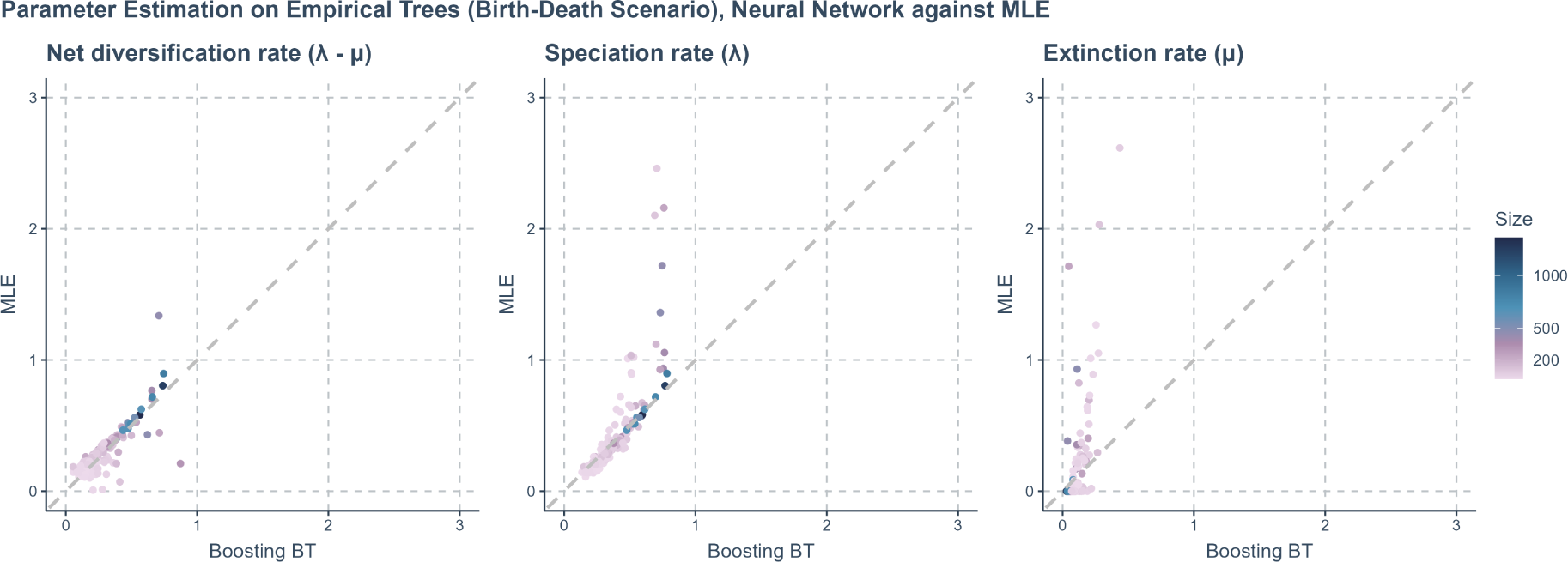
Comparing the estimations of Maximum Likelihood Estimation (MLE) and neural network methods (specifically, Boosting BT, which refers to using graph neural network to make first predictions and then using long short-term memory recurrent neural network to correct for residuals) on empirical trees under a birth-death scenario. Each panel, arranged from left to right, focuses on a specific parameter being estimated. X-axis: Represents the estimated values of the neural network. Y-axis: Represents estimated the values from MLE. A gray dashed line is included in each panel to indicate where the estimations from the neural network and MLE are exactly the same. The color of the points varies from purple to blue, with the gradient representing the size of the phylogenies measured by the total number of nodes (including root, internal, and tip nodes).

MLE generally provides a broader range of estimates on all the parameters except for carrying capacity on small phylogenies. Neural networks provide a broader range of carrying capacity estimates on small phylogenies and less frequently produce zero or near-zero estimates for extinction rates which we often observe for MLE. We also observed that MLE sometimes produces extreme values (ranging from 10,000 to infinity) for carrying capacity on empirical trees, see Figure 9 for the comparison between MLE and neural networks on empirical tree parameter estimation.

Generally, neural network estimates of all the parameters under the DDD scenario are close to the center (mean) of the distribution generated by the bootstrapping method. See Figure 13 in Appendix F for details.

### Other Scenarios

*Birth-Death Scenario* Neural network methods outperformed MLE in accuracy, particularly on smaller phylogenies, under the BD scenario in the simulated dataset (see Appendix H, Figure 19 and Figure 18). Both MLE and neural network methods give less accurate estimates on small phylogenies; this is more prominent for the MLE estimates.

On empirical phylogenies, similar to the DDD scenario, neural network methods seldom produce zero-estimation of extinction rate, unlike MLE, which often gives zero or near-zero estimation on extinction rate. Neural networks tend to give estimates within the parameter space of the training dataset. They predict conservative speciation and extinction rates yet are highly consistent with MLE estimation on the net diversification rate, defined as the difference between speciation and extinction rates (*λ − µ*). The consistency of prediction increases on larger empirical phylogenies. See Figure 10 for details.

### Protracted Birth-Death Scenario

Both maximum likelihood estimation (MLE) and neural network methods did not perform well on estimating parameters under the PBD scenario; MLE estimates were generally less accurate, but neural networks also failed to predict the parameters as all the parameter estimates are close to the mid-points of corresponding parameter spaces (Appendix I, Figure 20,). However, there are exceptions: neural networks seem to perform better on the speciation rate of the incipient species (*λ*_3_) and on the mean duration of speciation (*τ*) when the true value is between 0 and 2.

MLE estimates become significantly inaccurate as phylogenies become increasingly small (Appendix I, Figure 21); it is also noticeable that MLE estimates of the speciation completion rate (*λ*_2_) are very inaccurate, especially when the phylogenies are large. A general pattern is that both MLE and neural network methods achieve more accurate estimates on phylogenies with higher true values of the mean duration of speciation (Appendix I, Figure 22).

## Discussion

We have developed an ensemble learning based neural network approach that matches and sometimes outperforms the accuracy and robustness of maximum likelihood estimation (MLE) for estimating phylogenetic tree parameters. Our approach leverages different classes of neural networks by learning from the phylogenies, their branching times and their summary statistics simultaneously.

When trained, our neural networks can compute estimates faster than MLE on larger phylogenies as computation time is less affected by increases in phylogeny size. We considered boosting strategies most effective in eliminating systematic prediction errors in neural network estimates. Among them, Boost BT (which corrects GNN results using LSTM) achieved overall best performance which is comparable to, or even surpassing, the best case MLE, in terms of accuracy and robustness. We observed that generally the performance of the typical MLE was second-worst (Boost BT+SS was worst). Interestingly, some phylogenies, such as small trees and those shaped by relatively weak effects, pose significant challenges to both MLE and neural network methods.

Previous neural network methods applied to phylogenies have experimented with various architectures such as convolutional neural networks (CNN), GNN, and LSTM (Lajaaiti et al., 2023; Lambert et al., 2023; Voznica et al., 2022). The deep learning architectures we employed differ from those used in prior studies, making direct comparisons challenging. Additionally, while previous research focused on birth-death (Lambert et al., 2023) and trait-state-dependent models (Lajaaiti et al., 2023), our approach is novel in its application to models such as diversity-dependent diversification (DDD) and protracted birth-death (PBD) from a neural network perspective. Despite these differences, our findings align with recent studies in underscoring the potential of neural networks to infer diversification processes, offering a viable alternative to mathematically complex methods.

### Rethinking Neural Networks

Although performance was equal or better than MLE, the neural network approach is not without its shortcomings. The neural networks often defaulted to predicting values close to the mid-point of the true parameter space of the training dataset, indicating that they struggle to extract meaningful features from the dataset. This conservative prediction strategy minimizes overall error compared to random guessing. Examples can be found in the GNN predictions of carrying capacity from simulated DDD trees, the DNN estimates of the speciation and extinction rates (Figure 4, Figure 5 and Figure 6, but see Appendix L for a detailed investigation of possible under-performance of DNN on the summary statistics) and most neural network predictions of PBD related parameters (Appendix I, Figure 20), especially for smaller phylogenies.

This behavior, while effective in reducing apparent error metrics, can skew our understanding of a neural networks performance particularly when the focal parameter space is relatively narrow. Neural networks may consistently show smaller overall error compared to MLE, because the latter has no prior knowledge of the limits of the parameter space, which would lead to a false impression of better accuracy of the neural networks. We therefore recommend performing case-specific residual analyses on the neural network predictions and the MLE estimates, which are often overlooked or over-simplified.

We propose two strategies to minimize the influence of possible under-representing training dataset. The first strategy is to estimate parameters using MLE (if possible) and then training neural networks with a dataset generated by true parameter values that cover the MLE estimates, if the estimates seem reasonable. When MLE does not exist, or the estimates seem unrealistic, we can use the second strategy, that is, to train the neural networks and predict parameters on a relatively narrow training dataset, then retrain on a broader dataset generated from a parameter space with different means. We can examine if our predictions are subject to the range of the training dataset by observing whether the prediction changes. Generally, we recommend to train the neural networks with as large a dataset (sample size) and as broad a parameter space as possible.

Improving neural network predictions that are close to the mean is unlikely to be achieved by increasing the amount of training data: we did not observe major performance improvement when changing the size of the datasets (from 1,000 to 100,000 phylogenies per dataset). Instead, one might consider increasing the complexity of the network architecture, such as increasing their depth or adapting the scale of the hidden nodes (Zhang et al., 2021), but note that this may require larger data.

Although potentially beneficial, increasing the depth can also harm predictive power. In particular for GNNs, increasing their depth may lead to “over-smoothing” and “over-squashing”. Over-smoothing causes node features to become increasingly similar as more layers are added (Li et al., 2018), leading to a loss of distinct node embeddings across different clusters. Over-squashing involves the compression of expansive node information through bottleneck edges into a fixed-size vector, which is problematic in graphs with large diameters and long-range dependencies (Alon and Yahav, 2021), e.g.phylogenies. Both issues degrade node representations and distort information flow, making deeper GNNs potentially less effective than shallower ones (Dwivedi et al., 2022). Moreover, over-smoothing and over-squashing are intrinsically linked, creating a trade-off that cannot be easily resolved (Giraldo et al., 2023).

In our analyses, we observed that increasing the number of GraphSAGE layers beyond three in the differentiable pooling architecture destabilized the training process and reduced the accuracy of estimates on validation datasets, introducing more outliers.

We therefore opted to maintain two layers throughout our study. We explored newer algorithms designed to mitigate deep GNN issues (Chen et al., 2020; Gravina et al., 2022; Li et al., 2018), but found that these deeper architectures performed worse than our differentiable pooling approach with fewer layers. For DNN and LSTM, we also experimented with more complex architectures, different activation functions and various hyper-parameter optimizations but failed to achieve better performance.

### Fundamental Problems with Phylogenies

The lack of improvement when changing the amount of training data or the network architecture suggests that the real challenges of estimating parameters might not lie in the architecture of the networks, but might instead be attributed to underlying weak or absent phylogenetic signals. Whenever this is the case, we expect similarities in inaccuracies of both MLE and neural network approaches. This occurs, for example, for the carrying capacity when it is high and thus has a weak effect (measured by (*λ − µ*)*/K*).

Here, the phylogeny is typically not near the carrying capacity, allowing the number of species to grow (almost) unbounded. This may result in carrying capacity estimates that are arbitrarily high, especially in the MLE methods. The PBD scenario is known to present difficulties in recovering parameters reliably with MLE (Etienne et al., 2014) and we find similar poor performance with neural networks. A second case where accurate parameter estimation is complicated occurs when extinction processes erase critical information (Louca and Pennell, 2021), as observed in the decline of estimation robustness associated with increasing extinction rate.

More generally, small phylogenies tend to contain less information than large ones. In our results we see that estimation accuracy and robustness decline with decreasing size of the phylogenies. This trend is observed across both MLE and neural network methods. In the BD and PBD scenarios, where datasets have greater variability in phylogeny sizes, poor estimations for small trees could be explained by both low information content, or under-representation of such trees. To account for the potential effects of under- and over-representation of phylogenies of different sizes in our datasets, we conducted a supplementary study to explore whether the patterns we observed persist in a dataset with a re-balanced distribution of phylogeny sizes (see Appendix J, Figure 24). This shows the same patterns and they are therefore unlikely to be a result of under- or over-representation of different phylogeny sizes, and instead reflect low information content of small trees (compare Figure 18 and Figure 24).

Low information content for specific parameters is expected to be more likely as the complexity of the underlying diversification models increases (see the results of BD in Appendix H, the results of DDD and the results of PBD in Appendix I, from BD to DDD to PBD, the complexity increases). This difficulty may stem from the increasingly complex and substantial signals that the phylogenies are required to convey, which may not be fully captured in the stochastically generated data. When applying these methods to empirical phylogenies, there is a noticeable decline in the agreement between MLE and neural network estimates from the BD scenario to the DDD scenario.

### Confronting the Empirical Phylogenies

The processes of evolution within natural systems are often unknown. Determining the “true parameters” of an empirical phylogeny is challenging, even when they meet theoretical assumptions, making it difficult to evaluate which tool provides more accurate estimates. Therefore, choosing the right tool is crucial.

With neural networks, it is possible that the true parameter value is not part of the assumed parameter range for simulating the training data. In such cases, neural network accuracy decreases notably, as shown in a supplementary study (explained in Appendix K). We also noticed that when comparing the estimates of MLE and neural network methods on the empirical phylogenies (see Figure 9, Figure 10), MLE estimates spread wider than the neural networks (e.g. our BD training dataset comprises phylogenies simulated using speciation rate between 0 and 0.8 and extinction rate between 0 and 0.72, our neural networks never predict speciation rate larger than 0.8 or extinction rate larger than 0.72, see Figure 10, similar results can be found under the DDD scenario in Figure 9). Expanding the training dataset’s parameter space can resolve the generalization issue (we expanded our training datasets several times in the experiments), but this approach requires significantly more computational resources for both simulation and training of the neural networks.

Our supplementary study (explained in Appendix K) also reveals that neural networks tend to provide more accurate estimates of speciation and extinction rates from complete phylogenies than from extant ones under the BD scenario. This increase in accuracy was not observed in the DDD scenario. While complete phylogenies offer a broader picture and more contextual information, obtaining them is challenging because it is nearly impossible to account for all extinct species.

Our analyses indicate that GNN is more robust but more prone to systematic errors (GNN achieved the greatest possible robustness in estimating the speciation rate and carrying capacity among neural network methods). We show that using GNN as a base and other neural networks like LSTM to enhance GNN might effectively combine the advantages of different methods and information sources, thus strengthening overall generalization ability. Our boosting methods (e.g. Boost BT) perform the best in this context.

In conclusion, when applied with caution, neural network methods can be applied to other diversification scenarios where MLE is absent or non-tractable, as our best-performing neural network method showed comparable or even better performance to the best-case MLE. Our neural networks particularly perform better than MLE in terms of accuracy and robustness on small phylogenies and can be significantly faster when estimating very large phylogenies. Thus, if properly trained, neural network methods may substitute for or at least cross-reference with MLE estimates where they exist.

## Acknowledgements

We thank the Center for Information Technology of the University of Groningen for their support and for providing access to the Hábrók high performance computing cluster.

## APPENDIX

### A. Data Transformation Protocol

#### Protocol Transforming Phylogenies for Graph Neural Network

Phylogenetic trees are usually stored in the “phylo” data format in R. This data format is not directly compatible with GNN implementations. To facilitate graph convolutional operations, we transformed phylogeny from a “phylo” object into three major components: adjacency list, node feature matrix and graph attributes (see Figure 2). The adjacency list contains information on the connectivity between nodes and tips, the node feature matrix contains distances between nodes and tips, and the graph-level attributes include the true initial values to generate the phylogenies. These components are stored in separate tensors. In machine learning, a tensor is a mathematical object that generalizes scalars, vectors, and matrices to higher dimensions, allowing complex operations to be performed efficiently on multi-dimensional arrays.

#### Adjacency List

In the context of a phylogenetic tree, tip nodes usually represent taxonomic units such as species, while root nodes and internal nodes represent the points where two taxonomic units depart from each other. An edge in a phylogenetic tree represents the connection between two nodes, and as such describes the evolutionary relatedness between taxa. Each root node, internal node and tip node in an R “phylo” object is indexed sequentially, each edge is also sequentially indexed independently of node indices. The sub-list “edge” of a “phylo” object contains the adjacency list of a phylogenetic tree which describes the relationships between nodes. Each row of the adjacency list represents an edge, the first column contains the index (or numbering) of the ancestor node, and the second column contains the index of the descendant node.

This data structure effectively captures the tree’s branching pattern, showing how each taxon (or node) is connected to others. The adjacency list in “phylo” object uses a “1-based” indexing in R, we therefore element-wise deduct 1 from the list to convert it into “0-based” indexing which is compatible with the python environment.

We output the converted adjacency list within the “phylo” object as the adjacency list *E* of the graph representation, in PyTorch Geometric, which is conventionally named as “data.edge_index”. We store *E* as a “torch.long” long integer type tensor and transpose it such that it has shape [2*, num*_*edges*], where “num_edges” is the number of edges in the “phylo” object. This tensor has two dimensions. This way, the connections between nodes in the transformed graph are all single-directional, from the ancestor nodes to their descendants (if any). Training the GNN with graphs of non-directed edges gives no performance advantage, according to our tests in phylogenetic tree parameter estimation tasks. Single-directional data structure can save GPU memory and reduce the computation complexity.

#### Node Feature Matrix

In a “phylo” object in R, the “edge.length” sub-list defines the lengths of the edges in the phylogenetic tree. In a phylogenetic context, these lengths often correspond to evolutionary distances, time, or genetic change. “edge.length” is a numeric vector where each element corresponds to the length of the edge as defined in the adjacency list. The order of lengths in the “edge.length” vector aligns with the order of edges in the adjacency list.

For each tree, we aggregate information contained in “edge.length” to a node feature matrix. Each row of the matrix represents features contained in a node. The first column contains the edge length from a node to its direct ancestor node, the second and the third columns contain the edge lengths from a node to its two daughter nodes. We pad the row of the root node with an 0 in the first column as it has no ancestor. We also pad the rows of the tip nodes with two 0s in the second and the third columns as they have no descendants. The row order of feature matrix aligns with the order of edges in the adjacency list.

We output the node feature matrix of each tree as the node feature matrix *X* of the graph representation, in PyTorch Geometric, this is conventionally named as “data.x”. We store *X* as a “torch.float” floating point type tensor, it has shape [*num*_*nodes, num*_*node*_*features*], where “num_nodes” is the number of nodes (including tip nodes) in the “phylo” object and “num_node_features” in our case is 3, i.e. the phylogenetic distances from a node to its ancestor (if any) and two descendants (if any). This tensor has two dimensions. We do not store the phylogenetic distance information in edge features because GCN operators will eventually pass and aggregate the edge features into each of the node. Our data structure is simpler and so is the GNN architecture.

#### Graph-Level Attributes as Training Targets

We store all the parameters used to simulate a tree (ground truth values) in the graph-level attributes *Y*. These can have arbitrary length, which should be consistent with the number of the parameters to be estimated (the three diversification scenarios, BD, DDD and PBD, have different number of parameters). We store graph-level attributes as a “torch.float” floating point type tensor with length of the number of parameters we want to predict for each type of the phylogenetic tree. In PyTorch Geometric, graph-level attributes can be named as “data.y”. The graph-level attributes are used as training targets to compute loss (see Appendix B for the definition of loss).

#### Protocol Transforming Summary Statistics for Dense Neural Network

The summary statistics of a phylogeny are represented by a 1D vector, so the protocol for DNN is straightforward: we convert the vector into a tensor containing floating type data, with the shape [num_stats], where “num_stats” denotes the total number of statistics. This tensor has only one dimension. This conversion guarantees that each tensor is associated with its respective tree, with all contained statistics maintaining their original order. Within the PyTorch Geometric framework, these statistics are encapsulated as “data.stats” for each tree. When using DNN alone to estimate parameters from the summary statistics, the ground truth values of the parameters of the trees are stored in the same way as the graph-level attributes, as model training targets. When using DNN with other neural networks (e.g. in stacking and boosting strategies), they share the same ground truth values which are the graph-level attributes.

#### Protocol Transforming Branching Times for Recurrent Neural Network

To address the varying lengths in branching times across different phylogenetic trees, we standardize these sequences by padding them to match the length of the longest branching time sequence. This is achieved by appending zeros to the shorter sequences until they match the predefined maximum length. The padded sequences are stored in tensors containing floating type data. As the original branching times do not contain zero values, this padding strategy allows us to distinguish between original data and padding.

Consequently, we can pass masks of the sequences to the LSTM, which indicates the positions of the paddings, making LSTM concentrate only on the informative portions of the sequences, thereby optimizing its performance. When using LSTM alone to estimate parameters from the branching times, the ground truth values of the parameters of the trees are stored in the same way as the graph-level attributes, as model training targets. When using LSTM with other neural networks (e.g. in stacking and boosting strategies), they share the same ground truth values which are the graph-level attributes.

### B. Total Loss

Total loss comprises three key components: Huber loss, link prediction loss and entropy of regularization. Huber loss was used for optimizing regression accuracy while the remaining components focused on alleviating a possible issue where GNN can be hard to train, if incorporating the differentiable pooling method (Ying et al., 2018).

The Huber loss (Huber, 1992) for vectors *y* and *ŷ*, each with *n* elements, computed as the average loss across all elements, is given by:

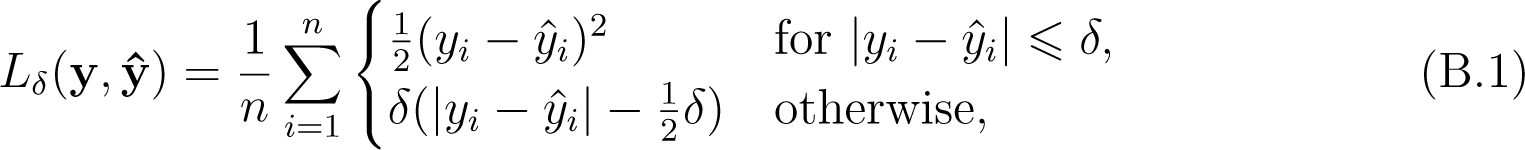

where y is the true value vector comprising the ground truth parameters used for simulating a phylogenetic tree, ŷ is the predicted value vector comprising the parameter predictions, *y_i_* and *ŷ_i_* are the *i*-th elements of y and ŷ respectively, *n* is the number of elements in the vectors y and ŷ and *δ* is the threshold parameter that defines the transition from squared to linear loss (here loss refers to the difference between ground truth and predicted values). In our research, we set *δ* = 0.8 for all the training sessions, making the neural networks more sensitive to smaller errors and more robust to outliers.

The total loss *L* is given by

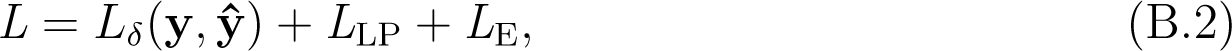

where *L*_LP_ is the link prediction loss and *L*_E_ is the entropy of regularization, see Ying et al. (2018) for their definitions.

### C. Neural Network Architecture

For the graph neural network, we used GraphSAGE (Hamilton et al., 2017), a sample-and-aggregate graph convolutional neural network, to capture a graph-level representation. GraphSAGE has achieved strong performance of learning from large graphs. We use graph neural network (GNN) to refer to the graph neural network approach which incorporates GraphSAGE.

GNN is mainly assembled from five GNN modules (see Figure 1-C for five blocks of boxes in yellow and orange colors). Each module comprises the same number of GraphSAGE operators (Hamilton et al., 2017), where the number of layers (GraphSAGE operators, as illustrated by the number of combined boxes within each GNN modules in Figure 1-C) *N_L_* = 1, 2*, …,* 6. Each GNN operator is accompanied by a Batch Normalization for 1D Inputs (BatchNorm1d, not shown in Figure 1) operator (Ioffe and Szegedy, 2015) and then a Gaussian Error Linear Units (GELU, as illustrated by the orange bands within the yellow boxes in Figure 1-C) activation function (Hendrycks and Gimpel, 2016). The GraphSAGE operators facilitate the convolution operation over graphs, capturing both local node features and their neighborhood information. The BatchNorm1d operator is commonly employed in neural networks to stabilize and accelerate the training process. The GELU activation layer is used for introducing non-linearity into the data. Learned features from all the GraphSAGE operators within a module are collected and concatenated. Larger *N_L_* will result in the GNN modules to aggregate information into each node from its more distantly connected neighbors. According to our experiments, the optimal case is *N_L_* = 2, all figures and results relating to GNN were reported on the optimal case.

The graph-learning process also involves graph coarsening operations. We incorporated the differentiable pooling (DiffPool hereafter) technique to better learn hierarchical representations of the graphs. DiffPool can aggregate graph nodes into clusters after each operation. It facilitates graph coarsening and captures intricate hierarchical structure, which makes it particularly suitable for graph-level tasks (Ying et al., 2018). In the first coarsening operation, the graph data inputs are passed to two GNN modules (pooling and embedding, see Figure 1-C for the blocks marked as “GNN pool1” and “GNN embed1”). The pooling group reduces the graph size, while the embedding group captures the node features. The filtered data from each GraphSAGE operator are concatenated (see Figure 1-C for the blocks of boxes marked as “concat1”) then passed to a DiffPool layer (see Figure 1-C for the red box marked as “diff-pool1”), which finalizes the first coarsening operation. The second coarsening operation is applied in the same way as the first (as represented by “GNN pool1”, “GNN embed2”, “concat2” in Figure 1-C), and the outputs from the second DiffPool layer (“diff-pool2” in Figure 1-C) are passed to the final (fifth) GNN module (“GNN embed3” in Figure 1-C). The nodes in a graph are dynamically clustered and reduced after each coarsening operation. The coarsening ratio at each operation is determined by a pre-set DiffPool pooling ratio. Let *N*_coarsened_ represent the number of nodes in the coarsened graph and *N*_original_ the number of nodes in the original graph. The DiffPool pooling ratio *ρ*_pool_ is given by 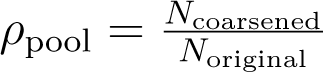. Throughout the study, we used a manually optimized value *ρ*_pool_ = 0.25. This is a manually optimized hyper-parameter.

After the final GNN module, the outputs are concatenated (“concat3” in Figure 1-C) and transformed by a global mean pooling operation (red ball “M” in Figure 1-C) to create a final graph representation. This graph representation is passed to a readout layer group (“readout” as represented by light blue boxes in Figure 1-C) consisting of two linear layers to perform graph-level regression which ultimately outputs a vector of *n* predicted parameters (“pred” as represented by a purple box in Figure 1-C). Only the first linear layer is followed by GELU (see the orange band of the first linear layer). All the linear layers incorporate dropout operations with a pre-set dropout ratio to prevent over-fitting and to utilize as many neuron connections as possible. Let *ρ*_dropout_ represent the probability *p* of disabling a connection between an input node and a hidden node of a linear layer in each epoch. The dropout ratio *ρ*_dropout_ is simply given by *ρ*_dropout_ = *p*. Throughout the study, we used a commonly picked value *ρ*_dropout_ = 0.5. This is a manually optimized hyper-parameter.

DNN’s major component is a stack comprises 5 linear layers (“DNN stack” in Figure 1-A), each followed by a BatchNorm1D (not shown in figure) and a GELU (the orange band within the boxes). All the linear layers within the stack incorporate dropout operations with *ρ*_dropout_ = 0.5. Learned features from all the linear layers within the stacks are collected and concatenated (“concat” in Figure 1-A). A single linear readout layer (“readout” in Figure 1-A) outputs *n* predicted parameters (“pred” in Figure 1-A). According to our experiments, stacking more linear layers gives no substantial improvement to the performance.

LSTM’s major component is a stack of 5 LSTM recurrent neural network layers (“LSTM stack” in Figure 1-B). The final hidden state from the last recurrent neural network layer is processed by a linear layer with *ρ*_dropout_ = 0.5 accompanied by a GELU (“linear” in Figure 1-B), then passed to a single linear readout layer (“readout” in Figure 1-B) that outputs *n* predicted parameters (“pred” in Figure 1-B). According to our experiments, stacking more recurrent neural network layers provides no substantial improvement to the performance.

The hyper-parameters not mentioned are set by their default values. The dimensions of the boxes do not map to any hyper-parameter settings, they are set for the best visual effect. The values below the boxes indicate their respective number of hidden neurons, their input and output neurons are not shown in the figure, they can be found in the configuration files in our GitHub repository “eveGNN” (Qin, 2023).

### D. Ensemble Learning

With bagging, we trained GNN, DNN and LSTM independently (“GNN”, “DNN” and “LSTM” blocks of boxes in Figure 3-Bagging), translated their outputs to parameter predictions through their own readout layers (three “readout” boxes next to the neural networks and three “pred” boxes next to the readout layers in Figure 3-Bagging) and then aggregated the predictions (red ball “A” in Figure 3-Bagging). We experimented with four aggregation methods: taking the mean, median, max and min values among the three predictions. We also recorded the individual predictions without aggregation.

With stacking, we trained GNN, DNN and LSTM simultaneously (“GNN”, “DNN” and “LSTM” blocks of boxes in Figure 3-Stacking) but without their own readout layers. We combined the features from DNN, the LSTM’s final hidden state, and GNN’s graph representation and fed to a meta-learner (“meta-learner” in Figure 3-Stacking) comprising linear neural network layers that learns to best readout parameter predictions from these combined outputs.

With boosting, there can be different pathways. In our illustration, GNN, DNN and LSTM were trained sequentially to iteratively correct residuals. For example, firstly, the GNN (“GNN” in Figure 3-Boosting) is trained from the graphs to make the initial predictions (“readout” and then “pred0” in Figure 3-Boosting) and from predicted and ground truth values of the parameters we computed the residuals (“res1” in Figure 3-Boosting)secondly, the DNN (“DNN” in Figure 3-Boosting) is trained to predict these residuals from the summary statistics (“readout” and then “pred-res1” in Figure 3-Boosting), learning to correct the GNN’s errorslastly, the LSTM (“LSTM” in Figure 3-Boosting) is trained to predict the residuals of the residuals (“readout” and then “pred-res2” in Figure 3-Boosting), which is the initial predictions minus the predicted residuals by the DNN, from branching times, to further improve the predictive accuracy. Finally, we subtracted the two residual terms from the initial predictions (red ball “S” in Figure 3-Boosting) to make the corrected predictions.

### E. Comparison between MLE Optimizers

On the phylogenies from the diversity-dependent diversification (DDD) dataset, we compared between three approaches: “Simplex”, “Subplex” and “DEoptim”. Simplex is a derivative-free optimization method that uses a simplex of solutions to iteratively explore and adjust within the parameter space, suitable for non-smooth objective functions but potentially slow for high-dimensional problems (Morgan and Deming, 1974). Subplex is an enhancement of the Simplex method, Subplex breaks high-dimensional optimization into smaller subproblems, each optimized using Simplex techniques, providing improved efficiency and effectiveness in complex parameter landscapes (Rowan, 1990). DEoptim (Differential Evolution) is a more recent population-based algorithm that applies evolutionary strategies such as mutation, crossover, and selection to efficiently navigate and optimize multimodal and complex objective functions (Ardia et al., 2010).

All three MLE methods encountered consistent optimization challenges, likely due to numerical issues related to machine precision limits or unexpected negative values during matrix operations. From a random sample of 2000 DDD phylogenies, the completion rates for each method were as follows: Simplex achieved 1966 completions from true parameter starts and 1910 from random starts; Subplex completed 1681 from true starts and 1612 from random starts; DEoptim finished 1122 from true starts and 999 from random starts. It is more difficult to estimate parameters from random starts, comparing to true starts.

For all the three optimization approaches, −1 will be returned as a parameter estimation if the likelihood becomes too small in the searching process. This means that the algorithm cannot find optima given the initial starting point of the parameters. It is highly possible that the unfinished estimations consisted of inaccurate or even −1 values. The comparison between MLE optimizers can be skewed due to less completion rate of the Subplex and DEoptim results.

In instances where optimization starting points were randomly set, a significant number of outcomes were trapped at local optima, failing to achieve global optima and often leading to inaccurate parameter estimates. This issue was less prevalent when starting points were the true parameters. For visual reference, see Figure 11 and Figure 12. Notably, in DEoptim’s best-case scenarios, estimation accuracy deteriorated significantly on larger phylogenies, as shown in the last row of Figure 12.

**Fig. 11.**
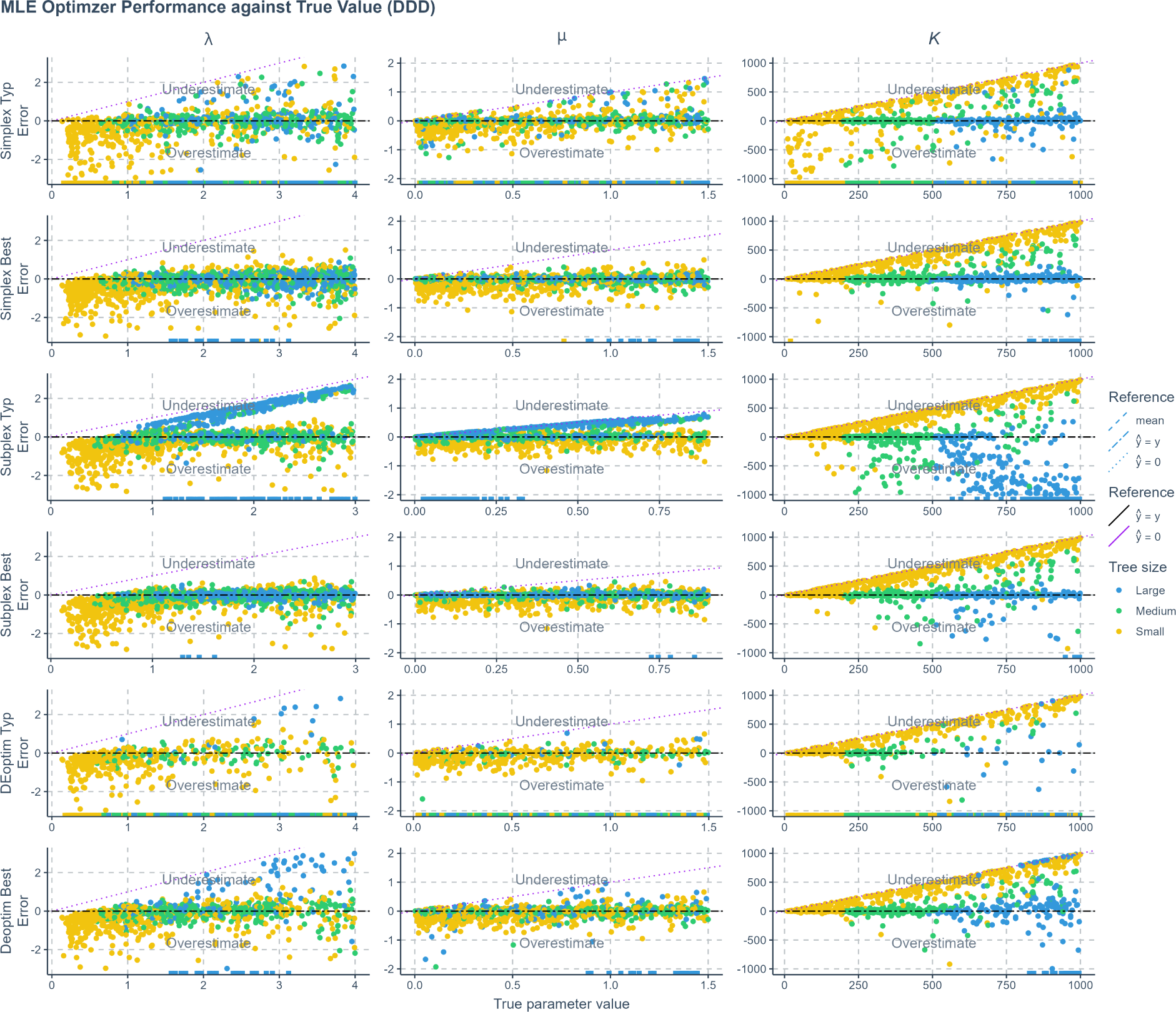
Error of maximum likelihood estimation using Simplex, Subplex and DEoptim optimzers applied to phylogenies simulated under a diversity-dependent diversification scenario, against true values. For each optimzer there were two cases. The typical case (Typ) refers to using the true parameter values as the starting points for the searching process; the best case (Best) refers to using randomly sampled values from the true parameter space as the starting points. The errors shown (y-axis) are the differences between the true parameters (x-axis) used to simulate the phylogenies and the values estimated by each method. Each row represents a method, and each column corresponds to the results for one specific parameter. Phylogenies are categorized based on their size: yellow for small phylogenies with fewer than 200 nodes (including root, internal, and tip nodes), green for medium-sized phylogenies with 200 to 500 nodes, and blue for large phylogenies with more than 500 nodes. Data points close to purple dotted lines (*ŷ* = 0) in MLE result panels indicate near-zero estimates. Black two-dash lines indicate accurate estimates (*ŷ* = *y* where *y* denotes the true parameter value). Small squares spreading along the x-axis signify optimization failures. Extremely deviating estimates are not shown in the figure. *λ*: Speciation rate. *µ*: extinction rate. *K*: carrying capacity.

**Fig. 12.**
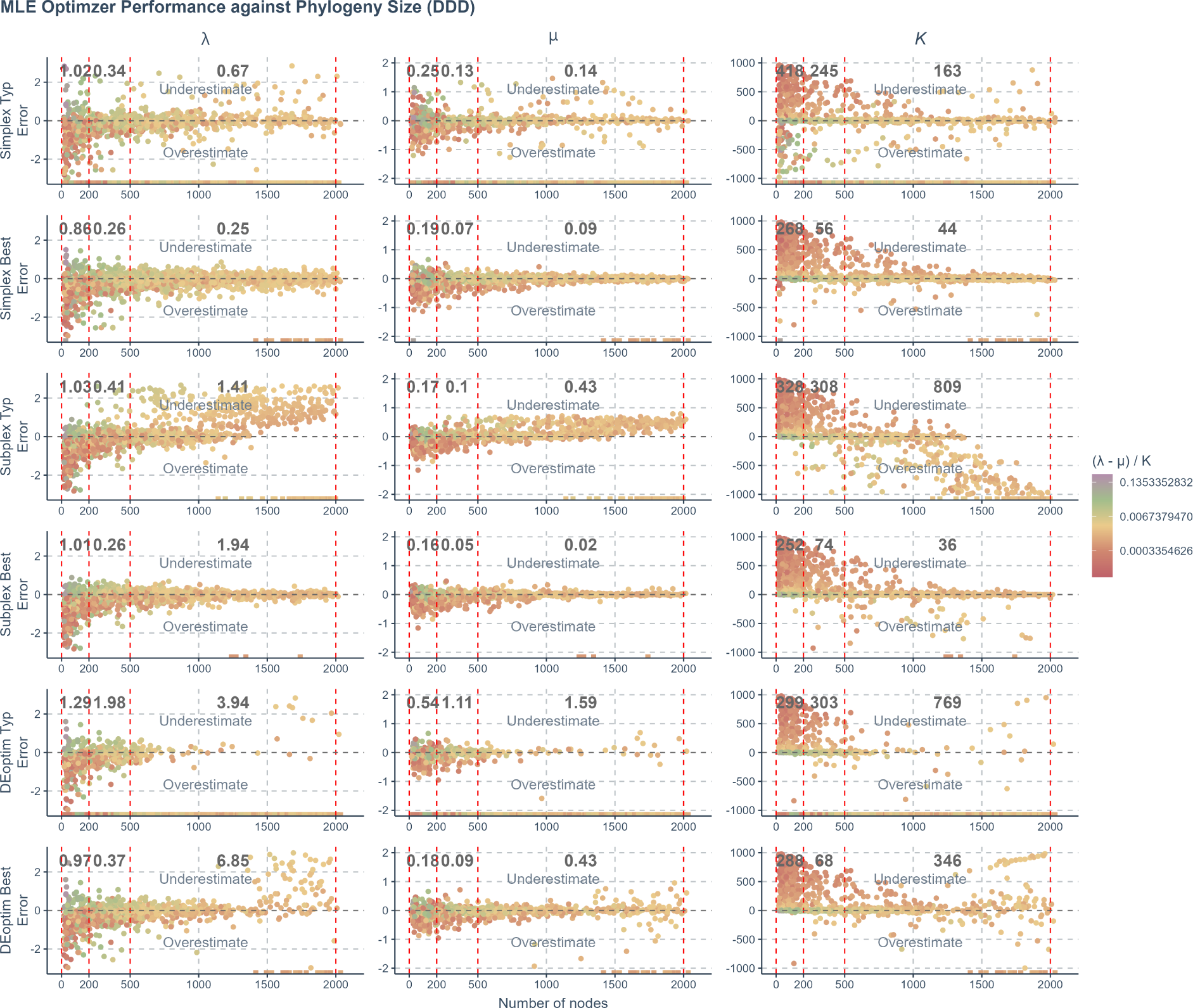
Error of maximum likelihood estimation using Simplex, Subplex and DEoptim optimzers applied to phylogenies simulated under a diversity-dependent diversification scenario, against the total number of nodes (including root, internal, and tip nodes) in the phylogenies. For each optimzer there were two cases. The typical case (Typ) refers to using the true parameter values as the starting points for the searching process; the best case (Best) refers to using randomly sampled values from the true parameter space as the starting points. The errors shown (y-axis) are the differences between the true parameters used to simulate the phylogenies and the values estimated by each method. Each row represents a method, and each column corresponds to the results for one specific parameter. Phylogenies are categorized based on their size into three sectors within each panel, separated by four vertical red dashed lines. From left to right, the sectors are: small phylogenies with fewer than 200 nodes, medium-sized phylogenies with 200 to 500 nodes, and large phylogenies with more than 500 nodes. The values shown in black within each sector are the mean absolute prediction errors of all data points in the sectors. Color coding: The color of the data points illustrates the strength of the carrying capacity effect, calculated as (*λ − µ*)*/K*. The color gradient transitions from red to purple, indicating increasing strength of the effect. This scale is transformed using log10 for clearer visual differentiation. Small squares spreading along the x-axis signify optimization failures. Extremely deviating estimates are not shown in the figure. X-axis: Size of the phylogenies. Y-axis: Error. *λ*: Speciation rate. *µ*: extinction rate. *K*: carrying capacity.

In the best cases, all the MLE optimizers are more likely to give accurate estimations on larger phylogenies while in the typical cases, larger phylogenies become a burden. Nevertheless, the MLE optimizers generally perform better on larger trees. All the MLE results showed similar trends of bias. We calculated the strength of the carrying capacity effect using the formula (*λ − µ*)*/K*, where *λ* represents the true speciation rate, *µ* the true extinction rate, and *K* the true carrying capacity. Phylogenies exhibiting a stronger carrying capacity effect typically link to more accurate estimates.

In the best-case scenarios, all MLE methods tended to yield more accurate estimates on larger phylogenies, while in typical cases, larger phylogenies posed challenges. However, all MLE methods generally performed better with larger trees, and all displayed similar trends of bias. We calculated the strength of the carrying capacity effect with the formula (*λ − µ*)*/K*, where *λ* is the true speciation rate, *µ* is the true extinction rate, and *K* is the true carrying capacity.

Subplex was the fastest among the tested algorithms, Simplex and DEoptim were slower. Simplex, although slower, completed the most computations and did not show a definitive performance disadvantage compared to Subplex or DEoptim. For this reason, in all comparisons between MLE and neural network methods, we consistently used results from the Simplex optimizer due to data coverage and reliability.

### F. Estimation Uncertainty for Empirical Trees

The rest of the figures of neural network estimation uncertainty on the empirical phylogenies under the DDD scenario can be found at https://github.com/EvoLandEco/eveGNN/tree/master/uncertainty

**Fig. 13.**
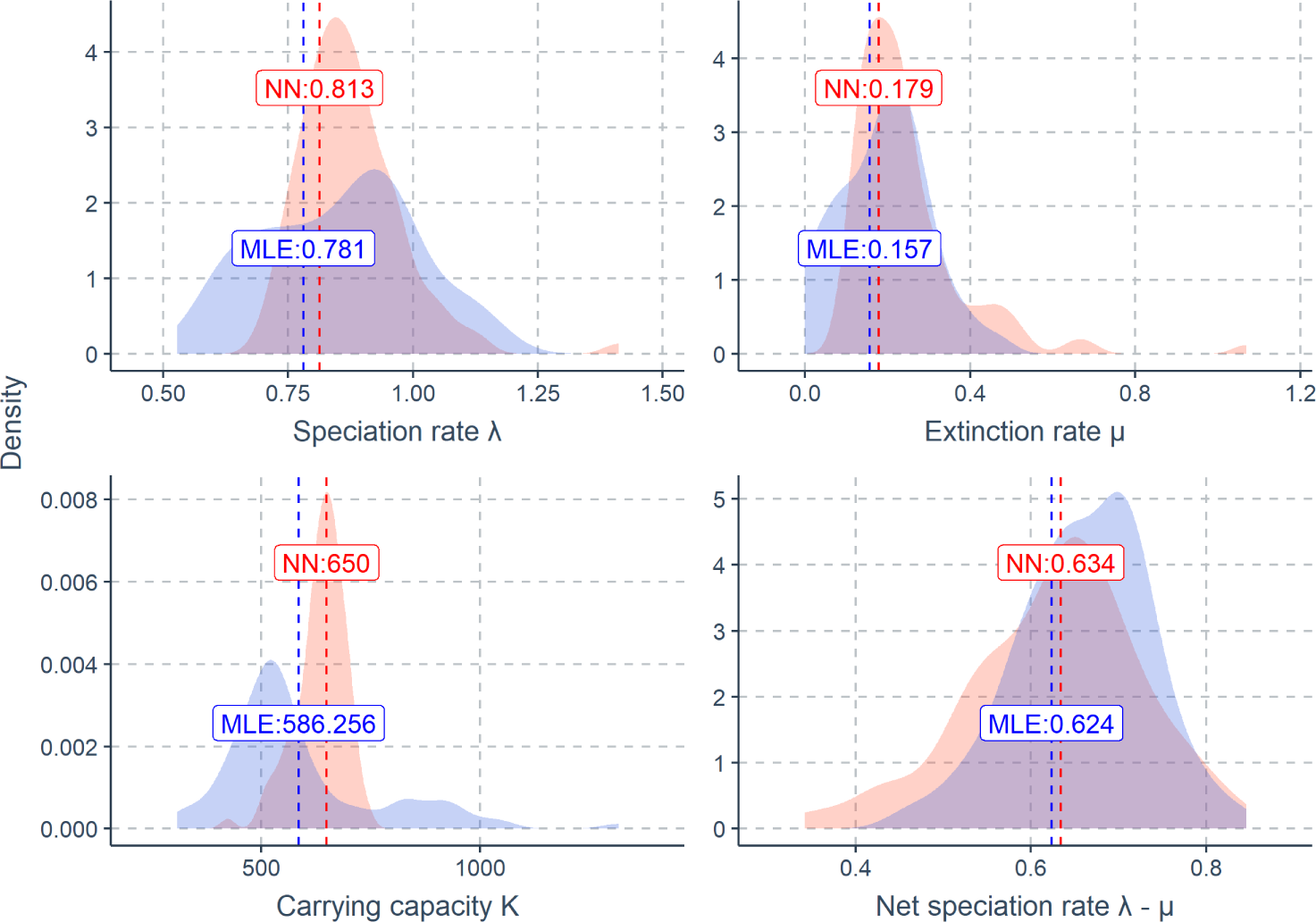
The neural network estimation uncertainty for a bird phylogeny (Furnariidae). The parameters are estimated using a pre-trained neural network (Boost BT, boosting strategy that corrects GNN results using LSTM) under a diversity-dependent diversification scenario. For reference, maximum likelihood estimation (MLE) is also used to estimate the same parameters. Each panel shows one parameter’s estimates using neural network and MLE methods with their uncertainties. The red dashed lines with red numbers indicate the estimates by the neural network method. The blue dashed lines with blue numbers indicate the estimates by the MLE method. Each pink area indicates the density distribution of a neural network estimate from 1000 bootstrap-simulated phylogenies, showing the uncertainty of neural network. Each blue area indicates the density distribution of an MLE estimate from the same set of simulated phylogenies, showing the uncertainty of MLE. X-axis: Parameter (Estimate) values. Y-axis: Density. *λ*: Speciation rate. *µ*: Extinction rate. *K*: Carrying capacity. *λ − µ*: Net speciation rate.

### G. Results under the Diversity-Dependent Diversification Scenario

**Fig. 14.**
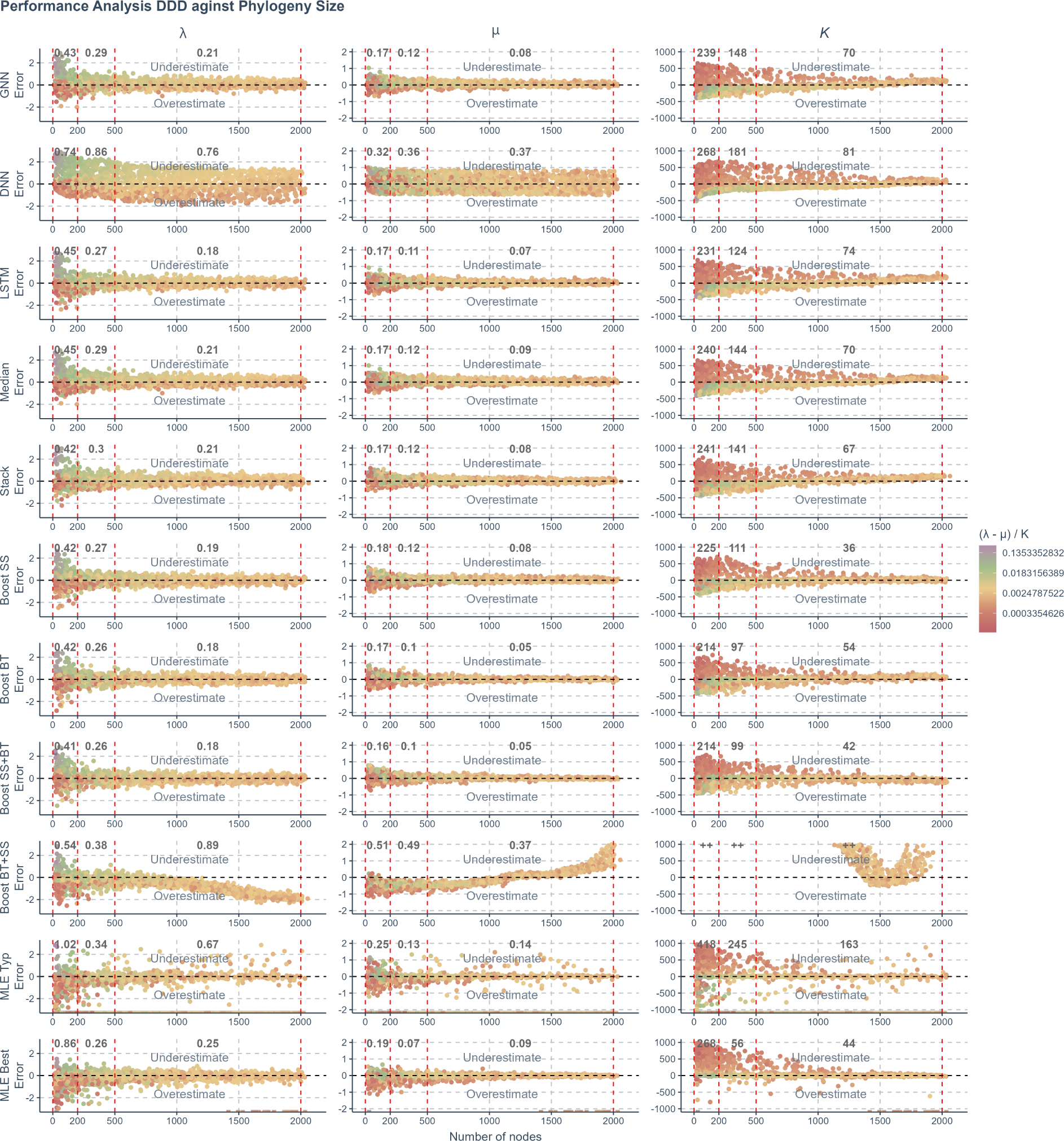
The prediction error of various methods applied to phylogenies simulated under a diversity-dependent diversification scenario, against the total number of nodes (including root, internal, and tip nodes) in the phylogenies. The errors shown are the differences between the true parameters used to simulate the phylogenies and the values predicted or estimated by each method. Each row represents a method, and each column corresponds to the results for one specific parameter. Phylogenies are categorized based on their size into three sectors within each panel, separated by four vertical red dashed lines. From left to right, the sectors are: small phylogenies with fewer than 200 nodes, medium-sized phylogenies with 200 to 500 nodes, and large phylogenies with more than 500 nodes. The values shown in black within each sector are the mean absolute prediction errors of all data points in the sectors. Color coding: The color of the data points illustrates the strength of the carrying capacity effect, calculated as (*λ − µ*)*/K*. The color gradient transitions from red to purple, indicating increasing strength of the effect. This scale is transformed using log10 for clearer visual differentiation. GNN: Predictions obtained by the graph neural network using the phylogenies transformed to graph format. DNN: Predictions by the dense neural network using summary statistics. LSTM: Predictions by the long short-term memory recurrent neural network using branching times. Median: Bagging strategy that takes the median value of the predictions from GNN, DNN, and LSTM. Stack: Stacking strategy that utilizes a meta-learner to integrate results from GNN, DNN, and LSTM. Boost SS: Boosting strategy that corrects GNN results using DNN. Boost BT: Boosting strategy that corrects GNN results using LSTM. Boost SS+BT: Sequential correction of GNN errors first using DNN, followed by LSTM. Boost BT+SS: Sequential correction of GNN errors first using LSTM, followed by DNN. MLE Typ: Maximum Likelihood Estimation results using random starting points for parameter optimization. MLE Best: MLE results using the true parameter values as the starting points for optimization. In the MLE result panels, small squares spreading along the x-axis signify optimization failures. Due to significantly lower accuracy, other aggregation methods from the bagging strategy are not displayed on the plot. X-axis: Size of the phylogenies. Y-axis: Error. *λ*: Speciation rate. *µ*: Extinction rate. *K*: Carrying capacity.

**Fig. 15.**
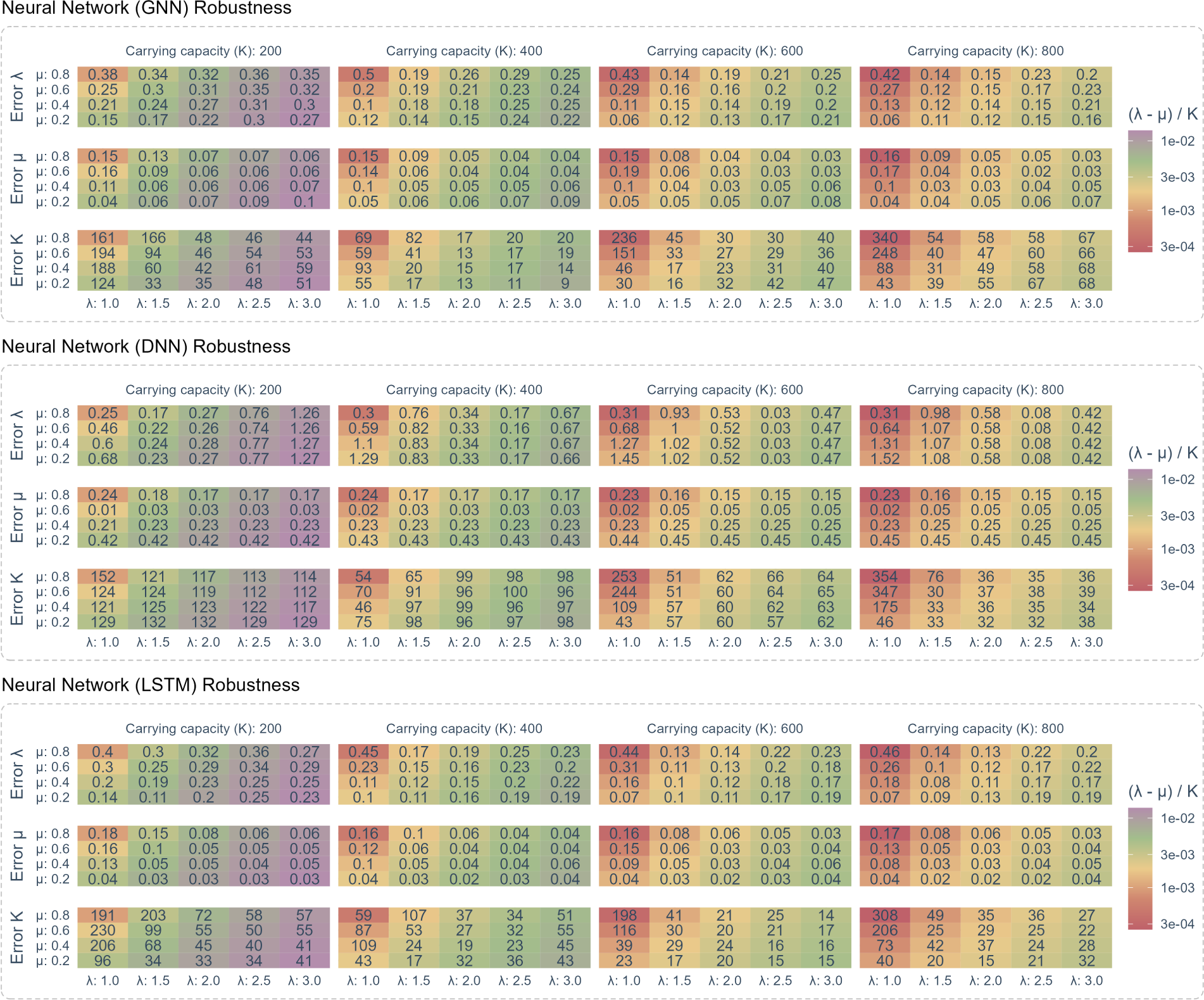
Robustness comparison between graph neural network (GNN), dense neural network (DNN) and long short-term memory recurrent neural network (LSTM) when operating independently. Same structure as the previous figure.

**Fig. 16.**
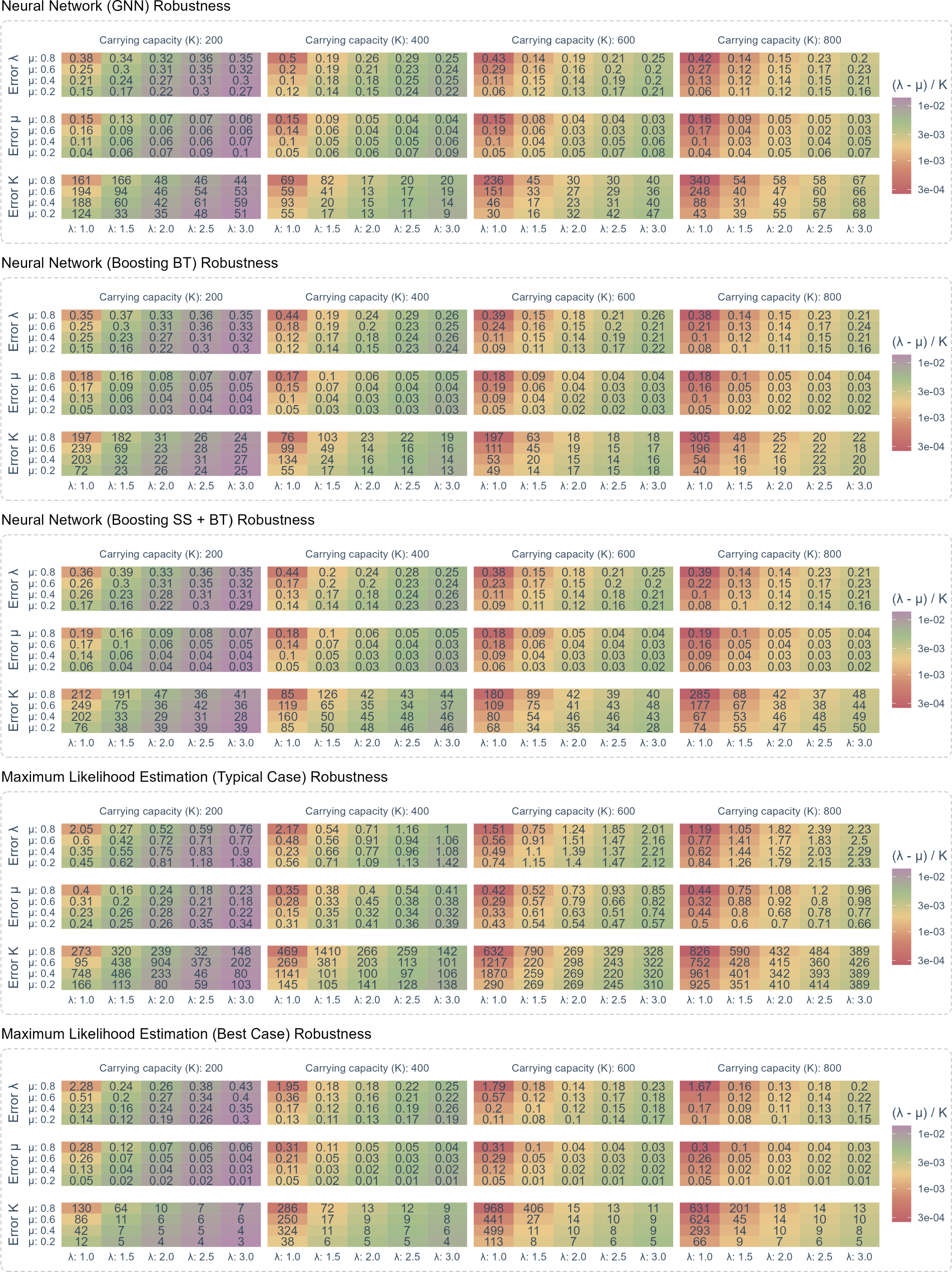
The robustness of neural network and maximum likelihood estimation was assessed on 80 sets of phylogenies, each containing 1000 trees randomly simulated under a diversity-dependent diversification scenario, employing identical parameter settings but varied in size, topology, and structure. Each segment delineated by dashed lines corresponds to distinct methods. Each column within a segment is associated with a specific carrying capacity (*K*) used in the simulation of the phylogenies. Each row within a segment details the mean absolute errors between the true and estimated values of a specific parameter, with parameter names labeled on the left side of each row. GNN: Graph neural network. Boosting BT: Graph neural network with long short-term memory recurrent neural network correcting its residuals using branching times. Boosting SS + BT: Graph neural network with dense neural network and long short-term memory recurrent neural network correcting residuals sequentially using summary statistics and branching times. Typical Case: Maximum likelihood estimation using random initial parameter as the starting point. Best Case: Maximum likelihood estimation using true parameter as the starting point. X-axis: Represents the true speciation rate (*λ*) used to simulate phylogenies. Y-axis: Represents the true extinction rate (*µ*) used to simulate phylogenies. Cell Content: The numbers displayed within each heatmap cell indicate the mean absolute error for a parameter, given the specific *λ*, *µ* and *K* settings. Color Coding: The background color of each cell illustrates the strength of the carrying capacity effect, calculated as (*λ − µ*)*/K*. The color gradient transitions from red to purple, indicating increasing strength of the effect. This scale is transformed using log10 for clearer visual differentiation. Note that the numerical values within the cells are not mapped to the background colors. For a detailed reference to the effect strength values corresponding to the background colors, refer to the figure legends.

**Fig. 17.**
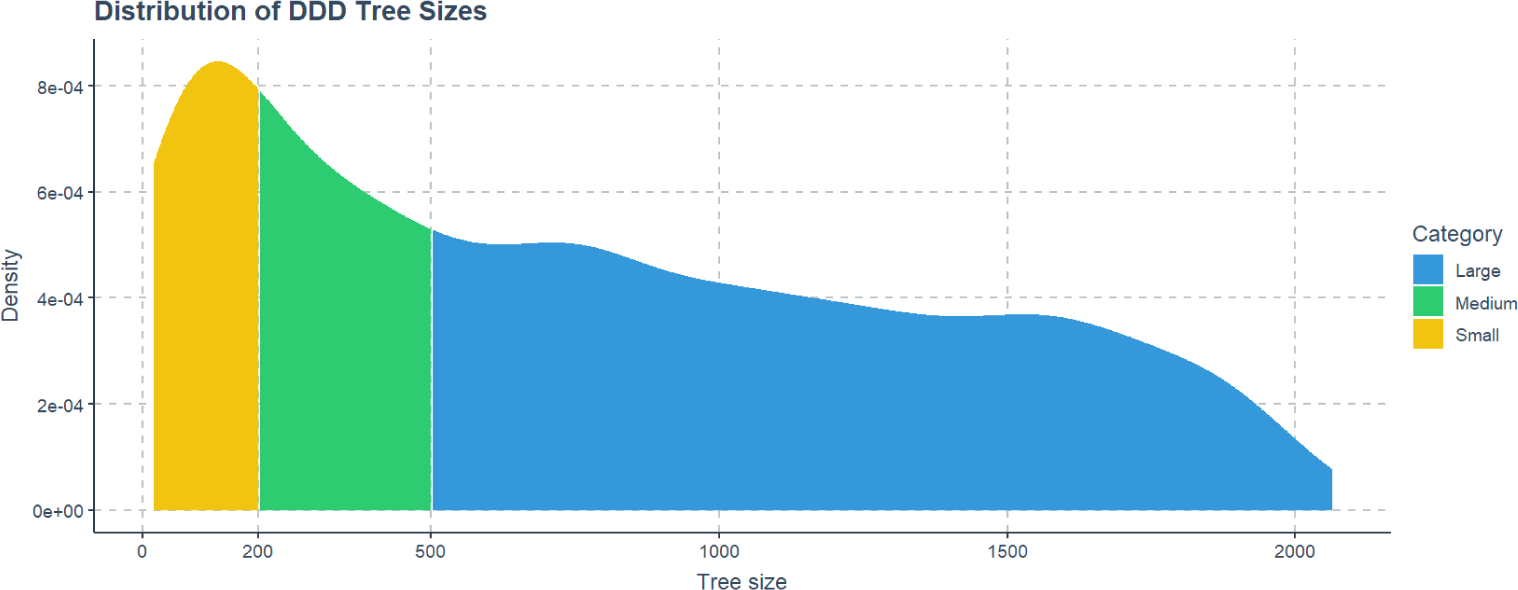
The density distribution of phylogeny sizes under the diversity-dependent diversification (DDD) scenario. The colors of the areas under the density curve indicate the three categories used in our analyses. Yellow area: Small-sized phylogenies with less than 200 nodes (approx. 100 tips). Greed area: Medium-sized phylogenies with more than 200 nodes and less than 500 nodes (approx. 250 tips). Blue area: Large-sized phylogenies with more than 500 nodes.

### H. Results under the Birth-Death Scenario

**Fig. 18.**
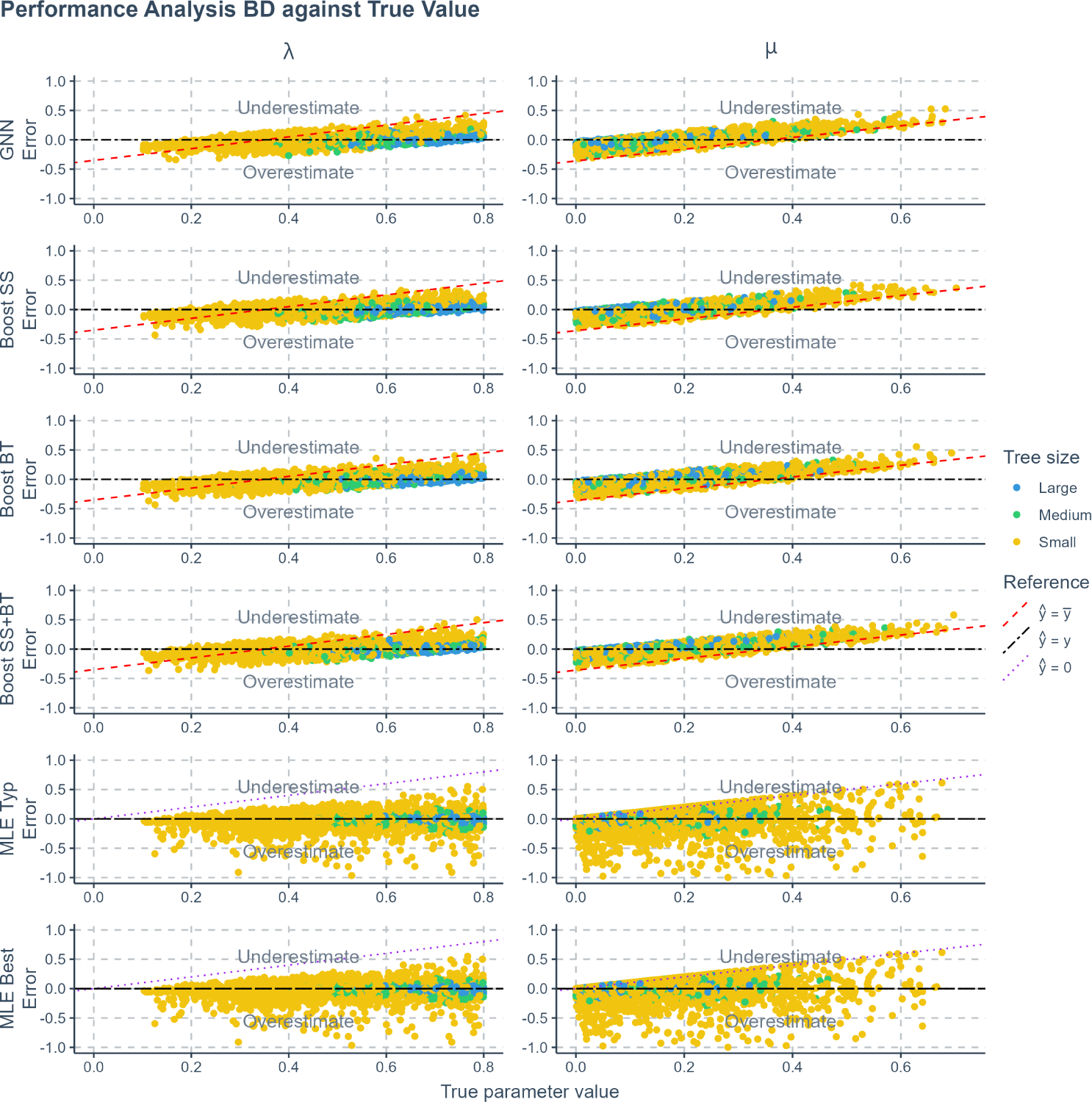
The prediction error (absolute error) of various methods applied to phylogenies simulated under a birth-death scenario, against true values. The errors shown are the differences between the true parameters used to simulate the phylogenies and the values predicted or estimated by each method. Each row represents a method, and each column corresponds to the results for one specific parameter. Phylogenies are categorized based on their size: yellow for small phylogenies with fewer than 200 nodes (including root, internal, and tip nodes), green for medium-sized phylogenies with 200 to 500 nodes, and blue for large phylogenies with more than 500 nodes. GNN: Predictions obtained by the graph neural network using the phylogenies. Boost SS: Boosting strategy that corrects GNN results using DNN. Boost BT: Boosting strategy that corrects GNN results using LSTM. Boost SS+BT: Sequential correction of GNN errors first using DNN, followed by LSTM. MLE Typ: Maximum Likelihood Estimation results using random starting points for parameter optimization. MLE Best: MLE results using the true parameter values as the starting points for optimization. Red dashed lines in panels representing neural network results indicate the mid-points of the parameter spaces (*ŷ* = *ȳ* where *ŷ* denotes a estimated parameter and *ȳ* denotes the mid-point of the parameter space). Data points close to purple dotted lines (*ŷ* = 0) in MLE result panels indicate near-zero estimates. Black two-dash lines indicate accurate estimates (*ŷ* = *y* where *y* denotes the true parameter value). In the MLE result panels, small squares spreading along the x-axis signify optimization failures. X-axis: True parameter values. Y-axis: Error, or difference between true and predicted values. *λ*: Speciation rate. *µ*: Extinction rate.

**Fig. 19.**
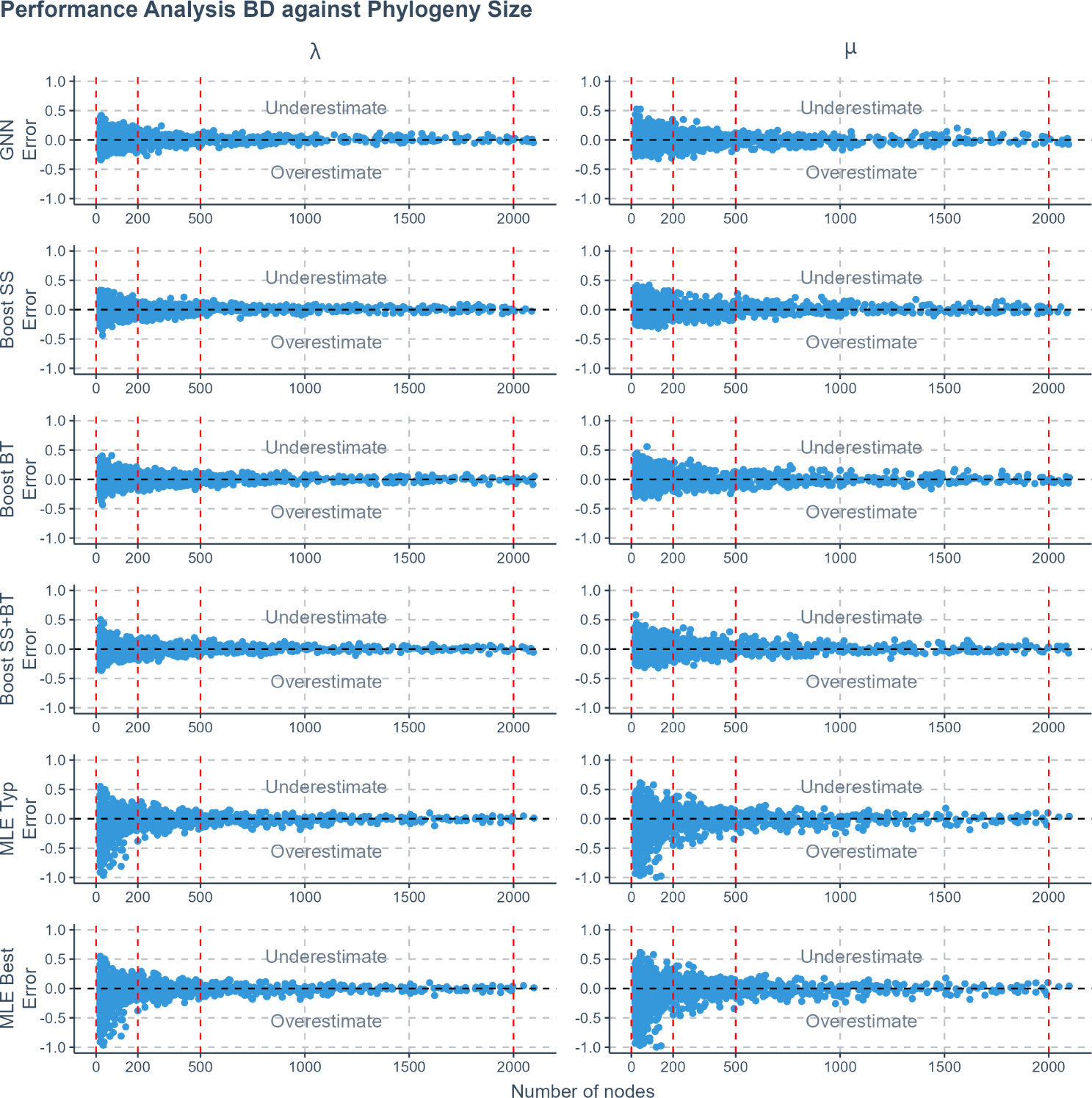
The prediction error (absolute error) of various methods applied to phylogenies simulated under a birth-death scenario, against the total number of nodes in the phylogenies. The errors shown are the differences between the true parameters used to simulate the phylogenies and the values predicted or estimated by each method. Each row represents a method, and each column corresponds to the results for one specific parameter. Phylogenies are categorized based on their size into three sectors within each panel, separated by four vertical red dashed lines. From left to right, the sectors are: small phylogenies with fewer than 200 nodes (including root, internal, and tip nodes), medium-sized phylogenies with 200 to 500 nodes, and large phylogenies with more than 500 nodes. GNN: Predictions obtained by the graph neural network using the phylogenies. Boost SS: Boosting strategy that corrects GNN results using DNN. Boost BT: Boosting strategy that corrects GNN results using LSTM. Boost SS+BT: Sequential correction of GNN errors first using DNN, followed by LSTM. MLE Typ: Maximum Likelihood Estimation results using random starting points for parameter optimization. MLE Best: MLE results using the true parameter values as the starting points for optimization. X-axis: Size of the phylogenies. Y-axis: Error. *λ*: Speciation rate. *µ*: Extinction rate.

### I. Results under the Protracted Birth-Death Scenario

**Fig. 20.**
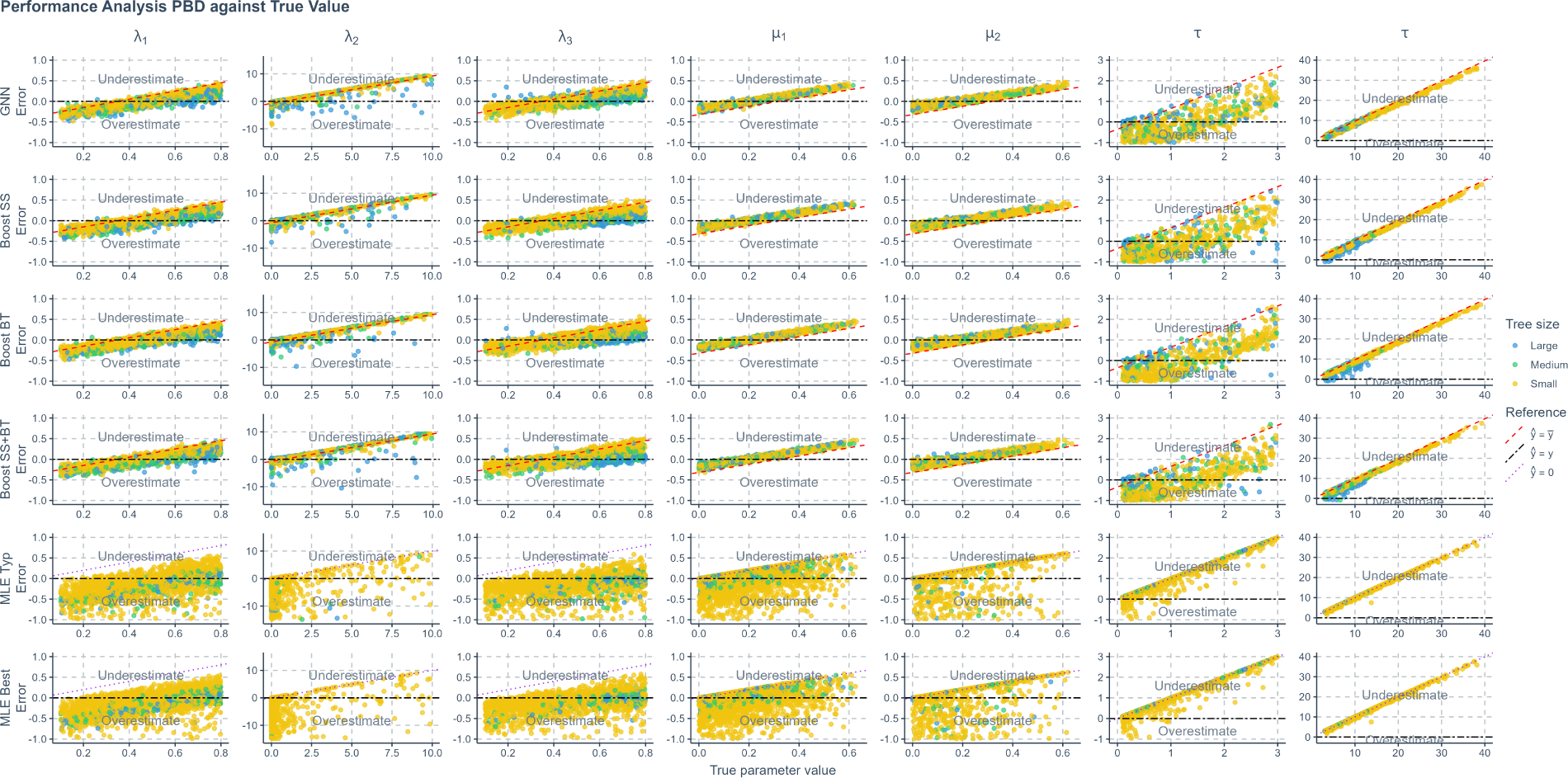
The prediction error (absolute error) of various methods applied to phylogenies simulated under a protracted birth-death scenario, against true values. The errors shown are the differences between the true parameters used to simulate the phylogenies and the values predicted or estimated by each method. Each row represents a method, and each column corresponds to the results for one specific parameter. Phylogenies are categorized based on their size: yellow for small phylogenies with fewer than 200 nodes (including root, internal, and tip nodes), green for medium-sized phylogenies with 200 to 500 nodes, and blue for large phylogenies with more than 500 nodes. GNN: Predictions obtained by the graph neural network using the phylogenies. Boost SS: Boosting strategy that corrects GNN results using DNN. Boost BT: Boosting strategy that corrects GNN results using LSTM. Boost SS+BT: Sequential correction of GNN errors first using DNN, followed by LSTM. MLE Typ: Maximum Likelihood Estimation results using random starting points for parameter optimization. MLE Best: MLE results using the true parameter values as the starting points for optimization. Red dashed lines in panels representing neural network results indicate the mid-points of the parameter spaces. Purple dotted lines in MLE result panels signify where estimated values are 0. X-axis: True parameter values. Y-axis: Error, or difference between true and predicted values. *λ*_1_: Speciation initiation rate of the good species. *λ*_2_: Speciation completion rate. *λ*_3_: Speciation initiation rate of the incipient species. *µ*_1_: Extinction rate of the good species. *µ*_2_: Extinction rate of the incipient species. *τ* : Expected duration of speciation.

**Fig. 21.**
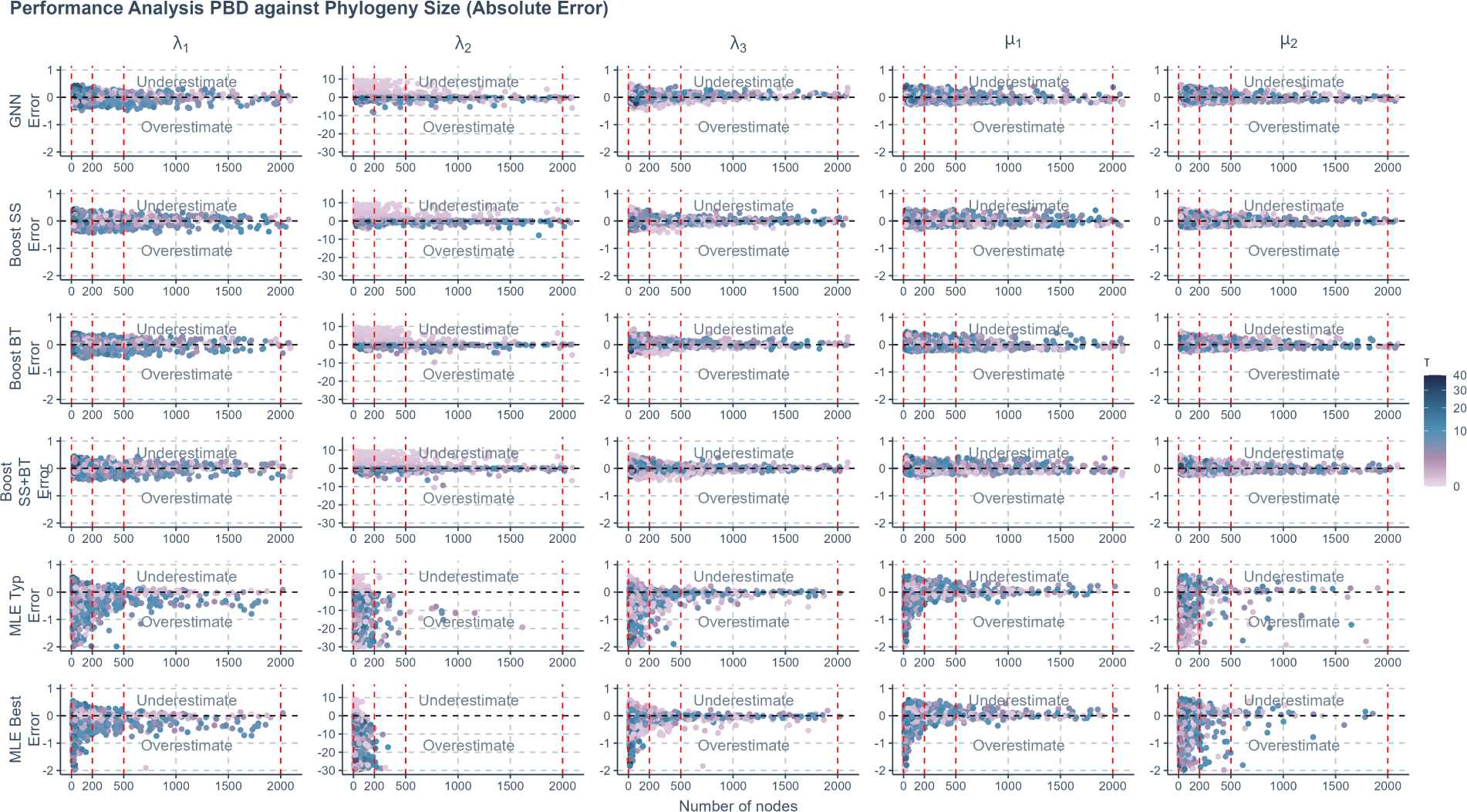
The prediction error (absolute error) of various methods applied to phylogenies simulated under a protracted birth-death scenario, against the total number of nodes in the phylogenies. The errors shown are the differences between the true parameters used to simulate the phylogenies and the values predicted or estimated by each method. Each row represents a method, and each column corresponds to the results for one specific parameter. Phylogenies are categorized based on their size into three sectors within each panel, separated by four vertical red dashed lines. From left to right, the sectors are: small phylogenies with fewer than 200 nodes (including root, internal, and tip nodes), medium-sized phylogenies with 200 to 500 nodes, and large phylogenies with more than 500 nodes. Color Coding: The color of the data points illustrates the expected duration of speciation. The color gradient transitions from light purple to dark blue, indicating increasing value of the duration. This scale is transformed using square root for clearer visual differentiation. GNN: Predictions obtained by the graph neural network using the phylogenies. Boost SS: Boosting strategy that corrects GNN results using DNN. Boost BT: Boosting strategy that corrects GNN results using LSTM. Boost SS+BT: Sequential correction of GNN errors first using DNN, followed by LSTM. MLE Typ: Maximum Likelihood Estimation results using random starting points for parameter optimization. MLE Best: MLE results using the true parameter values as the starting points for optimization. X-axis: Size of the phylogenies. Y-axis: Error. *λ*_1_: Speciation initiation rate of the good species. *λ*_2_: Speciation completion rate. *λ*_3_: Speciation initiation rate of the incipient species. *µ*_1_: Extinction rate of the good species. *µ*_2_: Extinction rate of the incipient species. *τ* : Expected duration of speciation.

**Fig. 22.**
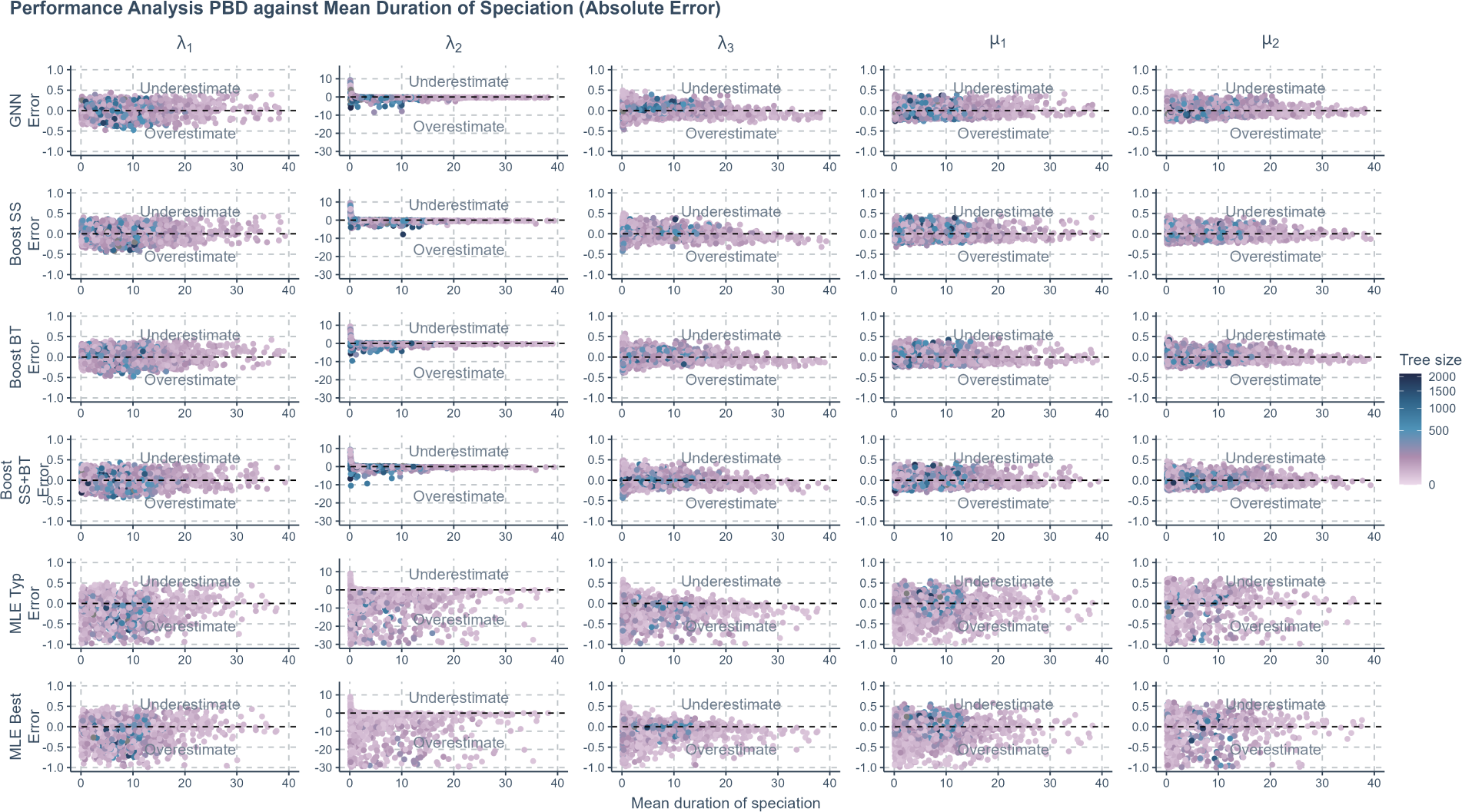
The prediction error (absolute error) of various methods applied to phylogenies simulated under a protracted birth-death scenario, against the true mean duration of speciation. The errors shown are the differences between the true parameters used to simulate the phylogenies and the values predicted or estimated by each method. Each row represents a method, and each column corresponds to the results for one specific parameter. Color Coding: The color of the data points illustrates the total number of nodes of the phylogenies. The color gradient transitions from light purple to dark blue, indicating increasing value of the node number. This scale is transformed using square root for clearer visual differentiation. GNN: Predictions obtained by the graph neural network using the phylogenies. Boost SS: Boosting strategy that corrects GNN results using DNN. Boost BT: Boosting strategy that corrects GNN results using LSTM. Boost SS+BT: Sequential correction of GNN errors first using DNN, followed by LSTM. MLE Typ: Maximum Likelihood Estimation results using random starting points for parameter optimization. MLE Best: MLE results using the true parameter values as the starting points for optimization. X-axis: Size of the phylogenies. Y-axis: Error. *λ*_1_: Speciation initiation rate of the good species. *λ*_2_: Speciation completion rate. *λ*_3_: Speciation initiation rate of the incipient species. *µ*_1_: Extinction rate of the good species. *µ*_2_: Extinction rate of the incipient species. *τ* : Expected duration of speciation.

**Fig. 23.**
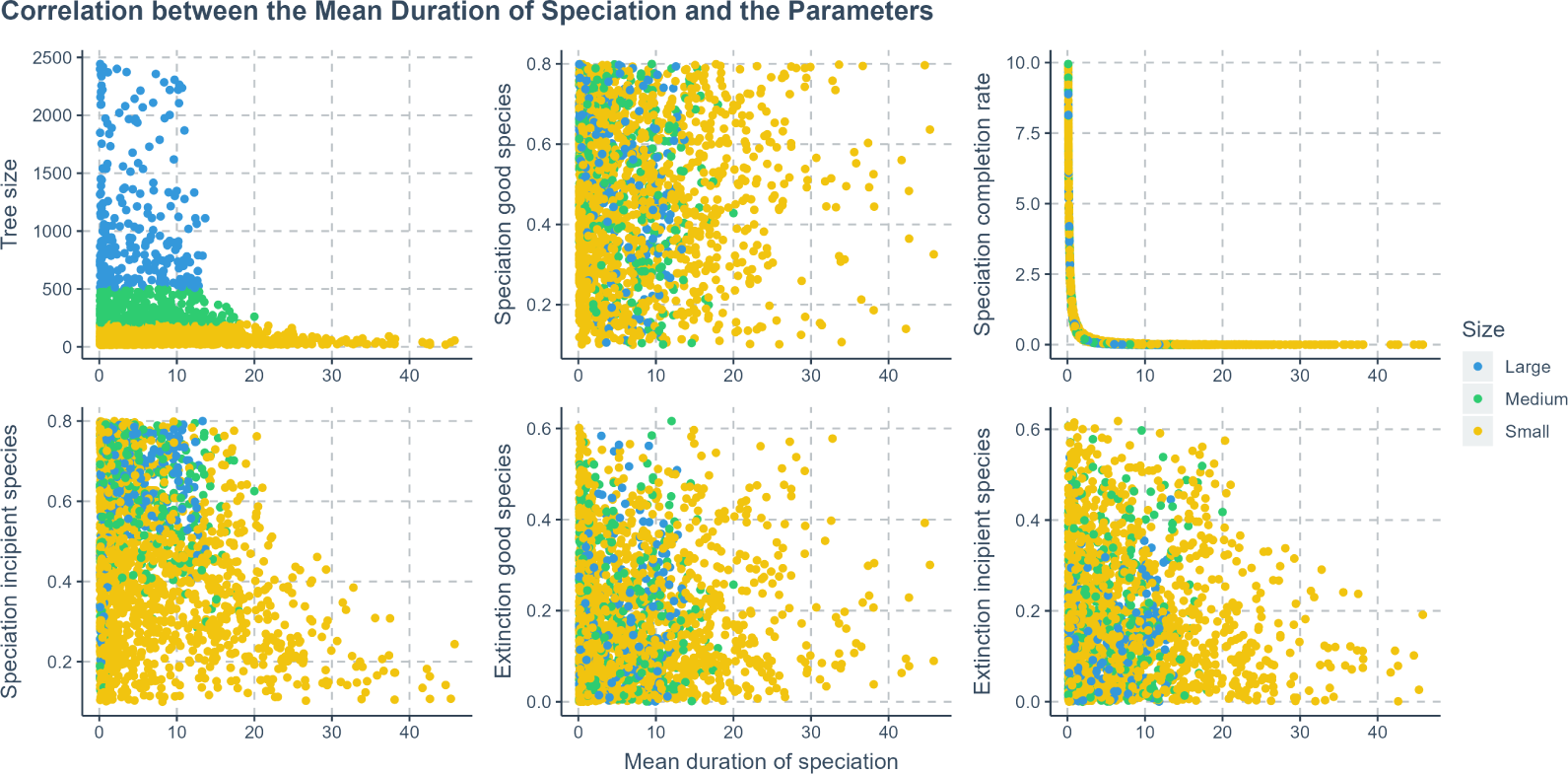
The correlation between the mean duration of speciation and the true parameter values under the protracted birth-death diversification scenario. Phylogenies are categorized based on their size: yellow for small phylogenies with fewer than 200 nodes (including root, internal, and tip nodes), green for medium-sized phylogenies with 200 to 500 nodes, and blue for large phylogenies with more than 500 nodes. X-axis: Mean duration of speciation. Y-axis: Tree size.

### J. Dataset Re-balancing

Birth-death processes without carrying capacity effect may have larger variance of tree size than the DDD trees. A skew in the frequency of tree size across datasets may appear that leads to a non-representative sample.

To address this issue, we re-balanced the BD dataset by creating 10 bins, each designated to hold phylogenies within specific size ranges, spanning from 10 to 2000 nodes in increments of 200 nodes per bin (the first bin accepts phylogeny of sizes 10 to 200). We randomly simulated phylogenies using parameters sampled from the same space as the BD training dataset and allocated them to these bins according to their sizes, continuing this process until each bin reached its target capacity of 10,000 phylogenies. This method leads to a more equal representation of phylogenies of each size range, reducing size-based sampling bias. The filled bins were subsequently combined to form a re-balanced dataset, which in total has 100,000 phylogenies.

To compare with the original BD dataset, we trained neural networks on the re-balanced dataset, and validated neural network performance on an additional testing dataset (10,000 phylogenies simulated using the same parameter space). We computed MLE estimates on 2,000 randomly sampled phylogenies from the testing dataset.

**Fig. 24.**
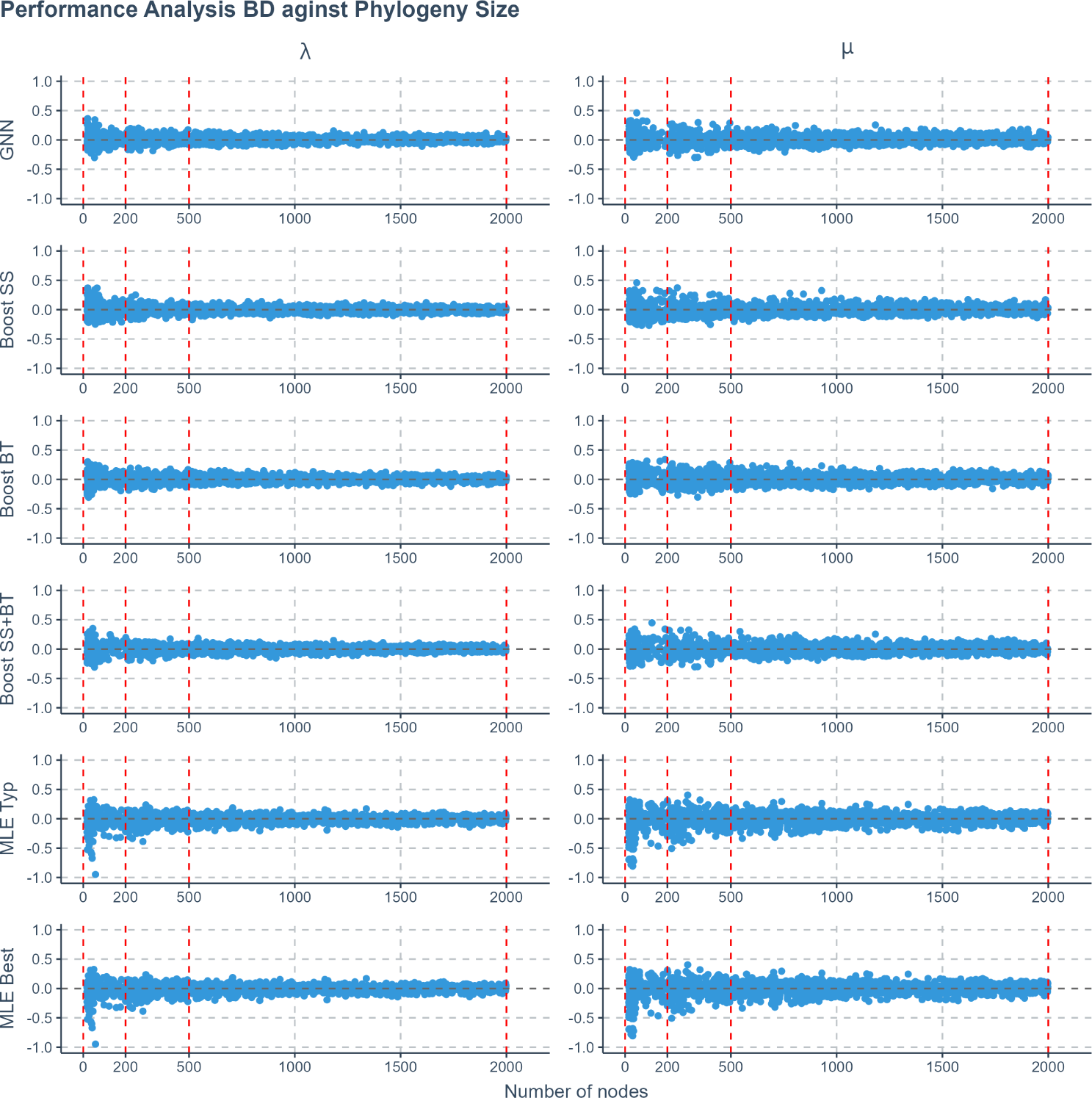
The prediction error (absolute error) of various methods applied to re-balanced phylogenies simulated under a birth-death scenario, against the total number of nodes in the phylogenies. The errors shown are the differences between the true parameters used to simulate the phylogenies and the values predicted or estimated by each method. Each row represents a method, and each column corresponds to the results for one specific parameter. Phylogenies are categorized based on their size into three sectors within each panel, separated by four vertical red dashed lines. From left to right, the sectors are: small phylogenies with fewer than 200 nodes (including root, internal, and tip nodes), medium-sized phylogenies with 200 to 500 nodes, and large phylogenies with more than 500 nodes. This scale is transformed using square root for clearer visual differentiation. GNN: Predictions obtained by the graph neural network using the phylogenies. Boost SS: Boosting strategy that corrects GNN results using DNN. Boost BT: Boosting strategy that corrects GNN results using LSTM. Boost SS+BT: Sequential correction of GNN errors first using DNN, followed by LSTM. MLE Typ: Maximum Likelihood Estimation results using random starting points for parameter optimization. MLE Best: MLE results using the true parameter values as the starting points for optimization. X-axis: Size of the phylogenies. Y-axis: Error. *λ*: Speciation rate. *µ*: Extinction rate.

### K. Unseen Data and Complete Phylogeny

We simulated additional datasets to explore the generalization ability of the neural networks when facing data with completely unseen true parameters, as well as to compare neural network performances between extant and complete phylogenies. Each simulated dataset was divided into in-sample and out-of-sample datasets. We used the in-sample datasets for training and testing and the out-of-sample datasets for evaluating the generalization ability of the trained neural networks. Validating trained neural networks on the out-of-sample datasets can provide insights into whether their performances are tailored to the peculiarities of the already seen data and whether they are robust to new, unseen phylogenies. For each tree we kept two versions: tree of all species (TAS) and tree of extant species (TES). See Table 2 for the parameter settings of the additional datasets, see Table 3 for the criteria of in-sample and out-of-sample dataset separation. To conserve GPU memory, the parameter space for additional datasets was deliberately kept smaller, given that the TAS dataset inherently contains far more information than the TES.

**Table 2.** List of simulated tree datasets. The type column specifies which function is used to generate the trees. The age column specifies the crown age of the trees. The N column specifies the number of trees in the dataset. The rest of the columns specify the lower (*a*) and the upper (*b*) bounds of the initial parameters for the tree simulations, all the parameters are sampled from *U* (*a, b*) except for *λ*_1_ of the protracted birth-death scenario. *λ*_1_ is computed as *λ*_1_ = 10*^e^* where *e* is sampled from *U* (*−*3, 1). *U* denotes uniform distribution. List A shows the parameter distributions of the birth-death trees and the diversity-dependent-diversification trees, *λ*: intrinsic speciation rate/birth rate; *µ*: intrinsic extinction rate/death rate; *K*: carrying capacity. List B shows the parameter distributions of the protracted birth-death trees, *λ*_1_: speciation-initiation rate of good species; *λ*_2_: speciation-completion rate; *λ*_3_: speciation-initiation rate of incipient species; *µ*_1_: extinction rate of good species; *µ*_2_: extinction rate of incipient species. *^∗^*In diversity-dependent-diversification simulations, the maximum extinction rate is capped at 1.5 if 0.9*λ >* 1.5.

| Type | Age | N | $\lambda_0$ | | $\mu_0$ | | $K$ | |
| --- | --- | --- | --- | --- | --- | --- | --- | --- |
| | | | $a$ | $b$ | $a$ | $b$ | $a$ | $b$ |
| BD | 10 | 60k | 0.1 | 0.6 | 0.0 | $0.9\lambda_0$ | - | - |
| DDD | 10 | 100k | 0.1 | 3.0 | 0.0 | $0.9\lambda_0^*$ | 10 | 1000 |

| Type | Age | N | $b_1$ | | $\lambda_1$ | | $b_2$ | | $\mu_1$ | | $\mu_2$ | |
| --- | --- | --- | --- | --- | --- | --- | --- | --- | --- | --- | --- | --- |
| | | | $a$ | $b$ | $a$ | $b$ | $a$ | $b$ | $a$ | $b$ | $a$ | $b$ |
| PBD | 10 | 100k | 0.1 | 0.8 | 0.001 | 10 | 0.1 | 0.8 | 0.0 | $0.8b_1$ | 0.0 | $0.8b_2$ |

**Table 3.** Criteria for in-sample and out-of-sample dataset separation. Trees generated from each model are separated into left out-of-sample group, in-sample group and right out-of-sample group, based on the parameter ranges. The Model column shows the model of a parameter; the Parameter column shows the corresponding parameter; the Left Out column shows the criteria for the left out-of-sample group; the In Sample column shows the criteria for the in-sample group; the Right Out column shows the criteria of the right out-of-sample group. *λ*: intrinsic speciation rate/birth rate; *µ*: intrinsic extinction rate/death rate; *K*: carrying capacity. List B shows the parameter distributions of the protracted birth-death trees, *λ*_1_: speciation-initiation rate of good species; *λ*_2_: speciation-completion rate; *λ*_3_: speciation-initiation rate of incipient species; *µ*_1_: extinction rate of good species; *µ*_2_: extinction rate of incipient species.

| Model | Parameter | Left Out | In Sample | Right Out |
| --- | --- | --- | --- | --- |
| BD | $\lambda_0$ | [0.10, 0.18) | [0.18, 0.52] | (0.52, 0.60] |
| BD | $\mu_0$ | [0.00, 0.08) | [0.08, 0.46] | (0.46, 0.54] |
| DDD | $\lambda_0$ | [0.00, 0.30) | [0.30, 2.70] | (2.70, 3.00] |
| DDD | $\mu_0$ | [0.00, 0.10) | [0.10, 0.80] | (0.80, 0.90] |
| DDD | $K$ | [10, 100) | [100, 900] | (900, 1000] |
| PBD | $b_1$ | [0.10, 0.18) | [0.18, 0.72] | (0.72, 0.80] |
| PBD | $\lambda_1$ | [0.001, 0.002) | [0.002, 5] | (5, 10] |
| PBD | $b_2$ | [0.10, 0.18) | [0.18, 0.72] | (0.72, 0.80] |
| PBD | $\mu_1$ | [0.00, 0.06) | [0.06, 0.58] | (0.58, 0.64] |
| PBD | $\mu_2$ | [0.00, 0.06) | [0.06, 0.58] | (0.58, 0.64] |

**Fig. 25.**
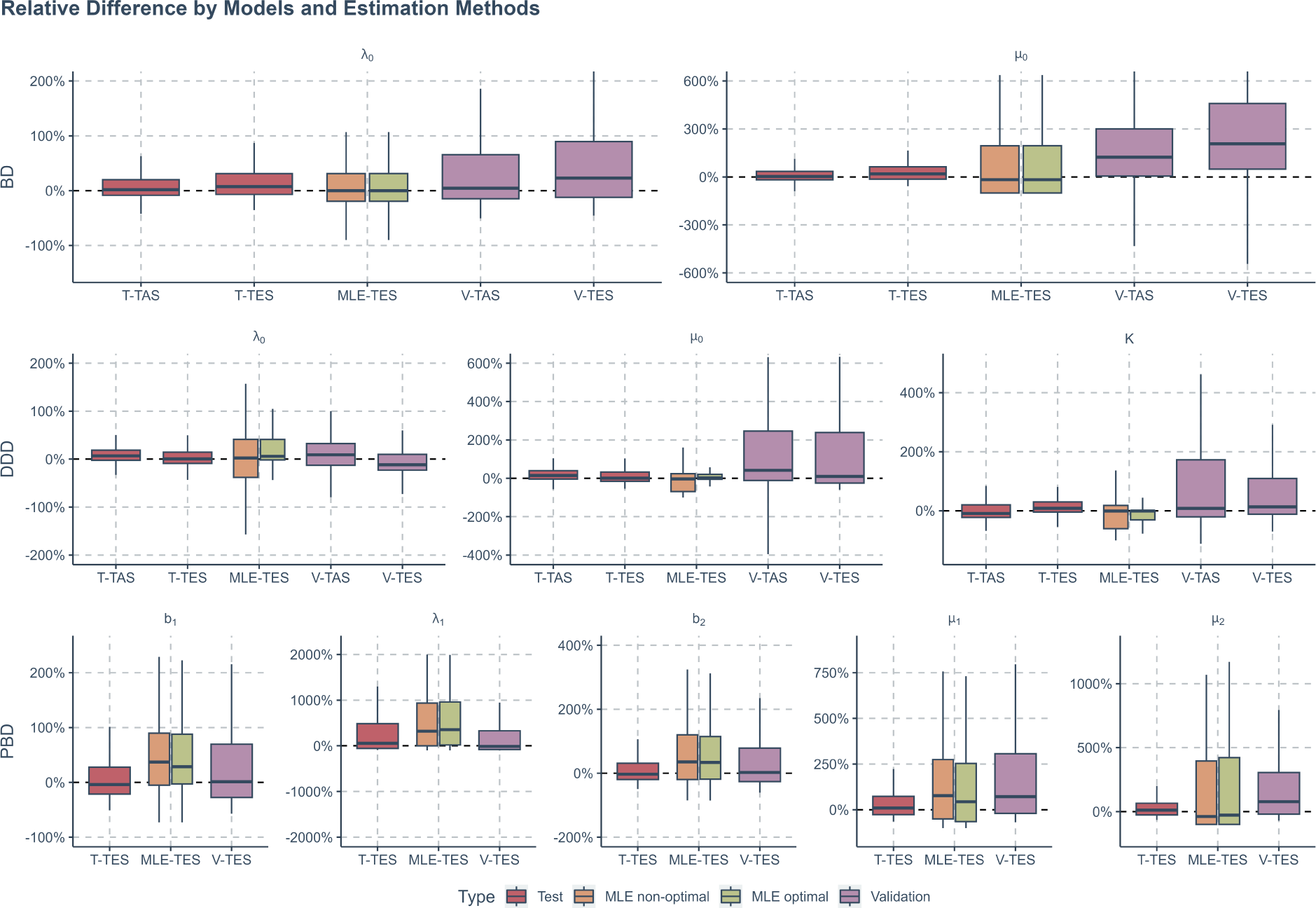
Comparisons of relative differences (in percentage) between ground true and estimated parameter values. From top to bottom, the panels in each row present the relative differences of trees generated by a specific diversification process. From left to right, the panels in each column present the relative differences of a specific parameter used when simulating the trees. Within each panel, each box represents a specific method for parameter estimation on a specific data set (as described in x-axis labels). Red boxes represent parameter estimation by using only graph neural network (GNN) on the in-sample datasets (Test in figure), yellow boxes represent the non-optimal maximum likelihood estimation (MLE) method on the complete datasets (direct outputs from simulation, without any separation), green boxes represent the optimal MLE method on the complete datasets, purple boxes represent parameter estimation by GNN on the out-of-sample datasets. BD - birth-death trees; DDD - diversity-dependent-diversification trees; PBD - protracted birth-death trees. *λ* - birth rate/intrinsic speciation rate; *µ* death rate/intrinsic extinction rate; *K* - carrying capacity; *λ*_1_ - speciation rate of good species; *λ*_2_ - speciation-completion rate; *λ*_3_ - speciation rate of incipient species; *µ*_1_ - extinction rate of good species; *µ*_2_ extinction rate of incipient species. T-TAS - GNN parameter estimation on full trees (with extinct lineages) in the in-sample data set; T-TES - GNN parameter estimation on extant trees (without extinct lineages) in the in-sample dataset; MLE-TES - MLE parameter estimation on extant trees in the complete dataset; V-TAS - GNN parameter estimation on full trees in the out-of-sample dataset; V-TES - GNN parameter estimation on extant trees in the out-of-sample dataset.

### L. Learning from Summary Statistics

To analyze the relationships between summary statistics and the true parameters used in the diversity-dependent diversification simulations, we computed Pearson correlation indices for each summary statistic against the true values of the three parameters. The absolute values of the correlations are visualized as a heatmap, detailed in Figure 26.

**Fig. 26.**
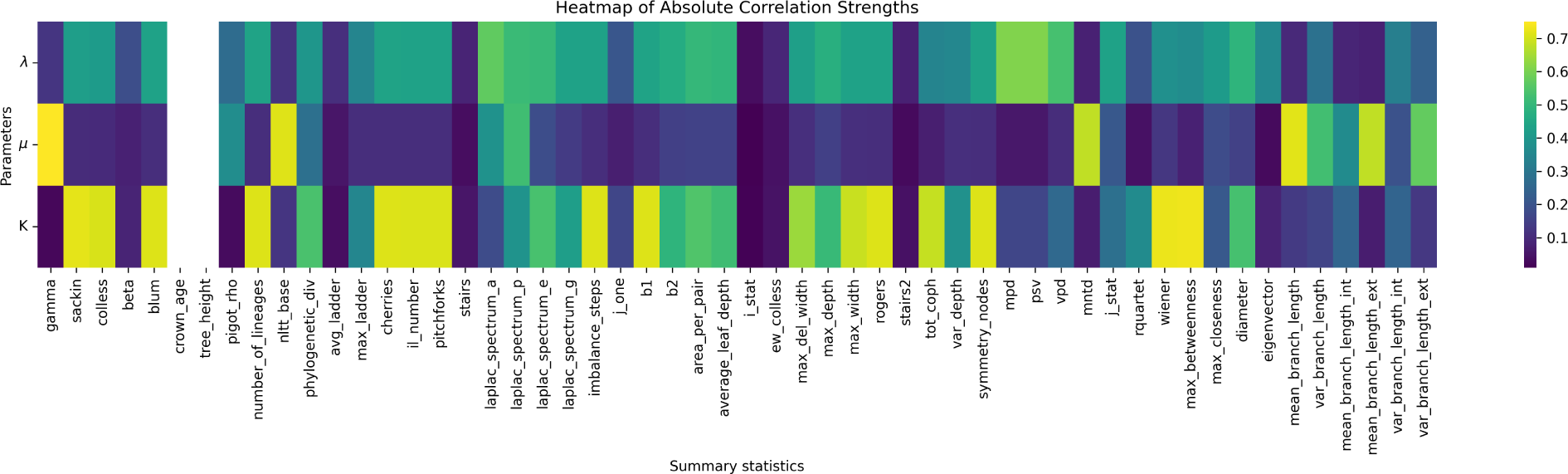
Heatmap of absolute correlation strengths between true parameters and summary statistics from simulated trees under a diversity-dependent diversification scenario. Each column corresponds to a specific summary statistic, while each row corresponds to a true parameter that was used to simulate the phylogenies. The true parameters are denoted as follows: *λ* for speciation rate, *µ* for extinction rate, and *K* for carrying capacity. The color gradient, ranging from dark purple to yellow, represents the increasing values of the absolute Pearson correlations between the summary statistics and the true parameters, See Appendix N for the details of the statistics.

The heatmap analysis reveals that a large number of summary statistics showed a high correlation with carrying capacity, whereas strong correlations with speciation and extinction rates are considerably less frequent. Generally, the correlation strength between the true parameters and extinction rate is markedly lower. These findings align with the observed performance of DNN, which generally yields good estimates for carrying capacity but is much less effective in accurately predicting speciation rates and particularly poor at estimating extinction rates.

We further evaluated the DNN’s performance by benchmarking it against a range of non-neural-network regression techniques using the same summary statistics. These included multivariate linear regression, Ridge regression, Lasso regression, random forest regression, and gradient boosting regression. Our findings indicated that while these traditional regression and machine learning methods generally outperformed the DNN, they still did not match the performance of our more complex neural networks.

Traditional regression methods and other machine learning techniques often outperform linear feed-forward neural networks in classical regression tasks. Such methods can stabilize performance with less data compared to neural networks, which usually require large datasets to generalize effectively. In our study, the dataset size – consisting of 100,000 entries across 54 statistics – may seem substantial, but it is still relatively modest when tasked with regressing multiple parameters simultaneously.

Despite these findings, DNNs have shown efficacy in enhancing the performance of other neural networks, particularly through the prediction of residuals using summary statistics. DNN might not be the optimal choice for estimating parameters from summary statistics alone, but they can be valuable in ensemble learning strategies.

### M. Meta Information of the Selected Empirical Trees

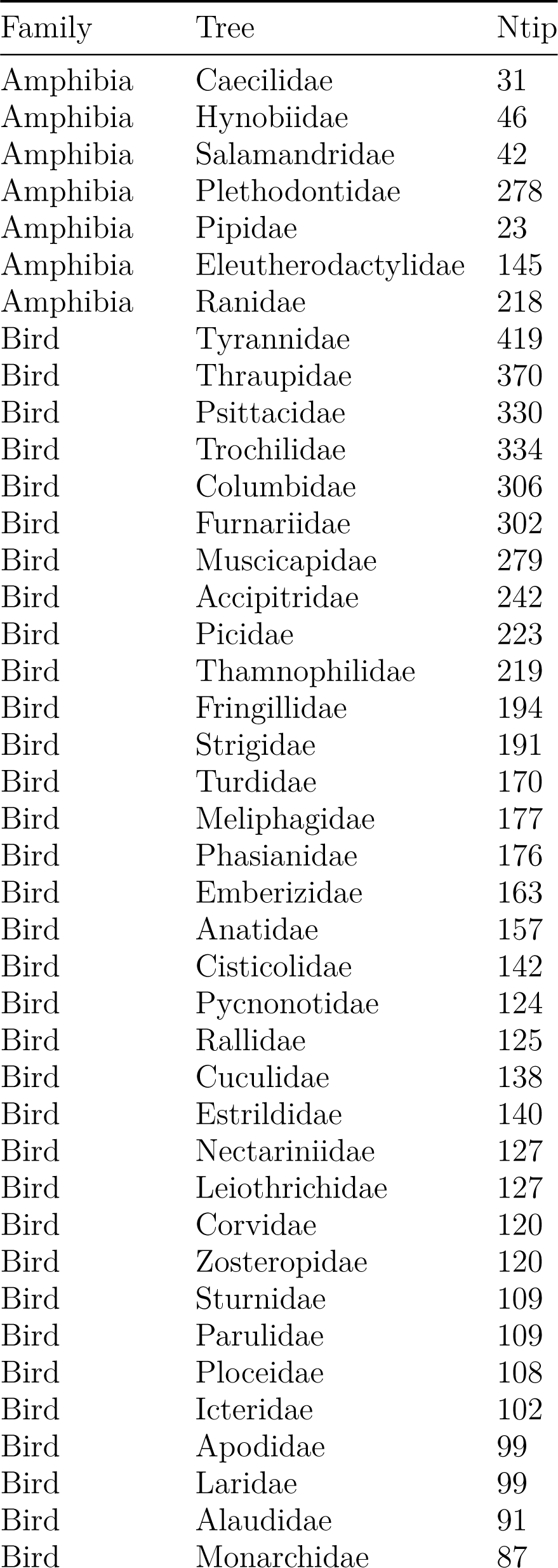
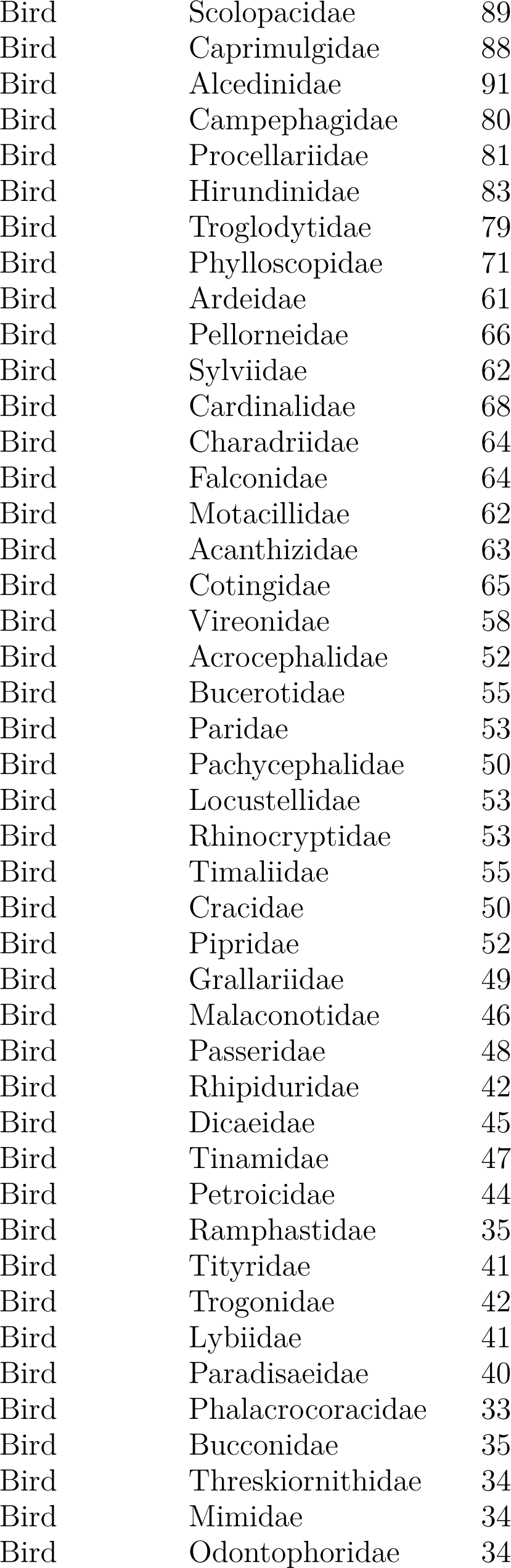
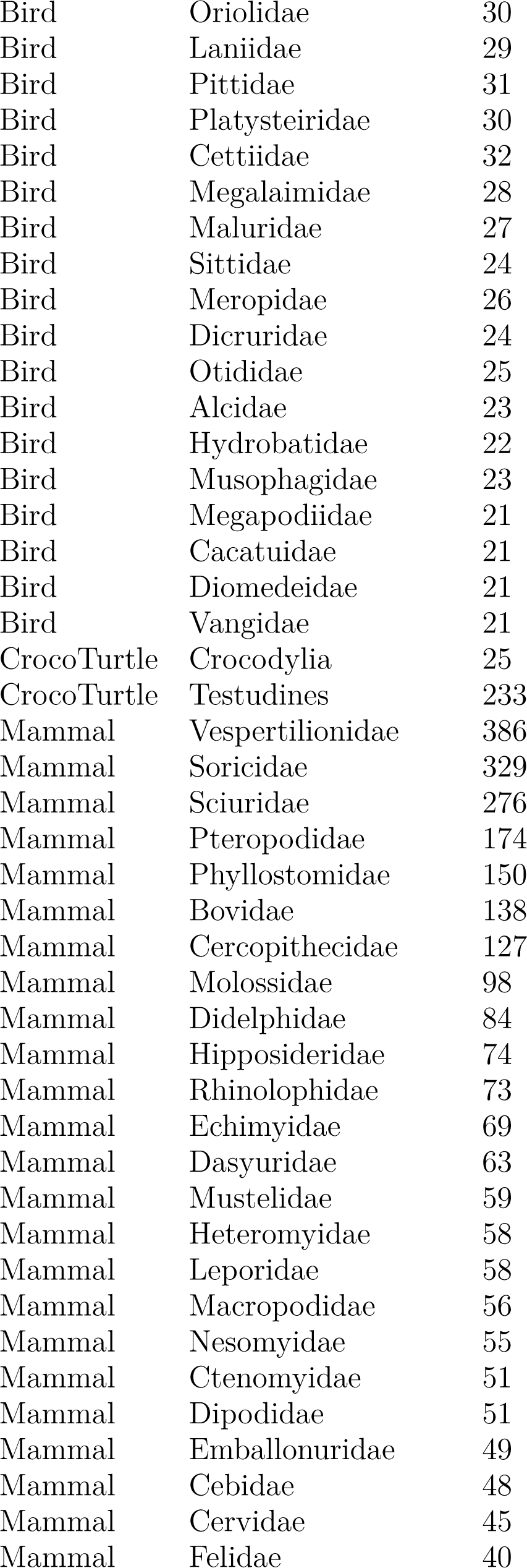
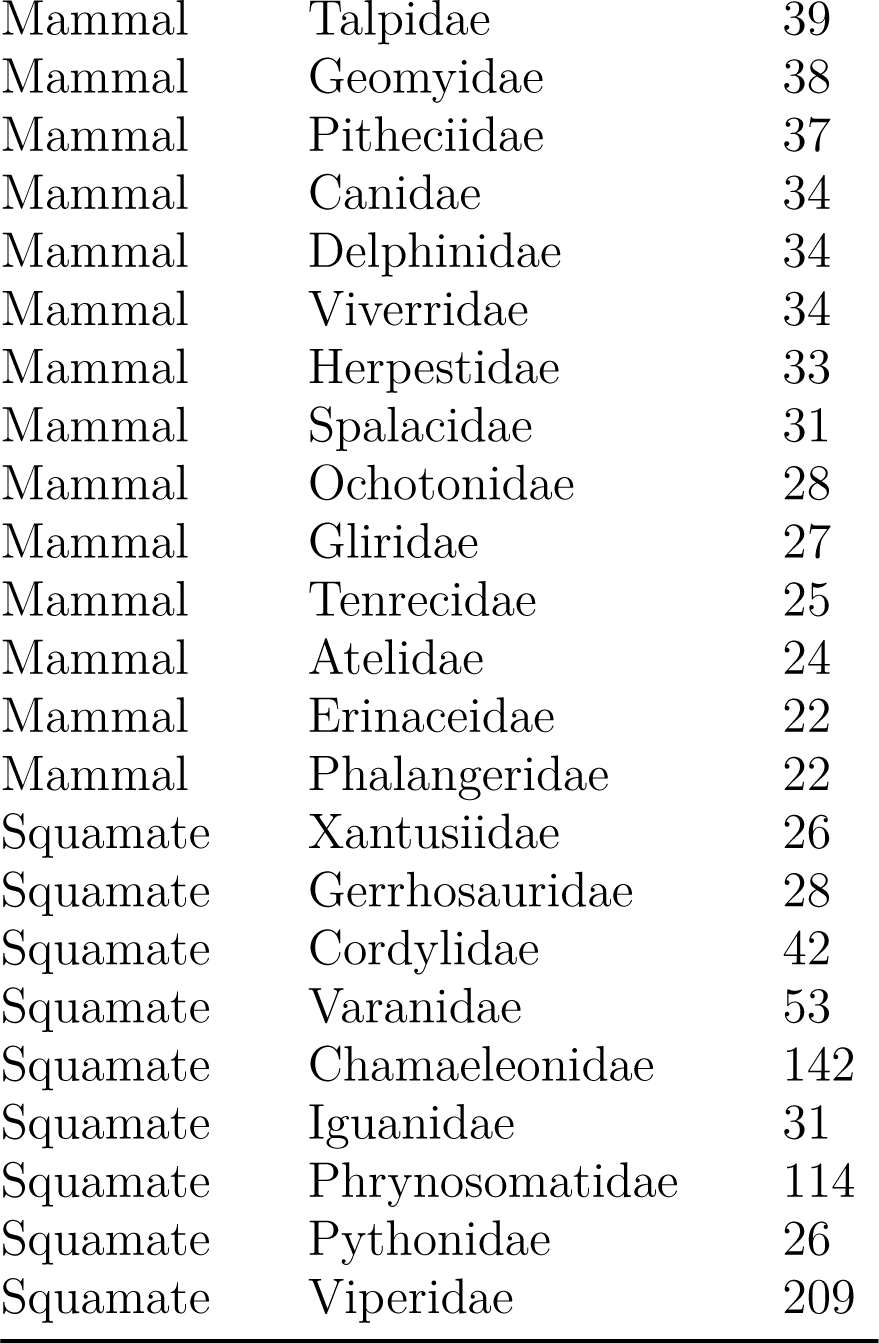

### N. List of Summary Statistics

**Table 5.**
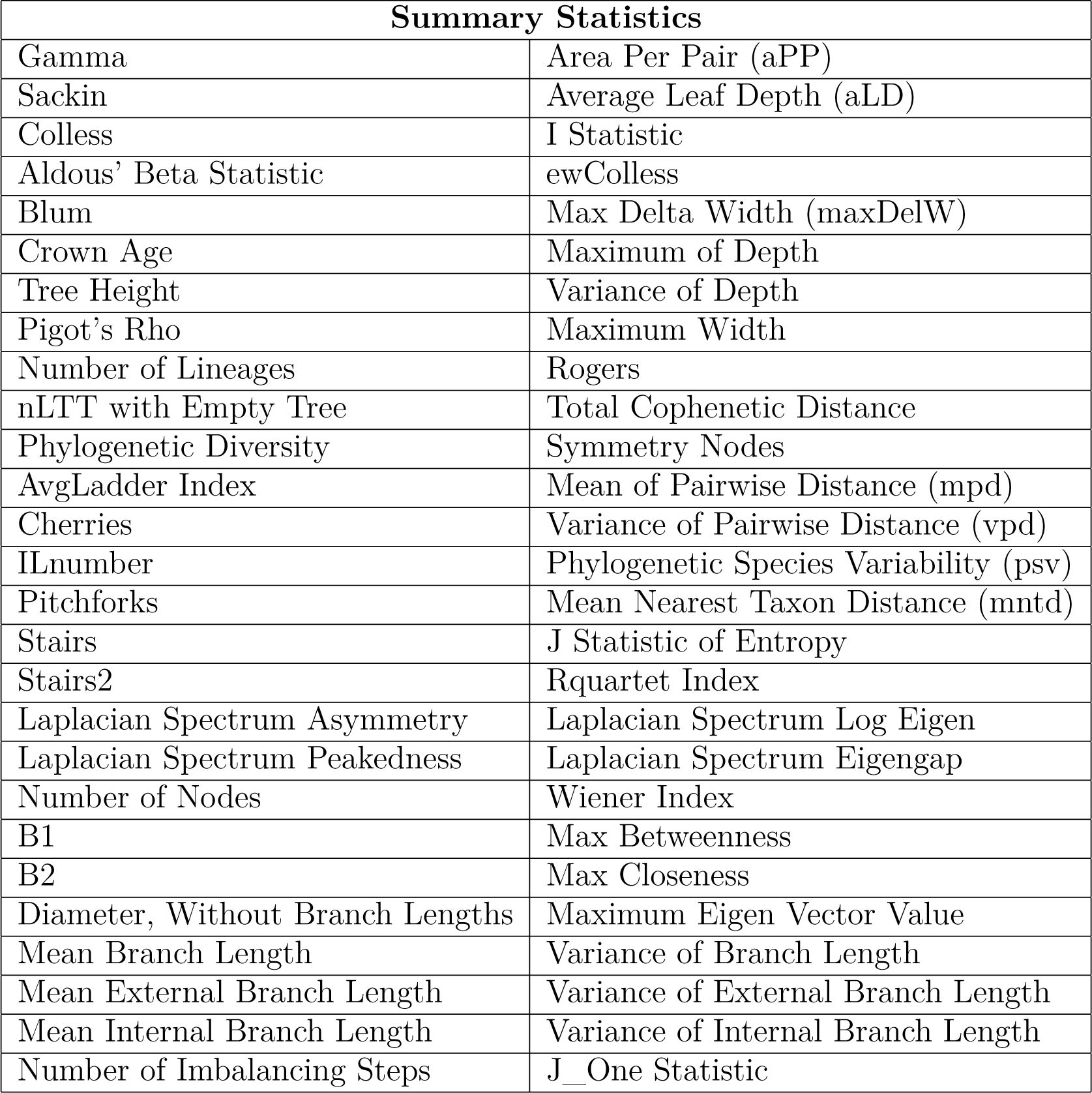
List of phylogenetic summary statistics used in neural network training.

